# Reduced LACTB expression in myeloid cells is associated with elevated succinylcarnitine levels and reduced Alzheimer’s disease risk

**DOI:** 10.64898/2026.03.24.711053

**Authors:** Carmen Romero-Molina, Ruben Gomez-Gutierrez, Wen Yi See, Tulsi Patel, Hayk Davtyan, Jiacheng Ma, Quanyun Xu, Michael Sewell, Kendra Allton, Morgan McReynolds, Olivia Calderon, Yaima L. Lightfoot, Guido Bommer, Carlos Cruchaga, Mathew Blurton-Jones, William J. Ray, Edoardo Marcora, Alison M. Goate

## Abstract

**Background:** Lactamase β (LACTB) is a serine β-lactamase-like mitochondrial enzyme associated with cancer progression, obesity, and lipid metabolism. *LACTB* is located in an Alzheimer’s Disease (AD) risk locus and has been associated with AD in a proteomic study.

**Methods:** We performed Mendelian Randomization (MR) analysis to estimate the association between *LACTB* expression, succinylcarnitine levels, and AD risk. We generated *LACTB* knock-down (KD) THP1 macrophages, *LACTB* knock-out (KO) iPSC-derived microglia and LACTB enzymatically-dead (ED) mice. The impact of LACTB loss-of-function in myeloid cells was characterized via transcriptomics, metabolomics, lipidomics, and functional assays. Finally, human LACTB KO microglia precursors were xenotransplanted into the brains of mice with amyloid pathology to assess *in vivo* interactions with amyloid plaques.

**Results:** MR analyses revealed that lower *LACTB* expression in myeloid cells may lead to reduced AD risk and higher levels of succinylcarnitine, a metabolite associated with AD risk. We identified LACTB as a primary enzyme responsible for succinylcarnitine hydrolysis. Transcriptional and functional studies showed that loss of LACTB enhances OXPHOS, and reduces protein synthesis and triglycerides. *LACTB* expression was upregulated following interferon or TNF stimulation, and its loss modified efferocytosis-related functions under inflammatory conditions. *In vivo*, xenotransplanted human LACTB KO microglia exhibited enhanced association with amyloid plaques.

**Conclusions:** Our findings define a previously unrecognized axis linking LACTB and succinylcarnitine to myeloid cell function and AD susceptibility. Given the druggability of LACTB and the potential for succinylcarnitine to serve as a translational biomarker, this enzyme represents a promising therapeutic target for modulation of neuroinflammation in AD.

## Background

Lactamase β (LACTB) is a β-lactamase–like protein that forms filamentous structures in the mitochondrial intermembrane [1]. Structural and biochemical analyses demonstrated that LACTB contains a conserved catalytic triad characteristic of serine proteases [2] and harbors a D-aspartyl endopeptidase (DAEP) activity that preferentially cleaves at D-Asp residues, which accumulate in long-lived proteins during aging and are present in Alzheimer’s disease (AD)–relevant proteins such as Aβ and tau [3–6].

LACTB has been implicated in controlling cell proliferation and differentiation, with emerging roles in regulating tumor progression and metabolic homeostasis [7]. LACTB has been nominated as an obesity risk gene [8], and more specifically, to be part of the macrophage-enriched metabolic network that might have a causal relationship with traits associated with metabolic syndrome [8–10]. Mice overexpressing LACTB have shown increased fat-mass-to-lean-mass ratio [9], and impaired glucose tolerance and insulin resistance [11].

The macrophage-enriched metabolic network, in which LACTB emerges as a key regulatory node, is significantly enriched within the immune/microglia brain module identified in late-onset AD, a module that also contains the TREM2 signaling adaptor TYROBP as a top causal regulator [12]. *LACTB* is located in a well-established AD risk locus (15q.22) [13–15], often labeled as “APH1B locus” due to the fact that this gene, located in close proximity to *LACTB*, encodes one of the subunits of the ɣ-secretase complex that cleaves APP and thus is highly relevant to AD pathogenesis. Importantly, LACTB has been associated with AD in a proteome-wide association study using brain protein abundance quantitative trait loci (pQTLs) [16] and nominated as an AD target by Agora [17,18]. The human genome contains another β- lactamase–like gene (on chromosome 8) named LACTB2, which has been associated with AD risk by a transcriptome-wide association study integrating myeloid expression quantitative trait loci (eQTLs) with GWAS data [19].

AD, the most common type of dementia, is a neurodegenerative disorder associated with the abnormal accumulation of amyloid plaques and neurofibrillary tangles in the brain, coupled with neurodegeneration, gliosis, lipidosis, and vascular alterations. Genetic studies have consistently implicated myeloid cells of the innate immune system in the etiology of AD [20–22]. Microglia are the resident macrophages of the central nervous system (CNS), responsible for maintaining brain health and responding to injury or infection, mainly through efferocytosis [23,24]. Microglial activation and/or dysfunction are hallmarks of AD and other neurodegenerative disorders. While microglia are the major CNS-resident macrophages, other macrophage populations—including perivascular, meningeal, and choroid plexus macrophages— also contribute to neuroinflammation and disease [25].

LACTB loss-of-function has been associated with increased succinylcarnitine levels both in mice [26] and human [27–29] metabolite GWAS. Importantly, higher succinylcarnitine levels in cerebrospinal fluid (CSF) and brain have been also associated with lower AD risk [30]. Together these data indicate that lower LACTB expression leads to higher succinylcarnitine levels, which in turn may be protective against AD. LACTB has a hydrolase activity [3], but it is still unknown whether its catalytic activity contributes to succinylcarnitine cleavage.

The aim of this work is to better understand the connection between LACTB expression, succinylcarnitine levels and AD risk in the context of myeloid cells.

## Methods

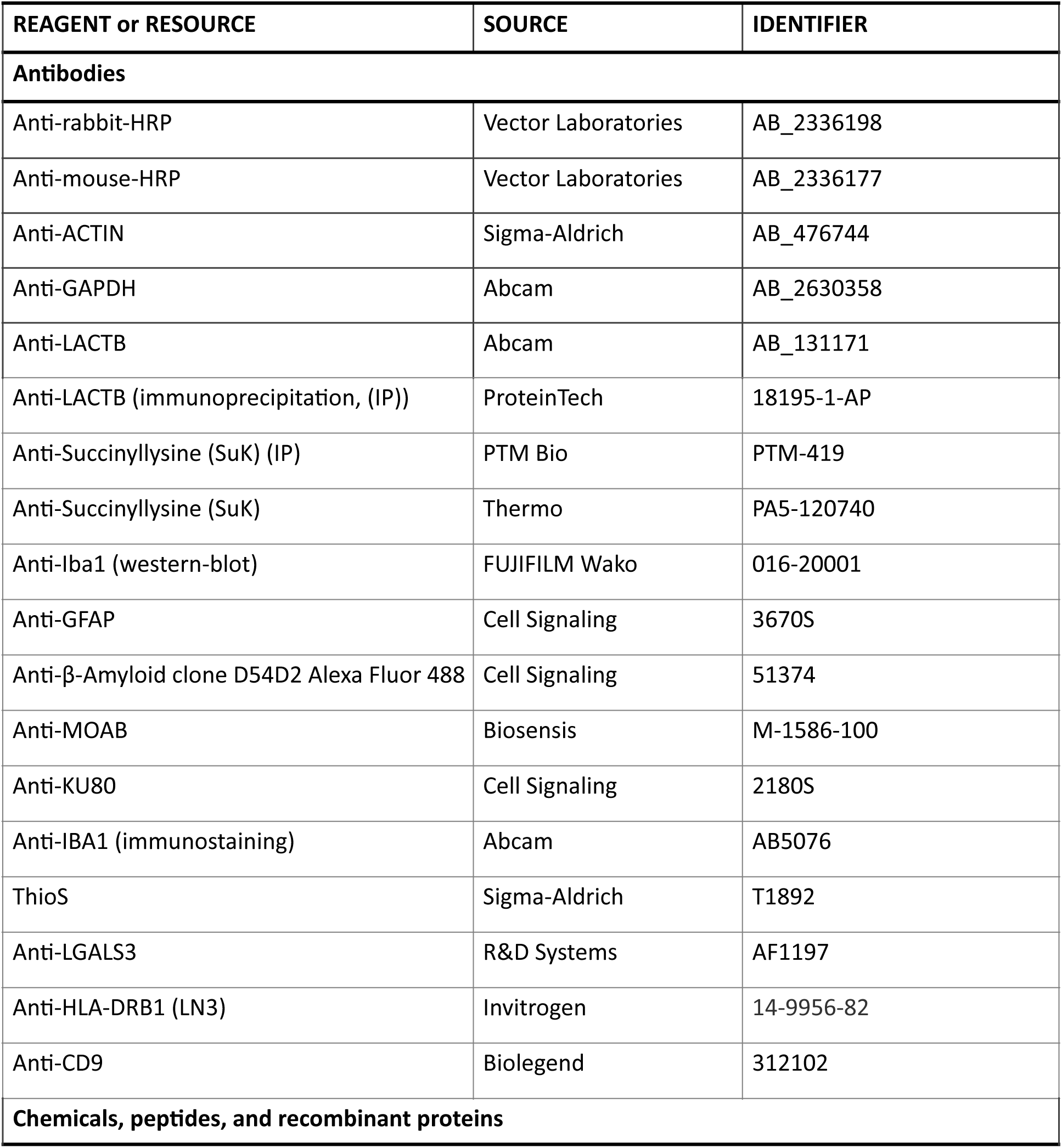
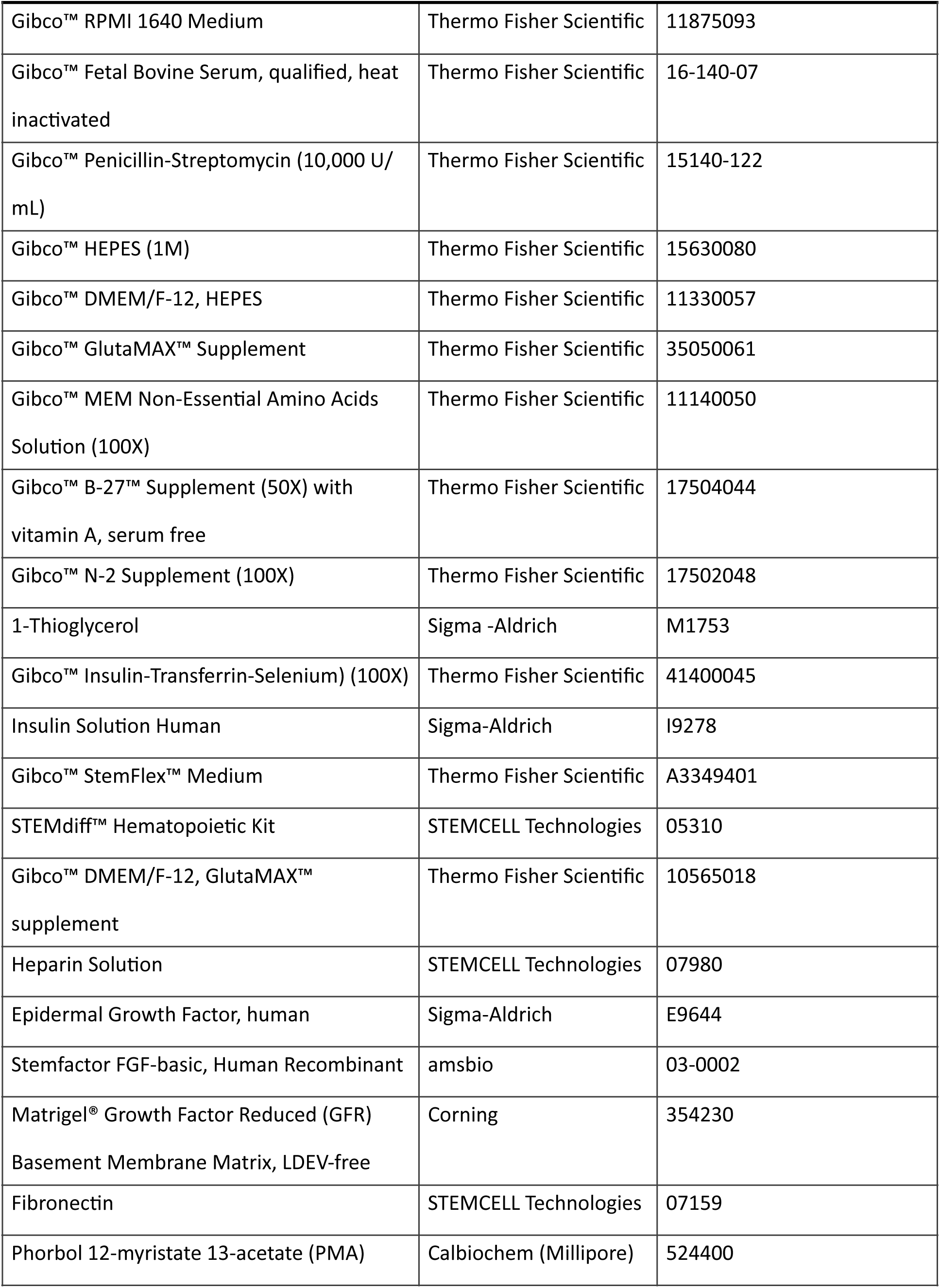
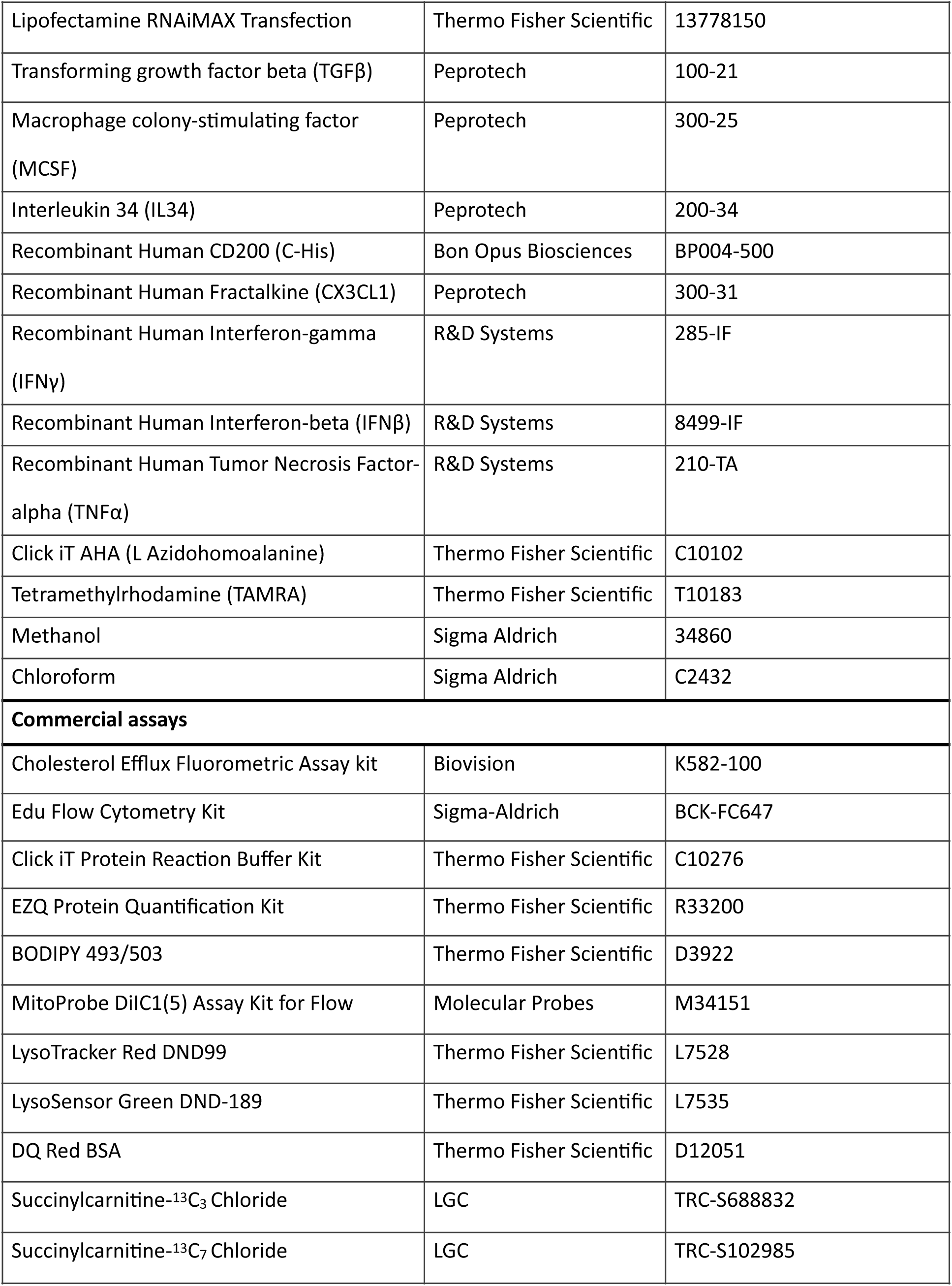
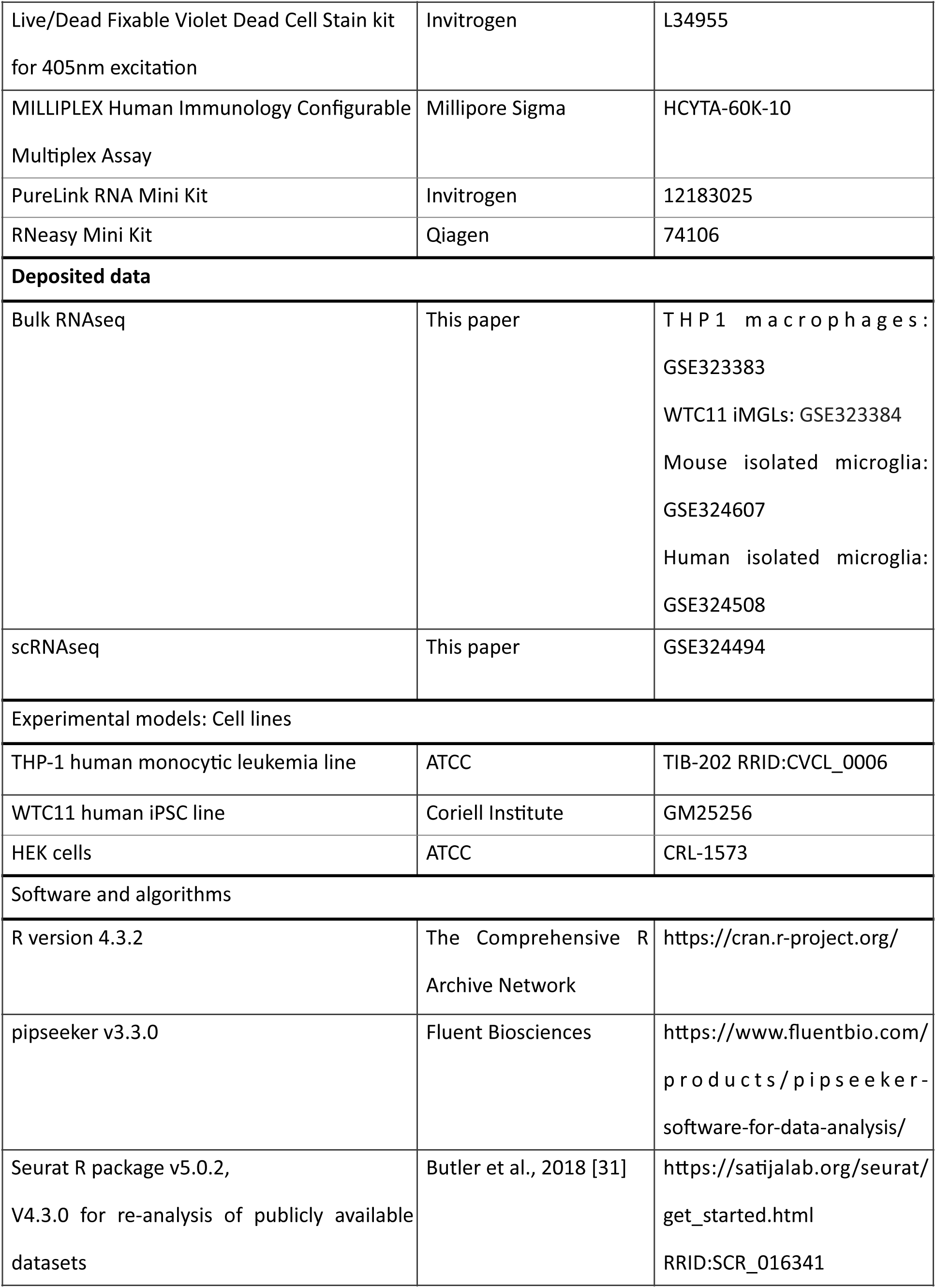
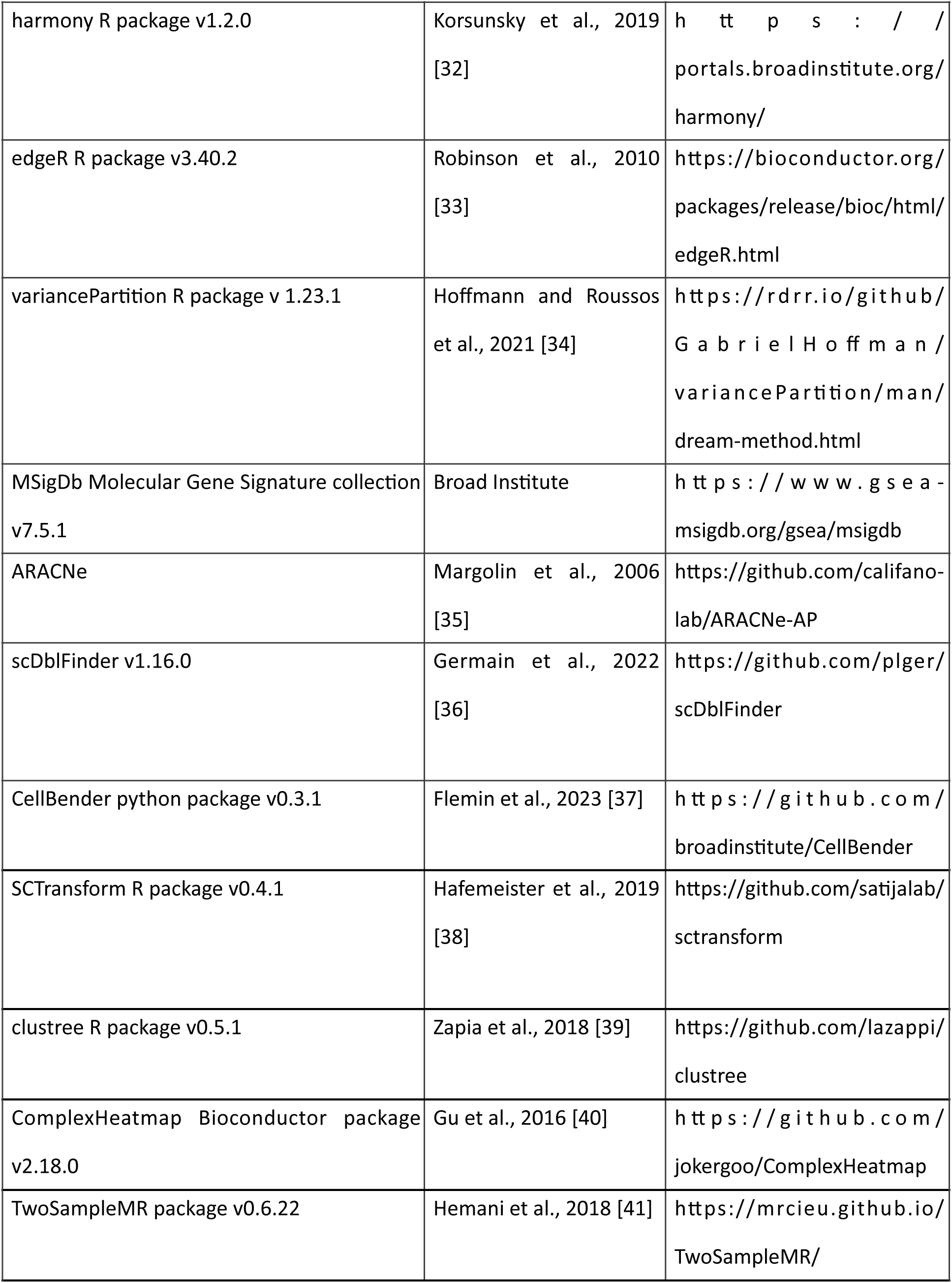
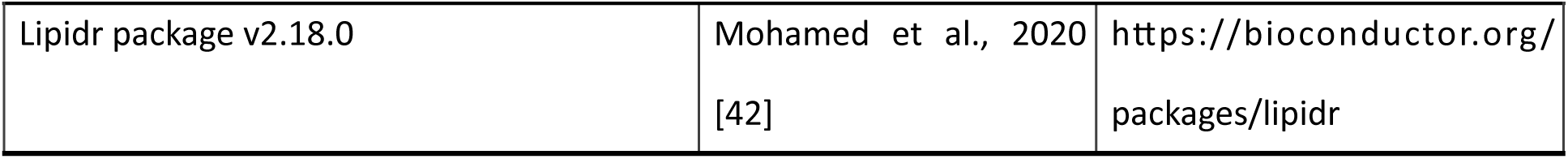

### Mendelian randomization

Two-sample Mendelian randomization (MR) analyses were performed using the TwoSampleMR R package [43]. Summary statistics from cis-eQTL (STARNET, [44]), cis-mQTL [30], and AD GWAS datasets [13] were harmonized to evaluate the causal relationships between LACTB mRNA levels, CSF succinylcarnitine levels, and AD risk. Independent genetic instruments were selected by linkage disequilibrium clumping using a European 1000 Genomes reference panel [45]. Exposure and outcome datasets were harmonized to align effect alleles and remove ambiguous palindromic variants. Causal effects were primarily estimated using the inverse-variance weighted method. Sensitivity analyses included MR-Egger, weighted median, simple mode, and weighted mode approaches. Heterogeneity was assessed using Cochran’s Q statistic, horizontal pleiotropy using the MR-Egger intercept test, and robustness using leave-one-out analyses. All analyses were conducted in R, and the complete reproducible analysis script is available in Additional file 1.

### Mouse models

Wild-type (WT) mice from the C57BL/6 x SJL mixed background (Jackson #034840) were used to assess the levels of LACTB protein and of succinylcarnitine in 2-month-old mice. WT mice (Jackson #006554) were used to assess the brain levels of LACTB protein and of succinylcarnitine in 2-month-old and 12- month-old mice. LACTB ED mice were generated by Ozgene. A targeting vector was constructed for the generation of a conditional knock-in of S162I into exon 3 of the Lactb gene (Additiona file 2) and electroporated into C57BL/6J embryonic stem cells. Transfected embryonic stem cells were selected in the presence of geneticin and screened for correct targeting to the Lactb locus by loss of allele qPCR. Chimeric animals were generated by injection of targeted embryonic stem cell clones into goGermline^TM^ blastocysts and the selection cassette was excised by breeding chimeras to flp expressing females to generate heterozygous conditional knock-in F1 animals. WT and LACTB ED mice were sacrificed at 3 months old to measure LACTB and succinylcarnitine levels.

5XFAD immune-deficient mice (hCSF1) capable of accepting human cells were utilized for xenotransplantation experiments. hCSF1 mice (available as JAX #017708) express human CSF1 on a Rag2/Il2rg deficient background, significantly increasing their ability to engraft and maintain human hematopoietic cells. hCSF1-5XFAD mice overexpress co-integrated APP and PSEN1 transgenes containing five familial AD mutations (the Swedish (K670N/M671L), Florida (I716V), and London (V717I) mutations in APP, and the M146L and L286V mutations in PSEN1) and recapitulate aspects of human AD, including age-dependent cognitive deficits, amyloid accumulation and neuroinflammation [46]. The offspring were backcrossed to hCSF1 mice to return the 3 hCSF1 mutations to homozygosity and maintain the 5XFAD transgene in the hemizygous state [47]. All mice were age-matched and group-housed on a 12-h/12-h light/dark cycle with food and water. Mice were housed with ambient temperature and humidity, and cages and bedding were changed every week. All procedures involving experimentation on animal subjects were done in accordance with the National Research Council’s guide for the care and use of laboratory animals and under an approved protocol.

*Induced pluripotent stem cells (iPSC) donor lines and Hematopoietic Progenitor Cells (HPC) generation*.

The WTC-11 human iPSC line (Coriell Institute for Medical Research, GM25256) was tested for genomic integrity at passage 36 using SNP-array technology (Global Diversity Array v1.0 BeadChip, Illumina). No CNV larger than 1.5 Mb or AOH larger than 3 Mb were detected on autosomal chromosomes. The known WTC-11 deletion of 2.9 Mb on Yp11.2 was detected. We used CRISPR-mediated Non-Homologous End Joining to generate isogenic control (WT) and LACTB KO clonal lines (3 independent clones per genotype), at the Stem Cell Core of the Icahn School of Medicine at Mount Sinai. All clones were validated by next-generation sequencing of both alleles to confirm indels resulting in frameshift mutations or premature stop codons. The use of independent clones is essential to attribute any effect to genotype and rule out that the differences observed are due to random clone to clone variability. Normal karyotype was confirmed after editing. iPSC lines were thawed into matrigel-coated 6-well plates and maintained by feeding every other day with Stemflex supplemented media (Thermo Fisher Scientific) at 37°C, 5% CO_2_. When 70-80% confluence was reached, iPSCs were passaged using ReLeSR dissociation reagent (STEMCELL Technologies, 05872). Hematopoietic stem cells (HPCs) were differentiated from iPSCs using the STEMdiff Hematopoietic kit (STEMCELL Technologies, 05310) according to a previously published protocol [48]. HPCs were collected after a minimum of ten days in culture. If viability was >75%, HPCs were frozen in Bambanker media (Wako Chemicals) or directly differentiated into microglial cells.

*iPSC-derived microglia differentiation*.

Hematopoietic stem cells were differentiated to induced microglial-like cells (iMGLs) using a previously published protocol [48]. iMGLs were maintained and fed with a microglial medium supplemented with three factors (100 ng/ml IL34, 50 ng/ml TGFβ, 25 ng/ml MCSF) for 25 days. On day 25, iMGLs were additionally supplemented with two factors (CX3CL1 and CD200, 100 ng/ml) for an additional three days. Mature iMGLs (day 28) were collected for transcriptomics, lipidomics, metabolomics or RNA/protein quantification or used for functional assays.

### THP1 differentiation and siRNA treatment

We used the monocytic human immortalized cell line THP1 (ATCC, TIB-202 RRID:CVCL_0006), which we treated with phorbol 12-myristate 13-acetate (PMA, 25 ng/ml) for 72 hours to differentiate into macrophages. THP-1 cells were cultured in RPMI medium supplemented with 10% FBS, 1x Penicillin Streptomycin and 10 mM HEPES. We used 40 nM small-interfering RNA (siRNA, ON-TARGETplus SMARTpool, L-021008-01-0005) for 6 days (before and after PMA addition) to reduce LACTB expression. Additional treatments were added during the last 24 hours.

### HEK cells culture and transfection

Stable HEK293 cell lines overexpressing wild-type (WT) or mutant (S164I) LACTB were generated by using the pCDH-CMV-EF1a-puro lentiviral system. HEK cells (ATCC, CRL-1573) were cultured in DMEM supplemented with 10% heat-inactivated FBS and 1% Penicillin/Streptomycin. Cells seeded at 3.5 x 10⁵/ well (6-well plate) were transduced at multiplicities of infection (MOI) of 1, 2, 5, and 10, followed by a 48-hour incubation. Selection for stably integrated clones was performed using puromycin (2 μg/mL) for 4 days, after which viability was assessed, and overexpression was confirmed via Western Blot and qPCR. Following transduction at varied MOIs and 4-day puromycin selection, clones were expanded and cryopreserved, with MOI 1 selected for LACTB WT overexpressing cells and MOI 5 for LACTB S164I/ Vector control overexpressing cells to ensure comparable expression.

### Isolation of microglia and astrocytes from mouse brain

Microglia were isolated from freshly dissected mouse brains using enzymatic dissociation followed by CD11b-based magnetic separation. Brains were sectioned into sagittal slices on ice and dissociated using the Adult Brain Dissociation Kit (Miltenyi Biotec) according to the manufacturer’s instructions with the gentleMACS Octo Dissociator with Heaters. The resulting cell suspension was filtered through a 100 μm cell strainer and centrifuged at 300×g for 10 min at 4°C. Myelin and debris were removed using Debris Removal Solution (Miltenyi Biotec) with density gradient centrifugation (300×g, 10 min, 4°C). The cell pellet was washed and resuspended in MojoSort buffer (BioLegend) and incubated with CD11b MicroBeads (Miltenyi Biotec) for 15 min at 4°C in the dark. Cells were washed and applied to MS columns placed in an OctoMACS separator (Miltenyi Biotec). After washing, magnetically labeled CD11b⁺ cells were eluted, representing the enriched microglial fraction. Isolated microglia were pelleted and stored at −80°C for downstream analyses.

Astrocytes were isolated using magnetic-activated cell sorting (MACS) from the cell pellet remaining after microglia isolation. The pellet was carefully resuspended in 160 µL of cold 1X MojoSort buffer (BioLegend, Cat# 480113) and incubated with 20 µL of FcR Blocking Reagent (Miltenyi Biotec, Cat# 120-097-678) for 10 min at 4°C. Following blocking, cells were magnetically labeled with 20 µL of Anti-ACSA-2 MicroBeads (Miltenyi Biotec, Cat# 130-097-678) for 15 min at 4°C. The cells were then washed with 1 mL of MojoSort buffer, centrifuged at 300xg for 10 min at 4°C, and resuspended in 500 µL of buffer. Magnetic separation was performed using an MS column (Miltenyi Biotec, Cat# 130-042-201) placed in an OctoMACS Separator (Miltenyi Biotec, Cat# 130-042-109). After applying the cell suspension and washing the column three times with 500 µL of buffer, the column was removed from the magnetic field. The positive astrocyte fraction was eluted by flushing the column with 1 mL of MojoSort buffer using a plunger. The resulting suspension was centrifuged at 2500xg for 5 min at 4°C. The pellet was washed with cold 1X PBS before being stored at −80°C.

### Murine bone marrow-derived macrophages (BMDMs) generation

Murine bone marrow cells were harvested from the femurs and tibiae of 12-week-old mice. The marrow was flushed using a 27-gauge needle with ice-cold PBS containing 0.5% BSA and 1 mM EDTA (Ca²⁺/Mg²⁺- free; biotin-free). Cells were dissociated with a syringe plunger and filtered through a 70-µm cell strainer to obtain a single-cell suspension. Red blood cells were lysed with ACK Lysis Buffer for 3 minutes followed by resuspension in 3 volumes of complete media (DMEM with GutaMax supplemented with 10% heat-inactivated FBS and 1% Penicillin/Streptomycin). Progenitor cells (5×10^6^) were seeded into 100mm dishes. Differentiation was induced using complete media containing 50 ng/mL recombinant M-CSF. Cultures were maintained at 37°C, 5% CO2. On day 4, fresh differentiation medium was added to the existing culture. After 7 days of differentiation, mature BMDMs were detached using Trypsin-EDTA (0.05%) and plated at 5 x 10^4^/well density in 96-well flat-bottom plates to measure Suc-D-Asp endopeptidase activity.

### Immunoprecipitation

Immunoprecipitation (IP) was performed using the Pierce™ Crosslink Magnetic IP/Co-IP Kit (Thermo Fisher Scientific, 88805) according to the manufacturer’s protocol, with the following modifications. Briefly, 750 µg of total cell lysate was used per immunoprecipitation. Lysates were incubated with 3.75 µg anti-LACTB antibody (Proteintech,18195-1-AP, rabbit IgG), anti-SuK antibody (PTM Biolabs, Cat. No. PTM-419, mouse IgG), or the corresponding control IgG (rabbit IgG, I-1000; mouse IgG, I-2000). Antibody–antigen complexes were captured using 30 µL of magnetic Protein A/G beads and incubated for 1.5 h with gentle rotation at 4°C. Beads were washed according to the kit protocol using IP lysis buffer, followed by a final wash with ultrapure water. Bound proteins were eluted in 35 µL of elution buffer, and eluates were immediately neutralized with the provided neutralization buffer. The entire eluted immunoprecipitated material was resolved by SDS-PAGE and transferred to membranes. Membranes were immunoblotted with anti-LACTB antibody (Abcam, ab131171) or anti-SuK antibody (Thermo Fisher Scientific, PA5-120740), as indicated. Blots were developed using Clean-Blot™ HRP- conjugated secondary antibody according to the manufacturer’s instructions.

### Succinyl-D-asp assay

To measure LACTB cleavage of succinyl-D-Asp substrate, cells were washed with DPBS and incubated with an assay buffer consisting of 10 mM Tris-HCl (pH 8.5), 200 mM NaCl, and 3 mM MnCl_2_ (for WTC11 microglia and THP1 macrophages) or in PBS (for BMDMs and HEK cells). The fluorogenic substrate succinyl-D-Asp (7-methoxycoumarin-4-acetyl-L-succinyl-D-aspartyl-L-lysine, #3222-v, Peptide) was added to each well at a final concentration of 5 or 20 μM. Kinetic fluorescence measurements (λex = 380 nm, λem = 460 nm) were performed immediately using a Varioskan or a BioTek Synergy Neo2 plate reader at 37°C, with 30-60 readings taken at 2-min intervals with shaking before each read (300 rpm). Following the kinetic run, cells were washed with PBS and lysed in RIPA buffer supplemented with protease and phosphatase inhibitors for 30 minutes on ice. Total protein concentration was determined via BCA assay, and fluorescence slopes were normalized to total protein content to account for variation in cell density.

*<u>Succinylcarnitine recovery assay</u>*.

In-house generated recombinant human LACTB (IACSp-1689-07c) was used to determine kinetic parameters for succinylcarnitine cleavage. Reactions were performed in an assay buffer containing 20 mM Tris-HCl (pH 8.5) and 1 mM TCEP (Tris(2-carboxyethyl)phosphine). LACTB was used at a final concentration of 100 nM. Where indicated, the inhibitor Ac-IEPD-CHO (10 µM final concentration, Abcam, ab142030) or vehicle (DMSO) was pre-incubated with the enzyme for 15 min prior to substrate addition. Ac-IEPD-CHO inhibitor is similar to Z-AAD-CMK inhibitor [2]; both are Granzyme B (a serine protease similar to LACTB) inhibitors with the difference that the aldehyde group (-CHO) makes the IEPD- CHO an reversible inhibitor compared to the -CMK being irreversible. Reactions (200 µL total volume) were initiated by adding ^13^C_7_-succinylcarnitine chloride (TRC-S102985) at varying concentrations and incubated for 2 h at room temperature with shaking (300 rpm). Reactions were quenched by addition of 800 µL of 80% methanol, followed by incubation at −80°C for 24 h to precipitate protein. Samples were centrifuged at 21,000×g for 15 min, and supernatants were collected for metabolite quantification. Substrate (^13^C_7_-succinylcarnitine chloride) and product (^13^C_3_-carnitine) were measured by targeted mass spectrometry.

### Western-blotting

Cells were lysed in RIPA buffer (Thermo Scientific, 89900) supplemented with Protease/Phosphatase Inhibitor Cocktail (Cell Signaling, 5872) following manufacturer’s instructions. Protein concentration was measured using BCA kit (Thermo Fisher Scientific, 23225) and equal quantities were used to prepare samples for western blotting. Samples were resolved by electrophoresis with Bolt 4–12% Bis-Tris Plus Gels (Invitrogen) in Bolt MES SDS running buffer (Invitrogen, B0002) and transferred using iBlot 2 nitrocellulose transfer stacks (Invitrogen). Membranes were blocked for 1 h and probed with primary antibodies in 5% BSA in PBS/0.1% Tween-20 buffer overnight at 4°C. Secondary antibody at 1:10,000 dilution was applied for 1 h at RT, visualized using WesternBright ECL HRP Substrate Kit (Advansta, K-12045), and measured using iBright imaging system (Applied Bioscience). Images were analyzed using ImageJ (NIH). Uncropped western blot images are provided in Additional file 3.

### Metabolomic assays

For THP1 macrophage collection (700,000 cells per sample), media was aspirated and cells were quickly washed with ice-cold 0.85% (w/v) ammonium bicarbonate in water. 1mL of buffer was added per well, and cells were gently detached using a cell scraper. Cell suspensions were transferred to tubes and centrifuged at 500 × g for 3 min. Supernatants were aspirated, and pellets were snap-frozen on dry ice and stored at −80°C.

For the collection of semi-adherent WTC-11 iMGLs (500,000 cells per sample), media was collected, and 1 mL of ice-cold DPBS was added to each well. Cells were gently detached using a cell scraper and pooled with their respective media. Samples were centrifuged at 300×g for 5 minutes, and supernatants were aspirated. Pellets were resuspended in 1 mL of ice-cold DPBS, centrifuged again at 300×g for 5 min. The final supernatants were aspirated, and the pellets were snap-frozen on dry ice and stored at −80°C.

For ion chromatography-mass spectrometry (ICMS) analysis, metabolites were extracted using ice-cold 50/30/20 (v/v/v) methanol/acetonitrile/water. Extracts were left on ice for 10 mins, centrifuged at 17,000 *g* for 5 min at 4°C, and supernatants were transferred to autosampler vials, and 10 μL was injected. IC mobile phase A (MPA; weak) was water, and mobile phase B (MPB; strong) was water containing 100 mM KOH. A Thermo Scientific Dionex ICS-6000+ system included a Thermo IonPac AS11 column (4 µm particle size, 250 x 2 mm) with column compartment kept at 35°C. The autosampler tray was chilled to 4°C. The mobile phase flow rate was 360 µL/min, and the gradient elution program was: 0-2 min, 1% MPB; 2-25 min, 1-40% MPB; 25-39 min, 40-100% MPB; 39-50 min, 100% MPB; 50-50.5, 100-1% MPB. The total run time was 55 min. To assist the desolvation for better sensitivity, methanol was delivered by an external pump and combined with the eluent via a low dead volume mixing tee. Data were acquired using a Thermo Orbitrap IQ-X Tribrid Mass Spectrometer under ESI negative ionization mode at a resolution of 240,000.

For Hydrophilic Interaction Liquid Chromatography analysis, the same samples were injected by liquid chromatography (LC)-MS. LC mobile phase (MP) A was 95/5 (v/v) water/acetonitrile containing 20 mM ammonium acetate and 20 mM ammonium hydroxide (pH∼9), and MPB was acetonitrile. Thermo Vanquish LC system included a Xbridge BEH Amide column (3.5 µm particle size, 100 x 4.6 mm) with column compartment kept at 30°C. The autosampler tray was chilled to 4°C. The mobile phase flow rate was 350 µL/min, and the gradient elution program was: 0-1 min, 85% MPB; 1-16 min, 85-5% MPB; 16-20 min, 5% MPB; 20-21 min, 5-85% MPB; 21-25 min, 85% MPB. The total run time was 25 min and data were acquired using a Thermo Orbitrap Exploris 240 Mass Spectrometer under ESI positive/negative ionization (polarity switching) mode at a resolution of 240,000.

^13^C_7_-succinylcarnitine and ^13^C_3_-carnitine were quantitatively analyzed by using an LC-MS/MS method. Acetyl d_3_-carnitine was used as an internal standard. An Agilent Technologies 1290 Infinite liquid chromatograph consisting of a binary pump, a multisampler with a 10-µL loop, and a column heater was used. The stationary phase was a Waters X-Bridge amide column (100 x 4.6 mm, 3.5 µm particle size). The column temperature was set at 40°C. MPA was 5% acetonitrile in water containing 20 mM ammonium acetate and 20 mM ammonium hydroxide and MPB acetonitrile. The MP was delivered at 0.3 mL/min in a gradient mode at 85% B from 0 to 3 minutes, linearly decreased to 30% B at 10 min, then to 2% B at 20 min, and equilibrated at 85% B from 20.2 to 25 min. The injection volume was 5 µL. Under these conditions, ^13^C_7_-succinylcarnitine, ^13^C_3_-carnitine, and acetyl d_3_-carnitine had retention times of 11.1, 11.0, 11.6, and 10.9 minutes, respectively. An Agilent Technologies 6460 Triple Quad mass spectrometer was operated in the positive electrospray ionization mode. The following operating conditions were set: desolvation gas temperature 300°C; nebulizer gas flow 11 L/min, nebulizer gas pressure 35 psi; shealth gas flow 11 L/min, shealth gas temperature 400°C; capillary voltage 4 kV. ^13^C_7_- succinyl L-carnitine, ^13^C_3_-carnitine, and acetyl d_3_-carnitine were monitored in MRM mode at mass transitions: 262.1 > 85.1, 270.2 > 86.0, 165.2 > 63.1, and 207.1 > 85.1, respectively. Stock solutions of ^13^C_7_-succinyl L-carnitine, ^13^C_3_-carnitine were prepared in water and its working solution was prepared by diluting the stock solution with a mixture of methanol/water (80: 20, v/v). Calibration curves were constructed in the concentration ranges from 0.078 to 20 ng/mL or 0.78 to 200 ng/mL. An aliquot of 50 µL of samples, standards, and QC samples was diluted with 150 µL of the internal standard solution containing acetyl d_3_-carnitine at 5 ng/mL in acetonitrile before the analysis by LC-MS/MS method.

All the raw files were imported to Skyline Daily for final analysis. For the labeled metabolites analysis, the fractional abundance of each isotopologue is calculated by the peak area of the corresponding isotopology normalized by the sum of all isotopology areas.

### Bulk RNAseq and qPCR analyses

RNA from human THP-1 macrophages and WTC11 iMGLs was extracted using the RNeasy Plus Mini kit (Qiagen, 74106) following manufacturer’s instructions. RNA was extracted from microglia isolated from LACTB ED mice using PureLink RNA Mini Kit (Invitrogen, 12183025) following the manufacturer’s instructions. mRNA quantity was measured using Nanodrop 8000 (Thermo Fisher Scientific). RNA was extracted from human xenotransplanted microglia using the RNeasy FFPE kit (Cat# 73504, Qiagen) according to the manufacturer’s protocol.

For both bulk and single-cell RNAseq, RNA was submitted to Azenta (New Jersey, NJ, USA) for QC, library preparation, and next-generation sequencing. Samples passed quality control with Qubit and BioAnalyzer showing RIN > 9.0. Libraries were prepared using TruSeq RNA Sample Prep Kit v2 and paired-end sequenced using HiSeq2500 at a read length of 150bp to obtain 20-30M mapped fragments per sample. Sequenced reads were assessed for quality (FastQC v0.11.8), trimmed for adapter contamination (Cutadapt v2.6), and aligned to the human genome hg38 (STAR v2.5.3a). Differential gene expression analysis (DGEA) was performed using a linear mixed model implemented in DREAM (Differential expression for repeated measures, variancePartition R package [49]). To identify pathways, we used Gene Set Enrichment Analysis (GSEA) [50]. Briefly, ranked lists were generated from differential gene expression analyses by ordering genes according to the signed Z statistic (Table 2). Our preranked lists were tested for enrichment against genesets from the Molecular Signatures Database (MSigDB v7.5.1, Broad Institute). Pathways displayed were selected among the most significant pathways, the full list is provided in Table 2. Enrichment scores were normalized by geneset size to generate normalized enrichment scores (NES) according to the standard protocol [50].

**Table 2.** Bulk RNAseq results. List of DEGs and GSEA analyses for the bulk RNAseq data in the manuscript in THP1 macrophages, WTC11 iMGLs *in vitro,* mouse microglia and WTC11 iMGLs *ex vivo*.

For qPCR analysis, reverse transcription reaction was performed with 1000 ng of RNA using High-Capacity RNA-to-cDNA kit (Thermo Fisher Scientific, cat. 4387406). 10 ng cDNA was used in the qPCR reaction with Power SYBR Green Master Mix (Applied Biosystems, cat. 4368706) run using QuantStudio 7 Flex Real-Time PCR System (Thermo Fisher Scientific). Primers were designed using the IDT software and are listed in Table 5. Ct values were averaged from two technical replicates for each gene, GAPDH Ct values were used for normalization. Gene expression levels were quantified using the 2-ddCt method.

**Table 5.** Primers for qPCR. Sequences of the primers used for qPCR.

### Analysis of publicly available data

Fastq files from publicly available single-cell RNA sequencing datasets were downloaded from the Gene Expression Omnibus for Mancuso et al. (GSE216999), Tuddenham et al. (GSE204702), and Smith et al. (GSE160936), and from the Synapse platform for Olah et al. (syn21438358), Prater et al. (syn51272688, accessed via data use agreement), and Lee et al. (syn52795287). Raw sequencing data were processed using Cell Ranger (v9.0, 10x Genomics). Doublet detection and removal were performed using scDblFinder (v1.16.0). Ambient RNA correction and batch integration were conducted using CellBender (v3.0.1) and Harmony (v1.2.0), respectively, where appropriate. All downstream analyses were performed using Seurat (v4.3.0), with normalization carried out using either the NormalizeData() function or in conjunction with SCTransform (v0.4.1). Clustering for each dataset was determined through differential expression analysis in Seurat and resolution assessment using the clustree package (v0.5.1). Pearson correlation analysis with the cor() function in R was used to assess correlations across clusters between aggregated expression for *LACTB* and leading-edge genes contributing to significantly enriched terms identified by GSEA in LACTB THP1 knockdown and iMGL knockout datasets. Correlation data were then visualized using the ComplexHeatmap() package.

### Single-cell RNAseq analyses

Paired-end FASTQ files were processed using Pipseeker v3.3.0 following the recommended pipeline. Reads were trimmed, aligned to the GRCh38 reference genome, and quantified to generate feature-barcode count matrices at multiple sensitivity levels; sensitivity level 3 was selected for downstream analyses. Count matrices were imported into Seurat v5.0.2 for quality control and analysis. Cells expressing fewer than 100 genes, less than 500 UMIs and greater than 95% of total UMIs, or with >10% reads mapped to the mitochondrial genome were excluded. Doublets were identified and removed using scDblFinder v1.20.2. Raw counts were normalized using SCTransform v0.4.1, with regression of mitochondrial mapping percentage. Principal component analysis was performed for dimensionality reduction, and samples were integrated using Harmony v1.2.0 to account for batch effects. Unsupervised clustering was conducted in Seurat and clusters were visualized using UMAP. Marker genes were identified using FindMarkers. Cluster annotation was based on (i) expression of established marker genes from the literature, (ii) enrichment of published myeloid gene signatures calculated with hypergeometric overlap, and (iii) biological pathway enrichment analysis using GSEA analysis to determine functional relevance.

For pseudobulk differential gene expression analysis (DGEA) between genotypes, cells were aggregated per sample to obtain average expression for each gene (N=3 independent samples per condition) using Seurat AggregateExpression. Pseudobulked DGEA was performed using the DREAM package in R [49], with adjustment for the experimental batch to account for technical variability. Geneset enrichment across clusters was calculated using the Seurat AddModuleScore function, which computes a per-cell module score to assess relative enrichment of gene signatures. Differential cell-type abundance between genotypes was assessed per cluster using Crumblr (under review [51]). Pseudobulk cell-type proportions were calculated per sample and differential abundance testing was performed using generalized linear modeling while accounting for batch.

### LACTB co-expression module generation

A normalized scRNA-seq gene-by-cell matrix was used to compute pairwise Pearson correlations, which were used to construct k-nearest neighbor graphs. Metacells were generated by aggregating counts from neighboring cells using the PISCES MetaCell framework (unpublished [52]) prior to construction of the LACTB co-expression module using ARACNe [35]. Genes with non-zero counts in at least 75% of metacells were retained for downstream analysis. ARACNe was applied with default parameters to reconstruct a gene co-expression module centered on LACTB, using a mutual information threshold of p=1×10^−8^ and 100 bootstraps. Only significant interactions (FDR-adjusted P < 0.05) were retained to define the LACTB co-expression module. Enrichment of genes co-expressed with LACTB across clusters was assessed using Seurat AddModuleScore, and enrichment in external published datasets was evaluated by hypergeometric overlap with cluster marker genes.

### Proliferation assays

We used the Edu Flow Cytometry Kit 647 (Sigma-Aldrich, BCK-FC647-50). Cells were stained with EdU at 10 μM for 2 hours and manufacturer’s instructions were followed. Briefly, detached cells were collected and washed with 1% BSA in DPBS. Cells were centrifuged 300xg 5 min and resuspended in the Fixative solution. Cells were washed again with 1% BSA in DPBS, centrifuged 300xg 5min and pellets were resuspended in saponin 1X buffer (20 min RT). The assay cocktail mix was added for 30 min RT. Cells were centrifuged 300xg 5 min and washed with saponin 1X buffer. Cells were centrifuged 300g 5min and resuspended in saponin 1X buffer for flow analysis (using a low-rate flow, as indicated in the manufacturer’s instructions). Cells were gated based on forward scatter and singlet discrimination. Non– EdU–treated cells processed in parallel were used to define the negative gate, accounting for the background signal generated by the Click-iT reaction.

### Seahorse assays

THP-1 monocytes were plated at a density of 50,000 cells per well in a Seahorse 96 well-plate; and WTC11 iMGLs at a density of 100,000 cells per well in a fibronectin-coated Seahorse 96 well-plate. Agilent Seahorse Mito Stress Test Kit (103015-100) was used to assess mitochondrial respiration. DMEM XF Base Media was supplemented with 1 mM pyruvate, 2 mM glutamine and 10 mM glucose. Cell media was replaced with 180 ul of this media and cells were incubated for 1 h at 37°C in a non-CO_2_ incubator. Cells were stimulated with 1 μM oligomycin, 0.5 μM FCCP, 0.5 μM Rot/AA.

### Protein synthesis assay

THP-1 monocytes and WTC-11 iMGLs were plated in 6-well plates at densities of 700,000 and 500,000 cells per well, respectively. On the day of collection for THP-1 macrophages, culture media was aspirated, and cells were washed once with methionine-free THP-1 media (methionine-free RPMI supplemented with 10% FBS, 1% Penicillin/Streptomycin, and 1% HEPES). Wells were replenished with 1 mL of methionine-free media and incubated for 1 hour to deplete endogenous methionine. For iMGLs, methionine-free incubation was omitted. Cells were then treated with 50 μM of Click-iT AHA (L-Azidohomoalanine), a methionine analog used for nascent protein labeling. Thapsigargin, a known inducer of the integrated stress response that suppresses protein translation, was used as a positive control at a final concentration of 250 nM for THP1 macrophages and 500 nM for iMGLs. Cells were treated for 2 h (THP1) or 3 h (iMGLs).

Following treatment, cells were collected, washed in DPBS, and lysed in RIPA buffer supplemented with protease and phosphatase inhibitors Cell Signaling, 5872S). Lysates were incubated on ice for 30 minutes with vortexing every 10 min, followed by sonication. Samples were then centrifuged at 13,000 rpm for 12 min, and the supernatants were collected as protein lysates. Click-iT protein labeling reactions and subsequent protein quantification using the EZQ Protein Quantification Kit were performed according to the manufacturer’s instructions.

### Lipidomic profiling and analysis

For THP1 macrophages, we used 3 million cells per sample, pooling 3 independent siRNA-treated wells for each. We performed 4 independent macrophage differentiations. For iMGLs, we used 2 million cells per sample, pooling cells from 6 wells of a 6-well plate.

Cell pellets were shipped to Columbia Biomarker Core (NY, US, [53]), where a standard lipid panel was performed. Lipidomics profiling was performed using Ultra Performance Liquid Chromatography-Tandem Mass Spectrometry [54]. Lipid extracts were prepared from cell lysates spiked with appropriate internal standards using a modified Bligh and Dyer method and analyzed on a platform comprising Agilent 1260 Infinity HPLC integrated to Agilent 6490A QQQ mass spectrometer controlled by Masshunter v 7.0 (Agilent Tecmethod and Santa Clara, CA). Glycerophospholipids and sphingolipids were separated with normal-phase HPLC as described before [55], with a few modifications. An Agilent Zorbax Rx-Sil column (2.1 x 100 mm, 1.8 µm) maintained at 25°C was used under the following conditions: mobile phase A (chloroform: methanol: ammonium hydroxide, 89.9:10:0.1, v/v) and mobile phase B (chloroform: methanol: water: ammonium hydroxide, 55:39:5.9:0.1, v/v); 95% A for 2 min, decreased linearly to 30% A over 18 min and further decreased to 25% A over 3 min, before returning to 95% over 2 min and held for 6 min. Separation of sterols and glycerolipids was carried out on a reverse phase Agilent Zorbax Eclipse XDB-C18 column (4.6 x 100 mm, 3.5 µm) using an isocratic mobile phase, chloroform, methanol, 0.1 M ammonium acetate (25:25:1) at a flow rate of 300 μl/min. Quantification of lipid species was accomplished using multiple reaction monitoring transitions [55] under both positive and negative ionization modes in conjunction with referencing of appropriate internal standards: PA 14:0/14:0, PC 14:0/14:0, PE 14:0/14:0, PG 15:0/15:0, PI 17:0/20:4, PS 14:0/14:0, BMP 14:0/14:0, APG 14:0/14:0, LPC 17:0, LPE 14:0, LPI 13:0, Cer d18:1/17:0, SM d18:1/12:0, dhSM d18:0/12:0, GalCer d18:1/12:0, GluCer d18:1/12:0, LacCer d18:1/12:0, D7-cholesterol, CE 17:0, MG 17:0, 4ME 16:0 diether DG, D5-TG 16:0/18:0/16:0 (Avanti Polar Lipids, Alabaster, AL). Lipid levels for each sample were calculated by summing the total number of moles of all lipid species measured by all three LC-MS methodologies and then normalizing that total to mol %. Results were analyzed using the lipidr package v2.18.0 [42].

### Electron microscopy

WTC11 iMGLs (2 clones per genotype) were seeded at 700,000 cells per well in a 6-well plate, combining 6 wells for each collection. After collection, cells were fixed with 2.5% glutaraldehyde in 100mM cacodylate buffer pH 7.4 + 60 mM sucrose and stored at 4°C. Samples were analyzed at Mayo Microscopy and Cell Analysis Core (Rochester, MN, US, [56]). Briefly, samples were rinsed three times for 5 min with a rinse buffer (100 mM cacodylate, 200 mM sucrose) at RT. Secondary fixation was performed for 60 min using 50 mM cacodylate, 100 mM sucrose, and 1% OsO_4_, followed by 3 washes in dH_2_O. A tertiary fixation step was carried out in 1% uranyl acetate (UAc) in dH_2_O for 60 min at RT or overnight at 4°C. Dehydration was performed through a graded ethanol series (25%, 50%, 70%, 95%, and 2 changes of 100% ethanol), followed by final embedding. Qualitative assessment was performed based on size, morphology, quantity and distribution.

### Neutral lipid and mitochondrial potential assays

Lipid droplet (LD) and mitochondrial potential were quantified using FACS. Cells were collected and incubated with 3.7 µM BODIPY (D3922) or 50 nM DiIC_1_(5) (M34151) for 30 minutes at room temperature (RT) or 37°C 5% CO_2_, respectively, protected from light. For FACS, single-cell data were acquired using an Attune flow cytometer (Thermo Fisher Scientific) and analyzed using FCS Express 7 (De Novo Software). Cells were gated based on forward scatter and singlet discrimination. Non–treated cells processed in parallel were used to define the negative gate.

### Cholesterol effux assay

Cholesterol eflux was measured using a Cholesterol Efflux Fluorometric Assay kit (Biovision, K582-100) following the manufacturer’s instructions. Cells were seeded in a 96-well plate at 50,000 cells/well. Cells were labeled with a Labeling Reagent for 1 h at 37°C followed by loading cells with Equilibration Buffer. After overnight incubation, media containing Equilibration Buffer was aspirated and replaced with media containing a cholesterol acceptor human HDL (40 µg/ml) for 5 h at 37°C. Following incubation, supernatants were transferred to flat bottom clear 96-well white polystyrene microplates (Greiner Bio-one, 655095). Adherent cell monolayers were lysed with Cell Lysis Buffer and incubated for 30 min at RT with gentle agitation followed by pipetting to disintegrate cells. Cell lysates were transferred into flat bottom clear 96-well white polystyrene microplates. Fluorescence intensity (Ex/Em=485/523nm) of supernatants and cell lysates was measured using a Varioskan LUX multimode microplate reader (Thermo Fisher Scientific, VL0000D0). Percentage of cholesterol efluxed was quantified: % cholesterol efflux = Fluorescence intensity of supernatant / fluorescence intensity of supernatant plus fluorescence intensity of cell lysate x 100.

### Cytokine measurements

Media from THP1 macrophages or WTC11 iMGLs were collected and centrifuged at 300xg for 5 min to eliminate floating cells. The supernatant was transferred to a clean tube and stored at −80°C until use. A custom 10-plex MILLIPLEX Human kit (Millipore; HCYTA-60K-10) was used to determine the levels of IL-6, IL-8, IL-10, IP-10, MCP-1, MDC, MIP-1α, MIP-1β, RANTES, and TNF-α.

### Efferocytosis assays (myelin, lysosensor, lysotracker and DQBSA)

Myelin fragments were isolated from human brain tissue (corpus callosum) as described in [57]. Isolated myelin fragments were labeled with pHrodo dye (Thermo Fisher Scientific, cat. P36600) in PBS for 30 min in the dark at RT, followed by two washes in PBS. Cell confluence and red fluorescence signal were quantified after 15 h of treatment (pH-rodo myelin 20 μg/ml) using the Incucyte S3 live imaging system. Total integrated density was calculated as mean red fluorescence intensity multiplied by surface area of masked object [RCU x µm^2^] and normalized by cell confluence (phase channel).

THP-1 macrophages and WTC11 iMGLs were incubated with 75 nM LysoTracker-Red (Thermo Fisher Scientific, L7525) for 20 min at 37°C followed by 1 µM LysoSensor-Green (Thermo Fisher Scientific, L7535) for 1 min at 37°C. To characterize hydrolytic capacity of lysosomes, cells were incubated with 1µg/ml DQ Red BSA (Thermo Fisher Scientific) for 24h at 37°C. After collecting the cells, single-cell data were acquired using an Attune flow cytometer (Thermo Fisher Scientific) and analyzed using FCS Express 7 (De Novo Software).

### Xenotransplantation

iPSC-derived hematopoietic stem cells (HPCs) from WT and LACTB KO WTC11 line (two clones per genotype) were transplanted into 5xFAD-hCSF1 (n=16 WT, n=17 LACTB KO HPCs) mice in accordance with previous publications [47,58,59]. P2 pups from the transplantation-competent 5xFAD-hCSF1 mouse strain were placed on ice to induce hypothermic anesthesia. The site of injection was disinfected. In between donor HPCs, needles were cleaned with consecutive washes of PBS, 70% ethanol, followed by PBS. Tools were kept sterile. Free-hand transplantation was performed using a 30-gauge needle affixed to a 10 μl Hamilton syringe. Mice received 1 μl of HPCs suspended in 1x DBPS at 62.5K cells/μl at each of 8 injection sites, totaling 500k cells/pup. Bilateral injections were performed at 2/5th of the distance from the lambda suture to each eye, injecting into the lateral ventricles at 3 mm and into the overlying anterior cortex at 1 mm, and into the posterior cortex in line with the forebrain injection sites, and perpendicular to lambda at a 45° angle. Transplanted pups were then returned to their home cages and weaned at P21.

*Brain tissue dissociation and human microglia isolation from xenotransplanted mouse brain*.

Six and half months after HPC transplantation, mice were anesthetized and intracardially perfused with 1X DPBS, half brains were dissected, the cerebellum was removed, and tissue was stored in RPMI 1640 until subsequent perfusions were completed. All steps were performed on ice or at 4°C with ice-cold reagents and all centrifuge steps were performed for 10 min at 400xg with full brake and acceleration unless otherwise stated. Brains were manually homogenized using a 7 mL Dounce homogenizer by adding 4 mL of RPMI 1640 and performing 10 strokes with the “loose” pestle (A) followed by 10 strokes with the “tight” pestle (B). Samples were then run through a pre-soaked 70 μM filter then the filter was washed with 10mL of RPMI 1640. The sample was pelleted by centrifugation and myelin was removed by resuspension in 8mL of 30% Percoll overlaid with 2 mL of 1X DPBS centrifuged at 400xg for 20 min with acceleration set to 0 and brake set to 4. The myelin band and supernatant were discarded, and cell pellets were resuspended in 70 μL MACS buffer (0.5% bovine serum albumin in 1X DPBS) + 30 μL Mouse Cell Removal beads (Miltenyi) and incubated at 4°C for 15 min. Magnetically labeled mouse cells were separated using LS columns and the MidiMACs separator (Miltenyi) while the unlabeled human cells were collected in the flow through. Human cells were then pelleted by centrifugation and transferred to −80°C for omics analysis.

### Immunohistochemistry (IHC)

Six-month-old mice were perfused with PBS and a hemibrain was fixed in PFA 4%, cryoprotected in sucrose solution and stored at −80°C. Hemibrains were coronally sectioned in a cryostat (30 μm) and stored as free-floating sections in a cryoprotective solution at −20°C.

For IHC, 4 sections per mouse were selected and placed on a 24-well plate. Tissue sections were washed three times in TBS-Tween (TBST) buffer for 5 min each. Only for Ku80/IBA1 staining, a 20-minute incubation in pretreatment buffer (0.6% H_2_O_2_, 0.1% Triton-X in TBST) and antigen retrieval (incubation at 80°C for 30 min in a heat block in sodium citrate buffer pH=6) steps were performed. Next, sections were washed three times in TBS-Tween, and incubated in blocking solution (5% normal donkey serum in TBST + 0.3% Triton X100) for 2 h. Sections were incubated in primary antibodies overnight at 4°C. Specifically for the Ku80/IBA1, sections were in blocking buffer overnight and the incubation with primary antibodies was at 37°C for 2 h on a thermoblock. After three washes with TBST buffer, sections were incubated for 1.5 hours at room temperature with secondary antibodies (Thermo Scientific 1:200) diluted in TBST. For additional staining for amyloid pathology, sections were washed again in TBST buffer three times and incubated with the antibody Aβ42 (β-Amyloid clone D54D2 Alexa Fluor 488 Conjugate, Cell Signaling Technologies #51374S, 1:250) overnight; or with Thioflavin S (Sigma-Aldrich #T1892, 1:10,000) for 2 min. In all cases, sections were washed again and stained with DAPI (ThermoFisher #D1306, 300 μM) for 5 min at room temperature. Sections were washed 3 times and mounted using PBS onto superfrost slides with TrueBlack mounting media (Biotium).

### Image analysis

A Keyence microscope was used to image full brain sections (single-plane 20x stitched images), except for microglia-plaque association and plaque-compaction analyses, where a Zeiss LSM900 confocal microscope was used to acquire z-stack 20 slides 20X images 2×3 tiles. A custom ImageJ Fiji [60] macro plugin was generated for each analysis. To calculate engraftment efficiency, the number of human microglia (Ku80^+^ cells) was divided by the total microglia number (IBA1^+^ cells).

To study the number of plaque-associated microglia Imaris software was used for 3D reconstruction. Volume masks were generated for microglia and plaques using a Gaussian filter and specific thresholds. The number of microglia associated with amyloid plaques was manually counted and divided by the corresponding plaque volume. We analyzed 3-4 brain sections per mouse.

### Statistical analysis

Data were visualized and analyzed using R [61]. Differences in marginal means between treatment groups were estimated and tested using the emmeans v1.11.2-8 R package [62] with appropriate linear mixed-effects regression models conditioned on the data using the lmer R package [63]. Batch, iPSC clone, and mouse identifiers were added as covariates in statistical models when appropriate; the model used for each analysis is provided in Table 1.

**Table 1.** Statistical analyses report. Details of the linear regression models used and corresponding outputs for each analysis in main and supplementary figures.

## Results

### Lower LACTB expression in myeloid cells leads to higher succinylcarnitine levels and lower Alzheimer’s disease risk

To evaluate the potential contribution of APH1B and LACTB to AD risk within the chromosome 15q GWAS locus, we performed Mendelian Randomization (MR) analyses integrating AD GWAS associations with gene expression and metabolite abundance quantitative trait loci (e/mQTLs). As shown in Figure 1A, our pairwise MR analyses show a causal chain whereby lower genetically encoded LACTB mRNA expression in myeloid cells (data from STARNET monocyte-derived-macrophage [44]) leads to higher succinylcarnitine levels in the CSF (dataset from [30]) and lower AD risk (dataset from [13]). We did not find a statistically significant association between genetically encoded APH1B mRNA expression in myeloid cells and either succinylcarnitine levels in the CSF or AD risk (Additional file 1). This does not exclude the possibility that APH1B may play a causal role in modulating AD risk in other cell types/states. Although LACTB is expressed in several cell types within the brain, it is more highly expressed in the immune cluster compared to other cell populations, in contrast to the expression of other mitochondrial proteins such as SDHA, dataset from [64] (Figure 1B). In addition, we tested whether there is an upregulation of LACTB when cells are differentiated into macrophages or microglia *in vitro*. Upon differentiation of THP1 monocytes towards macrophages or induced-pluripotent stem cells (iPSCs) to derived microglia (iMGLs), we observed an increase in LACTB expression at both mRNA and protein levels (Figure 1C, D). Consistent with this, the Abud et al., 2017 dataset [65], interrogated via Stemformatics website [66], confirmed increased LACTB expression in myeloid cells compared to hematopoietic progenitor cells (HPCs) or to iPSC (Supplementary Figure 1A).

**Figure 1.**
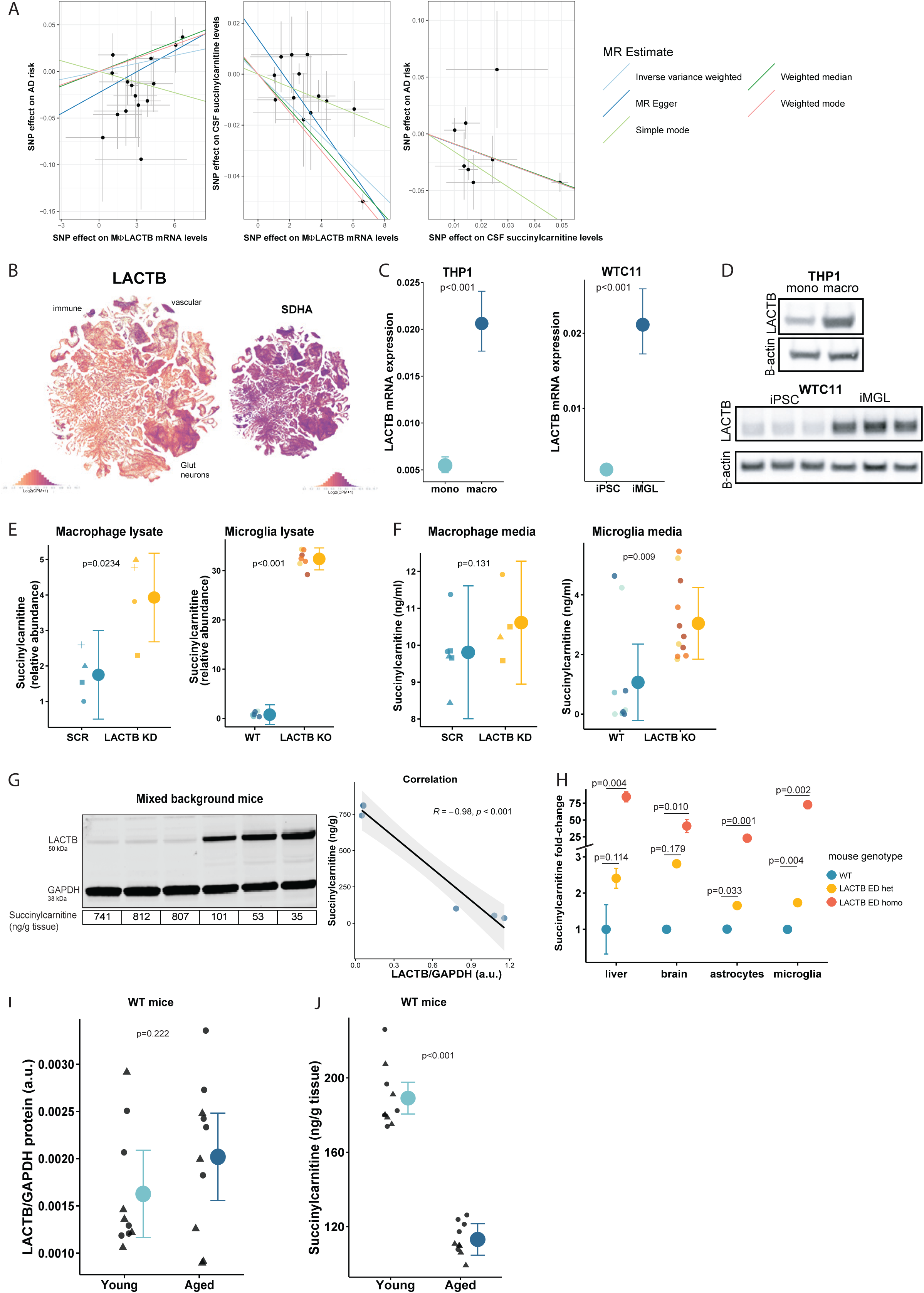
Lower LACTB expression in myeloid cells leads to higher succinylcarnitine levels and lower AD risk. A) Genetically predicted lower LACTB expression—instrumented using macrophage cis-eQTLs at the LACTB locus—is associated with lower AD risk (left panel). Genetically predicted lower LACTB expression is also associated with higher succinylcarnitine levels in the CSF (middle panel). Furthermore, genetically predicted higher succinylcarnitine levels are associated with lower AD risk (right panel). B) LACTB is expressed higher in the immune cluster compared to other brain cells, in contrast to another mitochondrial protein (SDHA), data from Brain Atlas [64]. C, D) LACTB mRNA (C) and protein levels (D) are increased upon differentiation (THP1 macrophages vs monocytes and WTC11 iMGLs vs iPSC). E) Succinylcarnitine levels are increased in the lysate of LACTB KD/KO myeloid cells compared to SCR/WT. F) Succinylcarnitine is increased in the cell culture media of LACTB KD/KO myeloid cells compared to SCR/ WT. G) Higher succinylcarnitine levels in mice with lower LACTB expression (C57B6 x SLJ background, 2 months old). H) Higher succinylcarnitine levels in the liver, brain, isolated microglia and astrocytes of LACTB enzymatically-dead (ED) mice compared to WT (n=2 mice per genotype, 3 months old). I) LACTB protein levels increase with age in WT mice (12 vs 2 months old). J) Succinylcarnitine levels decrease with age in WT mice (12 vs 2 months old). Graphs display individual data points (left) alongside estimated marginal means with 95% confidence intervals (right). For the raw data, dot shapes represent independent macrophage differentiations, and dot colors indicate distinct microglia clones. Statistical details are provided in Table 1.

Next, to investigate the impact of reduced LACTB levels, we generated LACTB knock-down (KD) THP1 macrophages and LACTB knock-out (KO) WTC11 iMGLs (Supplementary Figure 1B-F). In both models, we observed a significant increase in succinylcarnitine levels in cell lysates upon LACTB KD/KO (Figure 1E). We also observed a concordant increase in succinylcarnitine levels in the cell culture media from LACTB KD THP1 macrophages and LACTB KO WTC11 iMGLs compared to scrambled (SCR) and wild-type (WT) controls, respectively (Figure 1F).

We further validated the causal link between LACTB expression levels and succinylcarnitine levels *in vivo*. First, we used a mixed background (C57BL/6 x SJL) mouse model, where the SJL background includes a mutation that reduces LACTB protein levels (unpublished work [67]). We observed that there are mice with high or low LACTB levels in the brain lysate within the same litter, which was inversely correlated with succinylcarnitine levels (Figure 1G). Second, we generated a mouse model with a S162I mutation, inactivating the key residue within the LACTB catalytic pocket [2,68], thereby leading to a LACTB enzymatically-dead (ED) mouse model. This mutation does not alter LACTB mRNA expression in total brain lysate (hippocampus), isolated microglia or isolated astrocytes (Supplementary Figure 1G). However, using an antibody directed against an epitope located in the active site, we observed a mutation-dependent decrease in LACTB detection when one or both alleles were mutated (Supplementary Figure 1H). Succinylcarnitine levels in liver, brain lysate (posterior cortex), isolated microglia and isolated astrocytes were increased upon LACTB inactivation in a dose dependent manner (wild-type (WT) LACTB^+/+^< Het LACTB^ED/+^< Hom *LACTB^ED/ED^* mice) (Figure 1H, n=2 mice per group), corroborating that LACTB is a major determinant of succinylcarnitine levels *in vivo*.

As aging is the primary risk factor for AD, we measured the levels of LACTB and succinylcarnitine in aged mice. We observed a non-statistically significant increase in LACTB levels (Figure 1I) and a significant decrease in succinylcarnitine (Figure 1J) in the brain of aged mice (12 months old) compared to young (2 months old) WT mice.

In summary, we have demonstrated that lower LACTB levels in myeloid cells, which are associated with lower AD risk, or inactivation of LACTB enzymatic activity lead to higher levels of succinylcarnitine in mouse and human models.

### LACTB directly cleaves succinylcarnitine and succinylation inhibits LACTB activity

Next, we investigated LACTB enzymatic activity and its relationship to succinylcarnitine cleavage. LACTB possesses DAEP activity, cleaving proteins on the C-terminal side of D-Asp residues [3]. Using a succinyl- D-aspartate fluorescent substrate, we confirmed this activity in myeloid cells, and observed a marked reduction in substrate cleavage following LACTB KD in human THP1 macrophages, and a complete lack of cleavage in human LACTB KO iMGLs (Figure 2A, left and middle panels, respectively). This enzymatic activity is conserved in mice as observed in bone marrow-derived macrophages (BMDMs). LACTB ED heterozygous or homozygous animals exhibited partial or complete loss of DAEP activity, respectively, in BMDMs, relative to WT mice (Figure 2A, right panel).

**Figure 2.**
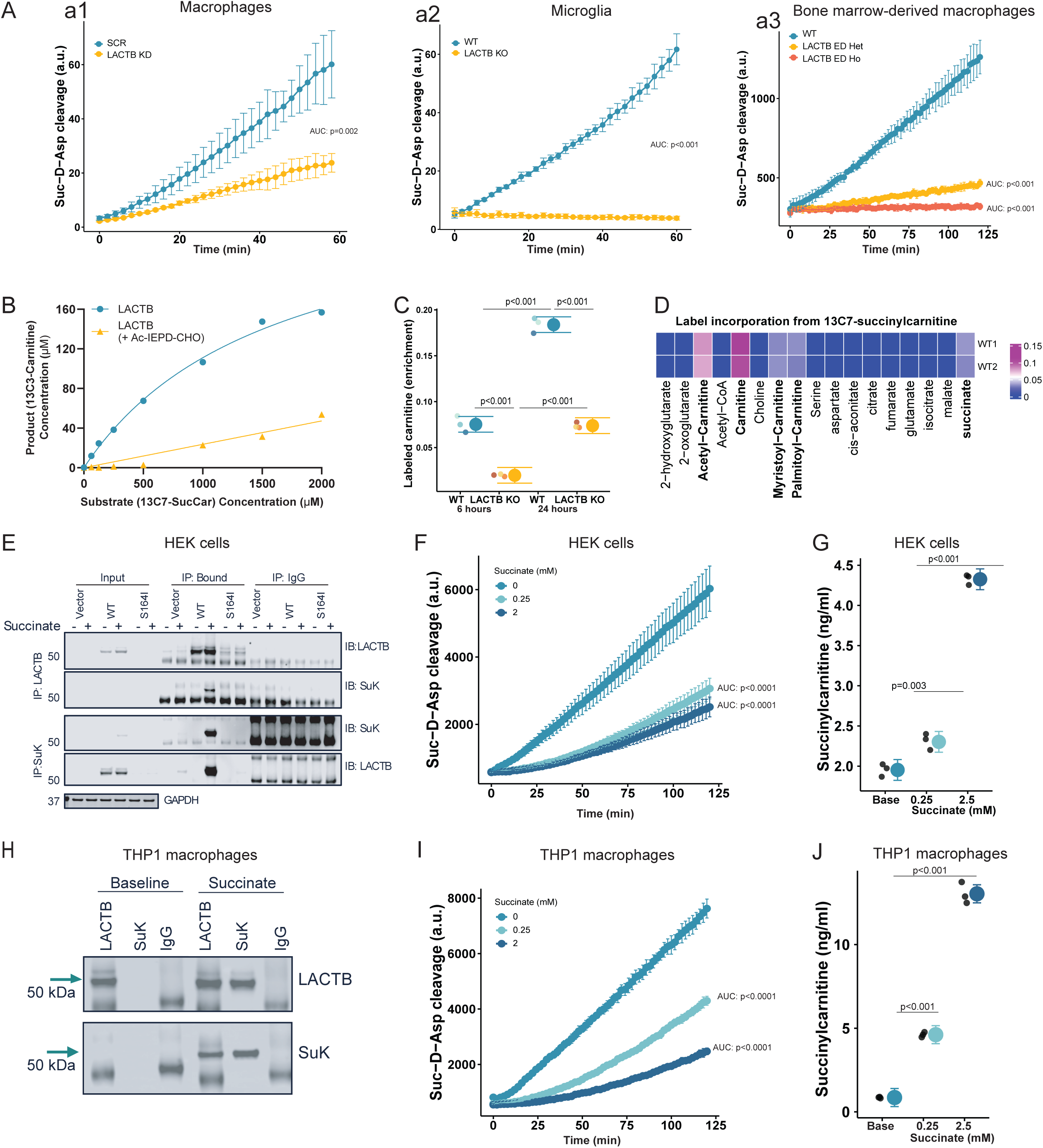
LACTB directly cleaves succinylcarnitine and succinylation inhibits LACTB. **A)** Representative curves of succinyl-D-Asp substrate cleavage over time, showing strong reduction/abolishment in LACTB KD/KO myeloid cells compared to SCR/WT (a1: n=3 THP1 macrophages differentiations, a2: n=3 clones per genotype WTC11 iMGLs, a3: mouse BMDMs n=6 mice per genotype). Area under the curve (AUC) was calculated and used as outcome variable in statistical analyses. B) ^13^C_3_-carnitine generation after substrate (^13^C_7_-succinylcarnitine) incubation with LACTB recombinant protein in the absence or presence of LACTB inhibitor (Ac-IEPD-CHO) in a cell-free assay. C) Reduced label incorporation into carnitine in LACTB KO iMGLs after incubation with ^13^C_3_-succinylcarnitine (n=3 clones per genotype) compared to WT WTC11 iMGLs. D) Representative heatmap showing label incorporation from ^13^C_7_-succinylcarnitine into the indicated metabolites in WT iMGLs. E-J) LACTB is succinylated after treatment with diethyl-succinate (5 μM for 4 hours) in HEK cells (n=2 transfections, 4-10 well replicates each) (E) and THP1 macrophages (n=2 differentiations, 4 well replicates each) (H). Succinylation of LACTB is associated with a reduced rate of succinyl-D-Asp cleavage in HEK cells (n=2 transfections, 3 well-replicates each) (F) and THP1 macrophages (n=2 differentiations, 3 well-replicates each). AUC was calculated and used as outcome variable in statistical analyses. (I) LACTB succinylation is associated with higher succinylcarnitine levels in HEK cells (n=3) (G) and THP1 macrophages (n=3) (J). Graphs display individual data points (left) alongside estimated marginal means with 95% confidence intervals (right). For the raw data, dot shapes represent independent macrophage differentiations, and dot colors indicate distinct microglia clones. Statistical details are provided in Table 1.

To investigate whether LACTB directly cleaves succinylcarnitine, we performed a cell-free assay using in-house generated recombinant LACTB and a custom ^13^C_7_-labeled-succinylcarnitine substrate (all carbons labeled, TRC-S102985, LGC). This tracer allows for direct quantification of the cleavage product. Incubation of recombinant LACTB (100 nM) with increased concentrations of ^13^C_7_-labeled- succinylcarnitine for 2 hours yielded a dose-dependent increase in the cleavage product ^13^C_3_-carnitine (Figure 2B), confirming direct LACTB enzymatic cleavage of succinylcarnitine. Importantly, this cleavage was abolished when recombinant LACTB protein was preincubated with a LACTB inhibitor (Ac-IEPD- CHO), further demonstrating that succinylcarnitine cleavage is dependent on LACTB activity.

To confirm these findings in a biological system, we conducted isotope tracing experiments in WT and LACTB KO iMGLs with commercially available ^13^C_3_-succinylcarnitine (TRC-S688832, LGC). As in the recombinant LACTB assay, we hypothesized that, after succinylcarnitine cleavage, there would be an increase in labeled carnitine (^13^C_3_-carnitine). Consistent with our cell-free results, we observed a progressive increase in the pool of labeled carnitine after 6 and 24 hours in WT iMGLs (Figure 2C). At both time points, there was a significant reduction in the abundance of labeled carnitine in LACTB KO iMGLs compared to control iMGLs (n=2 clones per genotype).

Finally, we utilized the ^13^C_7_-labeled-succinylcarnitine tracer to track the metabolic fate of both the carnitine and succinyl moieties following cleavage. After 24 hours of treatment in WTC11 iMGLs, we detected the expected labeling of carnitine, alongside modest enrichment in acetyl-carnitine, myristoyl- carnitine, palmitoyl-carnitine, and succinate (Figure 2D, n=2 clones). Taken together, experiments with purified protein and isotopic tracing demonstrate that LACTB not only cleaves succinylcarnitine under cell-free biochemical conditions but is also a major determinant of its catabolism in cells.

Lysine succinylation is a recently described post-translational modification that regulates protein function. Prior work has shown that LACTB activity is diminished when LACTB is succinylated in cancer cells [69], which may be a negative feedback loop regulating the cleavage of succinylcarnitine by LACTB. To explore LACTB succinylation, we transfected HEK cells with empty, LACTB WT or LACTB ED (S164I) vectors and treated them with diethyl succinate (2 mM). We observed an increase in LACTB succinylation in HEK cells overexpressing LACTB WT protein after the treatment, but not in HEK cells expressing LACTB ED (Figure 2E), indicating that disruption of LACTB active site impairs this post-translational modification. To investigate how succinylation influences LACTB enzymatic function, we measured LACTB activity (succinyl-D-aspartate fluorescent assay) and levels of succinylcarnitine. LACTB succinylation reduced the cleavage of the succinyl-D-aspartate substrate and increased succinylcarnitine levels, indicating inhibition of LACTB (Figures 2F and G). Similarly, we observed that treatment with diethyl succinate led to LACTB succinylation in macrophages (Figure 2H) and a corresponding reduction of LACTB activity, measured by succinyl-D-aspartate cleavage and succinylcarnitine levels (Figures 2I and 2J).

In short, we demonstrated that LACTB directly cleaves succinylcarnitine and that succinylation is a post-translational modification that inhibits LACTB activity.

### LACTB KD/KO in myeloid cells alters expression of genes in pathways related to oxidative phosphorylation, protein translation, cell cycle and immune response

To investigate the transcriptional consequences of LACTB KD/KO in myeloid cells, we performed bulk RNA-seq in our cell models. In THP1 macrophages (n=4), LACTB KD led to the upregulation of 82 genes (logFC > 1, p.value <0.05) and downregulation of 66 genes (logFC < −1, p.value <0.05) (Table 2). We performed Gene Set Enrichment Analysis (GSEA) with the “Biological Process” subset of Gene Ontology genesets, and found a positive enrichment of pathways related to the response to interferon (IFN) and to the tumor necrosis factor (TNF) (“GOBP_response_to_virus”), oxidative phosphorylation, and protein localization to the endoplasmic reticulum, in LACTB KD macrophages compared to macrophages treated with scrambled siRNA (SCR) as controls (Figure 3A and Table 2). We also found a negative enrichment of proliferation-related pathways (Figure 3A and Table 2).

**Figure 3.**
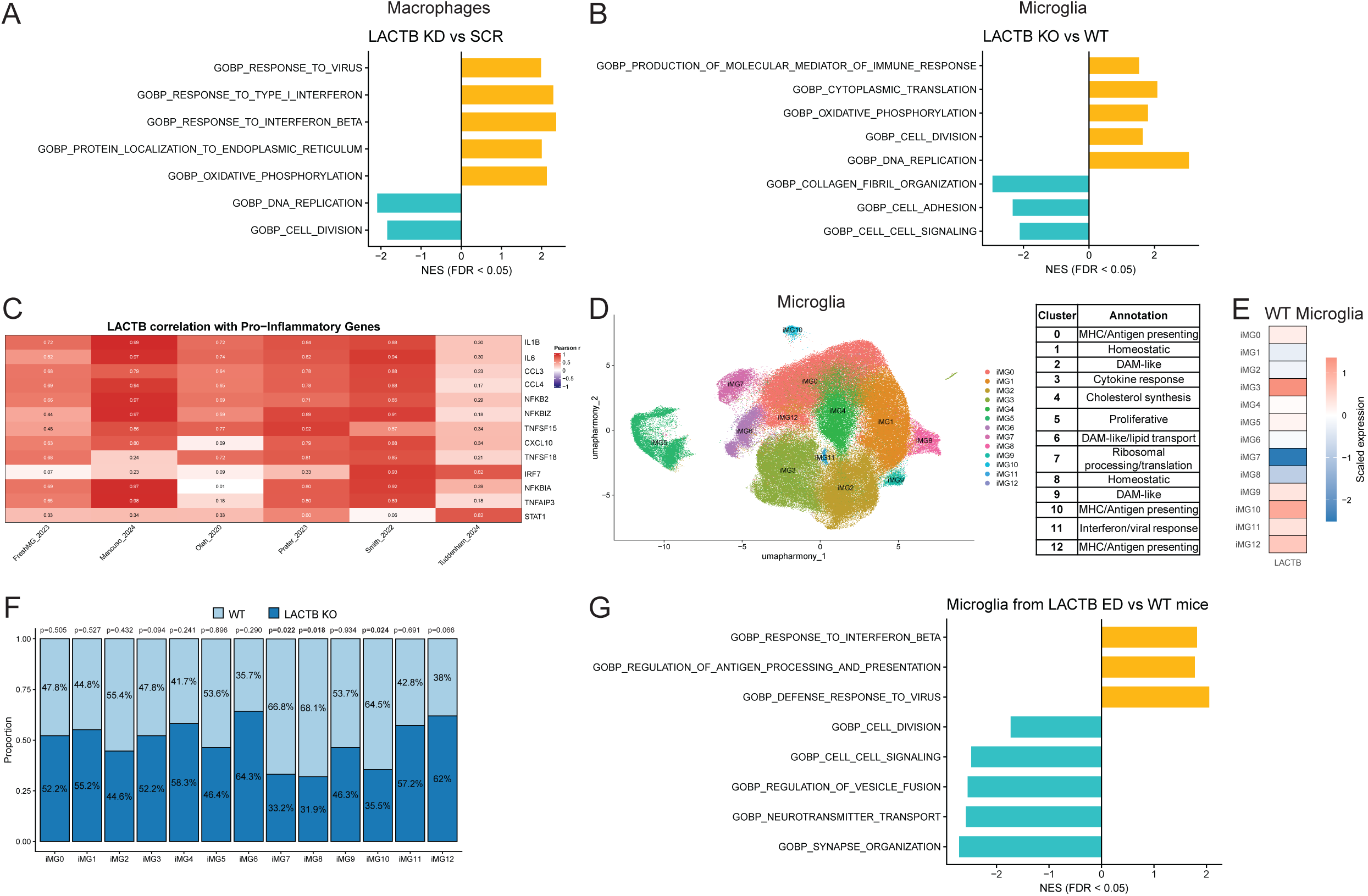
LACTB KD/KO in myeloid cells alters expression of genes in pathways related to oxidative phosphorylation, protein translation, cell cycle and immune response. A) Pathways enriched in GSEA analysis of bulk RNAseq data from LACTB KD THP1 macrophages compared to SCR (n=5 independent macrophages differentiations), NES = Normalized Enrichment Score. B) Pathways enriched in GSEA analysis of bulk RNAseq data from LACTB KO iMGLs compared to WT (3 clones per genotype, 2 differentiations per clone). C) Heatmap of Pearson correlation coefficients between LACTB expression and genes encoding pro-inflammatory cytokines across microglial states in publicly available human single-cell microglia datasets [70–75]. D) Single-cell clusters and corresponding annotations of WT and LACTB KO iMGLs (3 clones per genotype, 1-2 differentiations per clone). E) *LACTB* expression across clusters in WT iMGLs. F) Cluster proportions in WT and LACTB KO iMGLs. Cluster proportions were estimated using crumblr and compared by genotype using a generalized linear model with batch as a covariate (see Methods for further details). G) Pathways enriched in GSEA analysis of bulk RNAseq data from mouse microglia isolated from the brains of LACTB ED mice compared to controls (3 months old, n=3). Complete lists of genes and pathways are provided in Tables 2 and 3.

**Table 3.** Single cell RNAseq results. List of top markers for each cluster, statistics on cell cluster proportions and details of LACTB regulome derived from the analysis of single cell RNAseq in WTC11 iMGLs *in vitro*.

Next, we performed bulk RNA-seq on WTC11 LACTB KO and WT iMGLs (n=3 clones per genotype, 2 differentiations per clone). Similar to the observations found in THP1 macrophages, there was a positive enrichment of pathways related to the immune response, oxidative phosphorylation, and protein translation in LACTB KO iMGLs compared to isogenic controls (Figure 3B). In contrast, we found that complete loss of expression of LACTB led to a positive enrichment of cell proliferation-related pathways (“GOBP_DNA_replication”), compared to the negative enrichment observed in LACTB KD macrophages. Also, complete loss of LACTB led to a negative enrichment of cell adhesion and cell signaling pathways (Figure 3B).

To examine the relevance of these findings based on macrophage/microglia-like cells *in vitro* to microglial cells *in vivo* under disease conditions, we interrogated and re-analysed six publicly available single-cell RNAseq microglial human AD datasets [70–75], and performed correlation analyses to identify genes whose expression is associated with LACTB across microglial states. Our analysis focused on determining associations with leading-edge genes from significantly enriched pathways identified by GSEA of our bulk RNA-seq data derived from THP1 KD and iMGL KO systems. Strikingly, LACTB expression was most often strongly associated with genes implicated in pro-inflammatory pathways identified in both THP1 and iMGL leading-edge analyses, including IL1B, IL6, CCL3, and CCL4 (Figure 3C, Supplementary Figure 2A), which encode pro-inflammatory cytokines [76]. These findings further support the idea that LACTB acts as a modulator of immune responses in myeloid cells. Genes consistently negatively correlated with LACTB expression included the AD risk gene ABCA7, identified in GSEA leading-edge analyses for both THP1s and iMGLs (Supplementary Figure 2A), and implicated in pathways such as lipid metabolism and phagocytosis [77–79]. Additional negatively correlated genes included those involved in cell cycle regulation identified in our THP1 and iMGL leading-edge analyses, such as CDKN2C [80], MCM2 [81], and PCNT [82], implicating a role for LACTB in cell-cycle regulation in an *in vivo* context.

Single-cell RNAseq on WT and LACTB KO microglia (3 clones per genotype, 1-2 differentiations per clone) also implicated LACTB in the immune response. Across the 12 clusters identified (Figure 3D and Table 3), we found that LACTB was most highly expressed in clusters enriched for cytokine response (iMG3) and MHC/antigen presenting (iMG10) processes (Figure 3E and Table 3). In agreement with our bulk RNAseq results in microglia, our single-cell data showed alterations in the cell proportion for clusters enriched for ribosomal processing and translation functions (iMG7), and related to the immune response (iMG10, iMG12) (Figure 3F and Table 3).

Finally, we explored the *in vivo* effects of loss of LACTB enzymatic activity by performing bulk RNA sequencing and GSEA analysis of mouse microglia isolated from LACTB ED (homozygous) and control mice (3 months old, n=3) (Figure 3G and Table 2). In line with the *in vitro* results, we observed a positive enrichment of pathways related to IFN/TNF response and antigen-presentation. Negatively enriched pathways included proliferation, cell signaling and vesicle fusion.

### LACTB KD/KO in myeloid cells increases oxidative phosphorylation and reduces protein synthesis, and triacylglycerides

Previous reports have linked LACTB to differentiation and proliferation processes in cancer models [7]. Also, our data showed an increase in LACTB gene and protein expression after differentiation from THP1 monocytes to macrophages and iPSC to iMGLs (Figure 1C, D). To determine whether the observed differences were due to alterations in the differentiation process, we examined the expression of microglia/macrophage markers. As shown in Supplementary Figure 2B, LACTB KD/KO did not induce statistically significant changes in the mRNA expression of microglia/macrophage markers (CSF1R, P2RY12, CX3CR1, TGFBR1) in either THP1-derived macrophages or WTC11 iMGLs. In addition, as proliferation pathways were negatively/positively enriched in LACTB KD/KO macrophages/microglia, respectively, we analyzed cell division using a DNA intercalating agent (EdU). A statistically significant reduction in proliferation was observed in LACTB KD THP1 macrophages compared to SCR (Supplementary Figure 2C, left panel), as predicted by the transcriptional analysis (Figure 3A). However, no statistically significant changes were observed between LACTB KO and WT WTC11 iMGLs (Supplementary Figure 2C, right panel).

Our transcriptomic data also showed an enrichment in the oxidative phosphorylation pathway upon LACTB KD/KO in myeloid cells, so we measured mitochondrial respiration through Seahorse assays. Importantly, we observed that LACTB depletion increased basal and maximal respiration in human myeloid cells (Figure 4A). Given that LACTB forms filaments in the mitochondrial intermembrane, we also examined other mitochondrial features. LACTB KD/KO myeloid cells showed a non-statistically significant reduction in mitochondrial membrane potential compared to SCR/WT (Supplementary Figure 3A). No major qualitative morphological alterations in mitochondria were observed in LACTB KO microglia compared to WT, by electron microscopy (Supplementary Figure 3B). Our results suggest that LACTB loss enhances bioenergetic capacity without causing gross alterations to mitochondrial structure. Changes in cellular metabolism can directly impact protein synthesis, as energy and nutrient availability regulate translation initiation [83]. Consistent with this, our bulk RNAseq analysis of LACTB KD/KO m y e l o i d c e l l s r e v e a l e d e n r i c h m e n t o f p a t h w a y s r e l a t e d t o “GOBP_protein_localization_to_endoplasmic_reticulum” and “GOBP_cytoplasmic_translation” (Figure 3A and 3B). To further investigate this relationship, we identified genes co-expressed with LACTB using single-cell RNAseq data from WTC11 WT iMGLs (Table 3) and ARACNe [35], a network-based analysis tool that infers gene co-expression modules from single-cell RNA-seq data [84,85]. Module scores were then calculated across microglial clusters. We observed that genes co-expressed with LACTB were most enriched in cluster iMG7 (Supplementary Figure 3C and Table 3), which is characterized by ribosomal biogenesis and translation-related processes (Figure 3D). Finally, to examine the functional impact of LACTB KD/KO on protein translation, we measured nascent protein synthesis using a labeled methionine analog and observed a significant reduction in protein synthesis in LACTB KD/KO myeloid cells compared to SCR/WT (Figure 4B).

**Figure 4.**
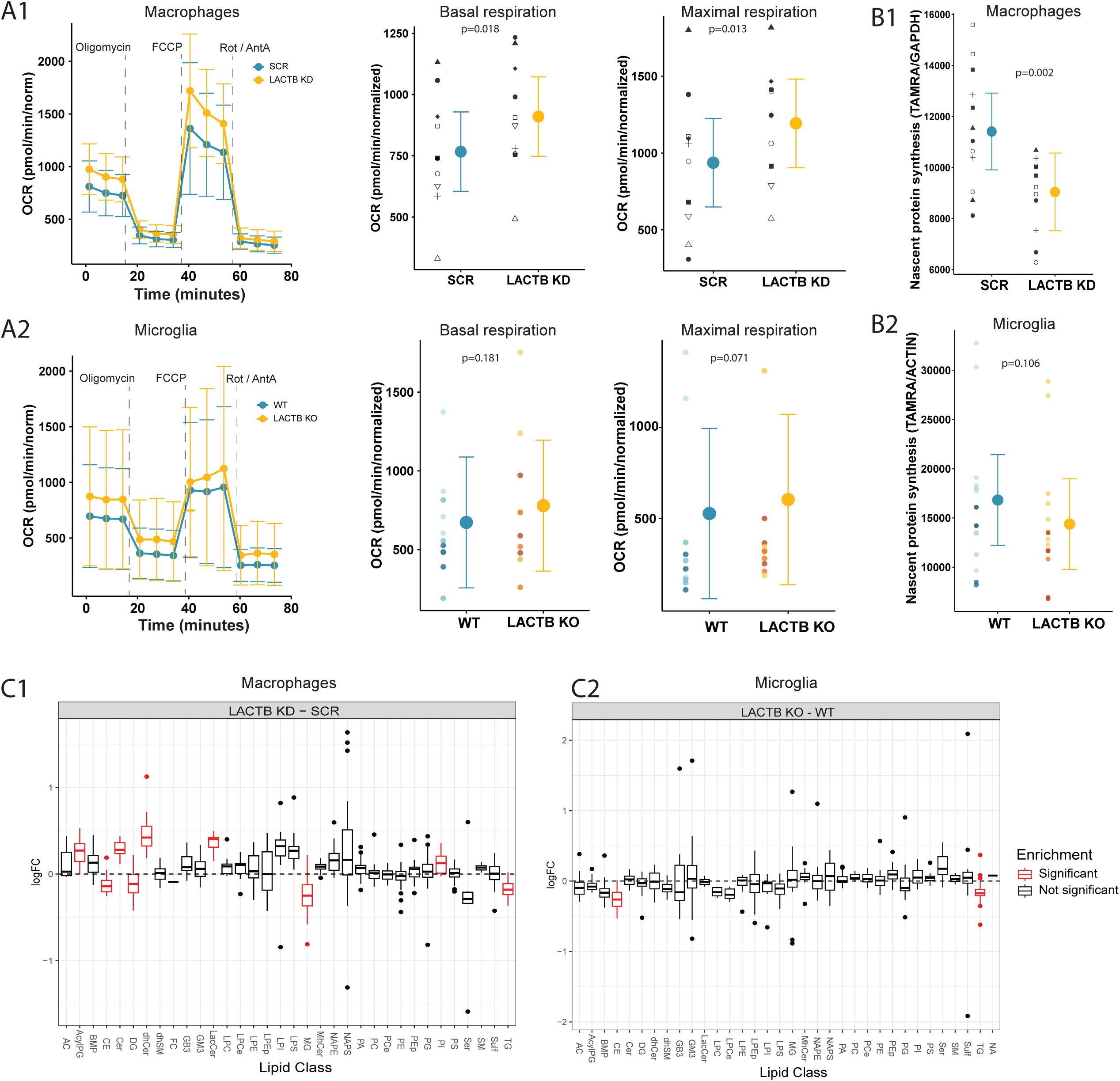
**LACTB KD/KO in myeloid cells increases oxidative phosphorylation and reduces protein synthesis, cholesteryl esters, and triacylglycerides**. A) Increased in OXPHOS (measured by mitostress seahorse assays) in LACTB KD THP1 macrophages (n=10 differentiations, 3-6 technical replicates each), and WTC11 LACTB KO iMGLs (3 clones per genotype, 3 differentiations per clone, 3-6 technical replicates each) compared to controls. B) Reduction in nascent protein synthesis (measured with a methionine analog) in LACTB KD/KO myeloid cells compared to SCR/WT. Graphs display individual data points (left) alongside estimated marginal means with 95% confidence intervals (right). For the raw data, dot shapes represent independent macrophage differentiations, and dot colors indicate distinct microglia clones. Full statistical details are provided in Table 1. C) Lipidomics alterations in LACTB KD vs SCR THP1 macrophages (n=3) and WTC11 LACTB KO vs WT iMGLs (n=3 clones, 1-2 differentiations each), showing a consistent reduction in CE and TG across the two myeloid cell types. Statistical details are provided in Table 4.

**Table 4.** Lipidomics results. Statistical outputs of the lipidomics analyses in THP1 macrophages and WTC11 iMGLs.

Our single-cell study in microglia showed a non-statistically significant increase in proportion of cells in iMG6 (a cluster related to lipid transport) in LACTB KO microglia compared to WT (Figure 3F and Table 3). In addition, LACTB has been nominated as an obesity gene in adipocytes [8] and the LACTB locus shows genome-wide significant associations with several plasma lipid/lipoprotein traits (phospholipids, cholesterol, cholesteryl esters, triglycerides, apolipoprotein A) in GWAS studies [86]. We performed lipidomics to assess how LACTB loss-of-function influences the lipidomic profile of myeloid cells (Figure 4C). Compared to SCR, LACTB KD macrophages exhibited increased levels of ceramides, including dihydroceramides (dhCer), ceramides (Cer), and lactosylceramides (LacCer) (Figure 4C1). Alterations in ceramide abundance may reflect changes in lysosomal function [87]. In contrast, ceramide levels were unchanged in LACTB KO iMGLs compared to WT (Figure 4C2).

In addition, we observed a decrease in cholesterol ester (CE) and in mono/di/tri-acylglycerides (MG/DG/ TG) in LACTB KD/KO myeloid cells compared to SCR/WT (Figure 4C). To further explore this finding, we measured the content of neutral lipids using BODIPY fluorescent dye (Supplementary Figure 3D). No statistically significant differences were found between LACTB KD and SCR macrophages. By contrast, LACTB KO microglia exhibited a non-statistically significant reduction in BODIPY fluorescence compared to WT. Finally, we assayed cholesterol efflux, measuring HDL-associated labeled-cholesterol in the supernatant relative to cellular content upon treatment with HDL as cholesterol acceptor. Although we observed a significant increase in cholesterol efflux in LACTB KD THP1 macrophages compared to SCR, we did not find statistically significant differences across genotypes in iMGLs (Supplementary Figure 3E). Together, these data suggest that reduced LACTB expression may contribute to lower TGs levels, potentially through enhanced cholesterol efflux or reduced neutral lipid accumulation, in a cell-type-dependent manner.

In short, we have observed that LACTB KD/KO increases oxidative phosphorylation, reduces protein synthesis and modifies lipid metabolism in myeloid cells.

### LACTB KD/KO in myeloid cells alters efferocytosis-related pathways after immune stimulation

Because pathways related to the IFN and TNF response emerged as some of the most consistently affected pathways in LACTB KD/KO/ED myeloid cells across model systems, we tested whether LACTB expression is modulated by these cytokines. Upon stimulation with IFN-β, IFN-γ, or TNF-α, we detected an increase in LACTB both at the mRNA and protein levels (Figure 5A and Supplementary Figure 4A). In line with this induction of LACTB following immune activation, succinylcarnitine levels decreased in cells treated with these cytokines (Figure 5B).

**Figure 5.**
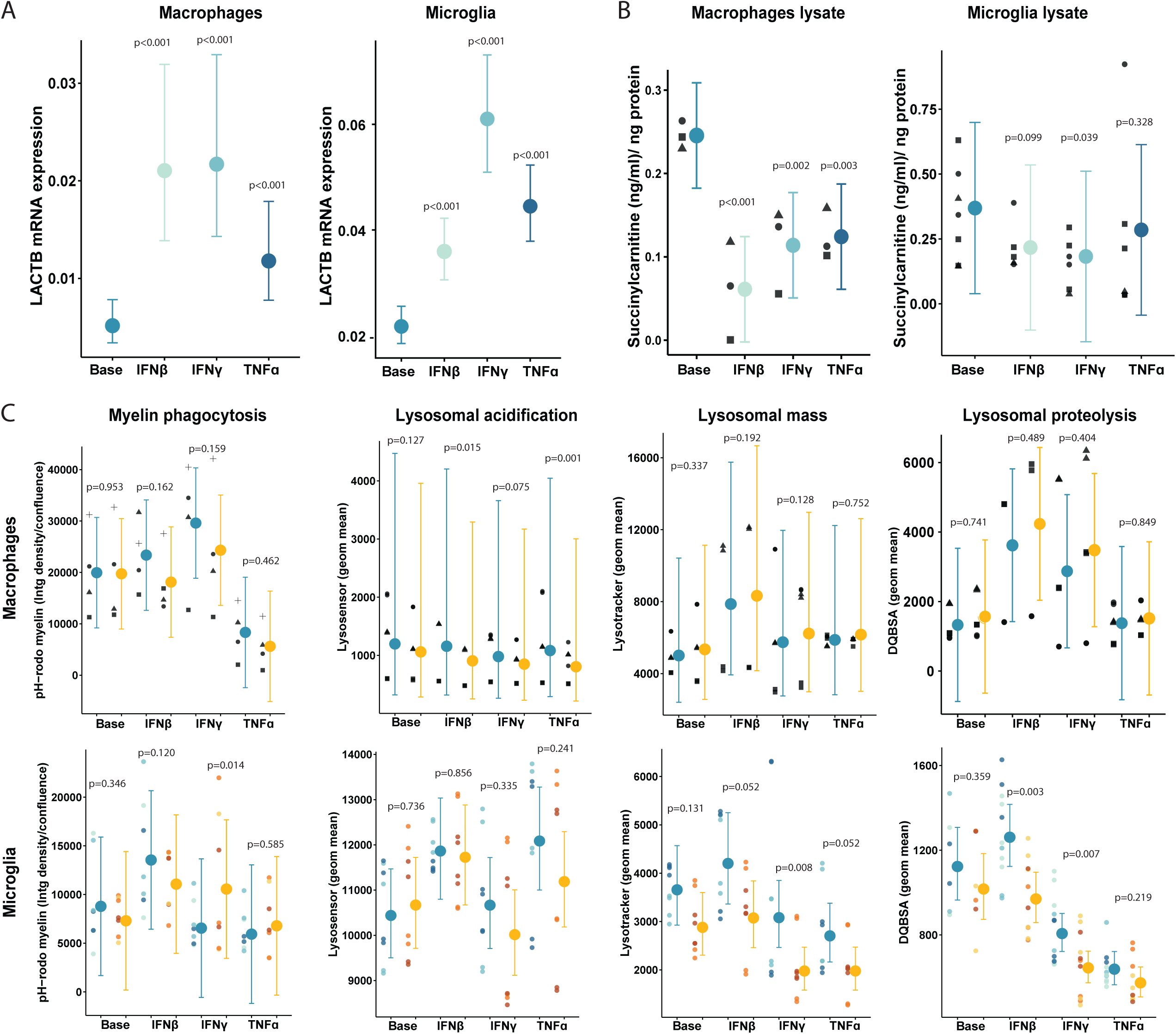
LACTB KD/KO in myeloid cells alters efferocytosis-related pathways after immune stimulation. A) LACTB mRNA expression increases after stimulation with IFN-β, IFN-γ, or TNF-α in THP1 macrophages (6 hours treatment) or WTC11 iMGLs (24 hours treatment). B) Succinylcarnitine levels decrease after stimulation with IFN-β, IFN-γ, or TNF-α in THP1 macrophages (6 hours treatment) or WTC11 iMGLs (24 hours treatment). C) Efferocytosis-related assays (myelin phagocytosis and lysosomal acidification, mass and proteolytic capacity) in LACTB KD/KO myeloid cells compared to SCR/WT. Graphs display individual data points (left) alongside estimated marginal means with 95% confidence intervals (right). For the raw data, dot shapes represent independent differentiations, and dot colors indicate distinct microglia clones. Statistical details are provided in Table 1.

To assess whether LACTB KD/KO would impair the transcriptional response to IFN-β, IFN-γ, or TNF-α treatment, we quantified the expression of IFN/TNF-responsive genes after 6–24 hours of stimulation with these cytokines (qPCRs, Supplementary Figure 4B). Overall, LACTB loss did not impair the global transcriptional response to these cytokines. However, LACTB KD THP1 macrophages showed increased induction of IL1B, STAT1, and TNF-α following IFNs treatment, accompanied by reduced expression of CLEC7A compared to SCR THP1 macrophages. LACTB KO iMGLs showed an increase in TNF-α and HLA- DQA1 compared to WT iMGLs. In addition, we analyzed cytokine secretion in cell supernatants (Supplementary Figure 4C). While LACTB reduction/loss did not broadly disrupt cytokine release in response to stimulation with IFN-β, IFN-γ, or TNF-α, THP1 LACTB KD macrophages secreted higher levels of IL-8, MCP-1, and MIP-1α (also know as CCL3, a cytokine found to be co-expressed with LACTB, Figure 3C), whereas LACTB KO iMGLs released lower levels of IL-6, IL-10, IP-10, and TNFα compared to SCR/WT cells.

As pro-inflammatory microglia activation has been proposed to be required to clear myelin [88,89], we sought to investigate the efferocytosis ability of myeloid cells after stimulation with IFN-β, IFN-γ, or TNF- α. Efferocytosis refers to the ability of macrophages to find, phagocytose, degrade, and dispose of lipid-rich cellular debris. In the brain parenchyma, efferocytosis is mainly performed by microglia and it is critically involved in AD pathogenesis [23]. We examined whether LACTB KD/KO myeloid cells exhibited altered efferocytic activity compared to SCR/WT following stimulation with IFN-β, IFN-γ, or TNF-α. In THP1 macrophages we observed that, although there were no differences between genotypes at baseline, there was a non-statistically significant reduction in myelin phagocytosis and a statistically-significant decrease in lysosomal acidification (lysosensor marker) in LACTB KD macrophages compared to SCR after stimulation with IFN-β, IFN-γ, or TNF-α (Figure 5D). In LACTB KO iMGLs, we observed a decrease in the lysosomal mass (lysotracker marker) and lysosomal proteolytic capacity (DQBSA processing) in LACTB KO iMGLs compared to WT after stimulation with these cytokines.

In both cell systems, LACTB KD/KO affected some steps of the efferocytosis pathway. These results are in line with the reduction in phagocytic and lysosomal capacity observed in DLAM THP1 macrophages [90] and PU1 KD BV2 cells [91], both thought to be associated with AD protection.

### LACTB KO human microglia show increased clustering around amyloid plaques in a mouse model of amyloid deposition

Finally, to address whether LACTB KO in human microglial cells would modify their function related to amyloid pathology *in vivo*, we xenotransplanted WT and isogenic LACTB KO (2 clones per genotype) microglia precursors into the brain of newborn hCSF1-5xFAD female mice [47,58], which differentiated into microglia. Mice were sacrificed at 6 months old (Figure 6A) and we confirmed engraftment of human microglia in the mouse brain, without significant differences between LACTB KO vs WT xenotransplanted microglia (Figure 6B). As shown in Figure 6C, isolated LACTB KO human microglia showed alterations in the metabolic panel compared to control human microglia, including a significant increase in acetyl-CoA and a non-statistically significant decrease in α-ketoglutarate, choline and serine. Due to the low sample input, we could not detect succinylcarnitine levels. However, we observed a decrease in carnitine, acetyl-carnitine, palmitoyl-carnitine and succinate in LACTB KO versus control microglia, which may be a result of lower succinylcarnitine cleavage (as shown in Figure 2C and D, *in vitro*).

**Figure 6.**
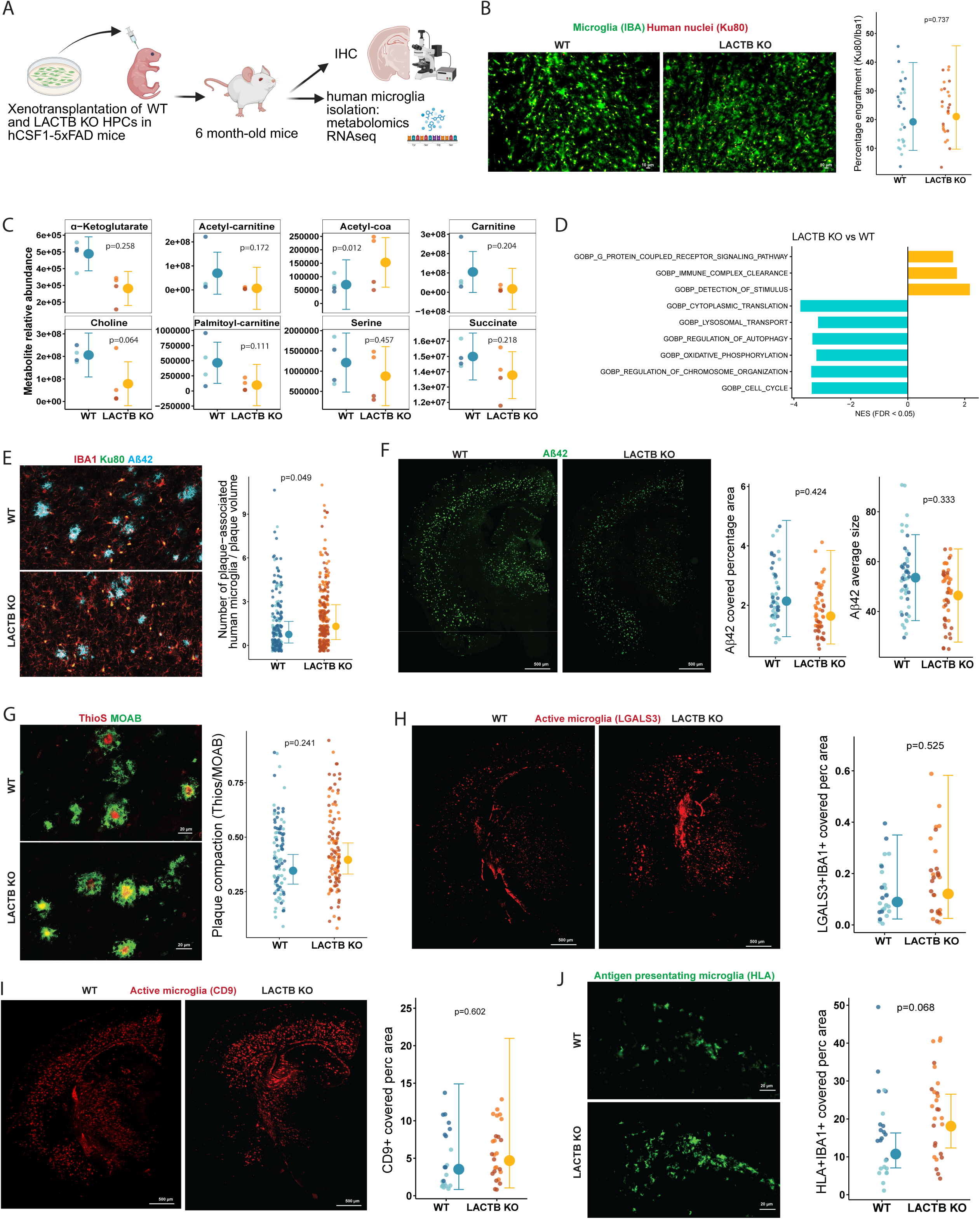
LACTB KO human microglia show increased clustering around amyloid plaques in a mouse model of amyloid deposition. A) Diagram summarizing the experimental design for xenotransplantation of human microglia precursors in mice. B) No differences in engraftment between human LACTB KO and WT microglia in hCSF1-5xFAD mice. C) Metabolomics in human LACTB KO or WT microglia isolated from the brains of hCSF1-5xFAD mice (n=2 clones per genotype, 2 mice injected per clone). D) Pathways enriched in GSEA analysis of bulk RNAseq data from human LACTB KO microglia isolated from the brain of hCSF1-5xFAD mice compared to human WT microglia (n=2 clones per genotype, 2 mice injected per clone). E-J) Immunostaining analysis of the number of human microglia (Ku80+Iba1+ cells) associated per plaque (Aβ42) (E), the area covered by plaques and the average plaque size (Aβ42) (F), the plaque compaction (ThioS/MOAB ratio) (G), and the area covered by active (LGALS3 (H), CD9 (I) markers) or antigen-presenting microglia (HLA marker (J)) microglia in hCSF1-5xFAD mice xenotransplanted with LACTB KO or WT human microglia (n=2 clones per genotype, 4-5 mice injected per clone). Graphs display individual data points (left) alongside estimated marginal means with 95% confidence intervals (right). For the raw data dot colors indicate distinct xenotransplanted microglia clones. Statistical details are provided in Table 1.

We also performed bulk RNA-seq and GSEA analysis in the isolated human xenotransplanted microglia (Figure 6D and Table 2). LACTB KO microglia were positively enriched for sensing-related pathways (e.g., “GOBP_G_protein_coupled_receptor_signaling_pathway”, “GOBP_detection_of_stimulus”, “GOBP_immune_complex_clearance”), consistent with increased surveillance and sensitivity to intercellular cues. Some of these pathways were not observed in iMGLs *in vitro*, likely reflecting the reduced complexity of monocultures. Conversely, LACTB KO xenotransplanted microglia showed a negative enrichment in protein-synthesis, oxidative phosphorylation and cell-cycle biological processes. These pathways were also altered in LACTB KD/KO/ED myeloid cells *in vitro*, although with some discrepancies in statistical enrichment directionality (Figure 3A, 3B and 3G). Interestingly, we observed a negative enrichment of lysosomal transport in this pathologic context, which aligns with the reduction in lysosomal acidification and proteolysis observed in LACTB KD/KO myeloid cells upon stimulation *in vitro* (Figure 5D).

To further characterize the *in vivo* consequences of LACTB KO in microglia, we performed immunohistochemistry followed by quantitative image analysis. We did not observe differences in the area covered by microglia (IBA1+), intensity, or average size of microglia (Supplementary Figure 5A). However, we found a statistically significant increase in the number of LACTB KO human microglia (IBA1+KU80+) around amyloid plaques compared to WT (Figure 6E). We also found a non-statistically significant decrease in the amyloid plaque area, plaque number, and average size measured by Aβ42 immunostaining (Figure 6F and Supplementary Figure 5B). When classifying the amyloid plaques by size, we observed fewer large plaques (> 40 μm) in mice xenotransplanted with LACTB KO vs WT human microglia (Supplementary Figure 5B). In line with these results, a non-statistically significant reduction in plaque load was observed using another amyloid marker (MOAB, Supplementary Figure 5C). We also double-stained the amyloid plaques with a diffuse amyloid marker (MOAB) and a dense core marker (ThioS) and observed a non-statistically significant increase in plaque compaction (higher ThioS/MOAB ratio) in hCSF1-5XFAD mice xenotransplanted with LACTB KO compared to mice xenotransplanted with WT human microglia (Figure 6G).

Finally, in order to characterize microglial activation, we immunostained with antibodies to LGALS3 and CD9. We observed a non-statistically significant increase in CD9 and LGALS3 activation markers (area covered in cortex plus hippocampal regions) in mice xenotransplanted with LACTB KO human microglia compared to WT (Figure 6H), suggesting mild alterations in microglial activation. Because, at the transcription level, we observed that LACTB KO led to alterations in pathways related to antigen-presentation (Figure 3F: iMG10, iMG12 and Figure 3G), we also immunostained for the HLA-DRB1 marker. We observed an increase in HLA microglia (area covered in the subiculum region) in mice xenotransplanted with LACTB KO human microglia microglia (Figure 6J).

In short, LACTB KO altered the metabolism and immune response of human microglia *in vivo*, which led to mild changes in activation profile and higher association with amyloid plaques.

## Discussion

Lactamase β (*LACTB*) is a mitochondrial enzyme encoded by a gene that resides within an AD risk locus on chromosome 15; however, its functional contribution to disease pathogenesis has not been investigated. Although broadly expressed, its expression is enriched in myeloid and other immune cells and our analysis using monocyte-derived macrophage eQTL data indicates that lower LACTB expression in myeloid cells is associated with reduced AD risk. Importantly, in a similar study using pQTL data, lower LACTB protein levels in the brain have also been associated with decreased AD risk [16]. Together with LACTB2, they are the only two β-lactamase genes in the human genome, and lower LACTB2 expression in monocytes has also been associated with reduced AD risk by TWAS in an independent study [19].

LACTB loss-of-function mutations in mice and humans have also been genetically linked to higher levels of a specific, poorly characterized metabolite: succinylcarnitine [28,67]. Here, we validated these findings showing that partial or total loss of LACTB protein, or loss of its enzymatic activity, is associated with higher succinylcarnitine levels across multiple *in vitro* and *in vivo* models. We also showed that LACTB hydrolyzes succinylcarnitine into carnitine and succinate, two critical metabolites at the intersection of glycolysis and lipid metabolism. Higher succinate availability may lead to an increase in APP and tau succinylation, which have been shown to promote Aβ accumulation and plaque formation, and tau aggregation to tangles [92]. This activity of LACTB may at least in part explain why its loss is associated with higher succinylcarnitine levels. Remarkably, higher succinylcarnitine levels in human CSF and brain are associated with lower AD risk [30], thereby suggesting a causal path from genetically encoded lower LACTB expression in myeloid cells to lower AD risk, possibly mediated by elevated succinylcarnitine levels. In addition, the oxoglutarate carrier SLC25A11, which is located in the inner mitochondrial membrane and can exchange succinate, has been nominated as an AD risk gene in an African American admixture GWAS [93]. On the other hand, succinylation of LACTB reduces its enzymatic activity ([11,69] and our data), and this is particularly relevant given that SUCLG2—which represses LACTB succinylation at the K288 residue [94]—is genetically linked to AD risk, cognitive decline and succinylcarnitine levels [95]. SUCLG2 is predominantly expressed in microglial cells in postmortem brains of both AD patients and age-matched controls [95].

LACTB is a mammalian homolog of bacterial β-lactamases; however, the physiological relevance of its DAEP activity in eukaryotic cells remains poorly understood. Age-related studies have documented an accumulation of D-Asp residues in long-lived proteins [4], and DAEP activity may prevent the accumulation of racemized proteins in aging [96]. More specifically, racemization of L-aspartate residues at position 7 and/or 23 of Aβ to D-Asp alters aggregation kinetics of amyloid fibrils [5]; and an accumulation of D-Asp residues has been detected in tau proteins forming paired helical filaments [6]. In addition, tau can be cleaved at D421 by caspases, promoting aggregation [97], and the recessive AD-associated APP mutation A673V shifts the preferential β-cleavage site of APP by BACE1 from the glutamate residue at position 11 of Aβ to the aspartate residue at position 1 (D672 of APP), resulting in higher C99 generation (and subsequent Aβ production) [98]. These functionally relevant aspartate sites in disease-relevant proteins like APP, Aβ and tau may also represent potential targets of LACTB serine hydrolase activity. In fact, LACTB has been identified as the first mammalian protein with intrinsic DAEP activity ([3] and our data) and is able to directly cleave a 10-residue peptide corresponding to Aβ1–10 containing D-Asp at position 7 [3]. Further studies are required to explore the ability of LACTB to modify APP processing or Aβ and tau aggregation *in vitro* and *in vivo*.

Through transcriptomics and functional assays, we showed that LACTB alters the metabolism of myeloid cells, leading to an increase in oxidative phosphorylation and a reduction in triacylglycerides. Boosting TREM2 signaling (known to be protective against AD) with an activating antibody shows a similar phenotype in iPSC-derived microglia [99]. In addition, we observed that LACTB KD/KO leads to a reduction of protein synthesis in myeloid cells. Notably, LACTB co-expression module genes are enriched in this microglial sub-population, which is characterized by the expression of ribosomal processing and protein synthesis-related genes. Interestingly, the subunit 28 ribosomal RNA expression is altered in CA1 and dental gyrus in AD patients [100] and de novo proteome is disturbed in young APP/PS1 mice prior to symptom onset [101]. More specifically, a repression in translation has been described upon microglia LPS-activation *in vitro* [102]. In this regard, LACTB KO human xenotransplanted microglia showed an increase in activation markers compared to WT. By shifting energy away from bulk protein synthesis and lipid storage, LACTB reduction may promote a bioenergetically efficient state—characterized by high OXPHOS—that supports microglial function and counters the metabolic exhaustion seen in AD [103].

IFN and TNF response-related clusters have been identified in several mouse and human single-cell microglia studies [104–107]. Our data showed increased LACTB levels and decreased succinylcarnitine levels in THP1 macrophages and iPSC-derived microglia stimulated with IFN-β, IFN-γ, or TNF-α *in vitro*. In addition, LACTB KO/KD resulted in altered expression of genes involved in the IFN/TNF response. After IFN/TNF stimulation, myeloid cells reduced myelin phagocytosis and lysosomal activity. Similarly, lower PU1 levels, associated with AD protection, reduced myelin phagocytosis [91] and THP1 macrophages polarized towards a lipid disease-associated state also showed lower phagocytic and lysosomal activity [90]. Lower microglia efferocytosis activity in pro-inflammatory conditions could prevent microglial cell exhaustion and excessive synaptic pruning.

Finally, in a mouse model of amyloid deposition, LACTB KO in human microglia did not markedly alter amyloid plaque density, similar to what has been observed in KO experiments with other myeloid AD risk genes such as TREM2 and PLCG2 [59,108]. However, opposite to what has been observed with TREM2 and PLCG2 KO, LACTB KO in human microglia led to increased microglia-plaque association and activation measured by an antigen presentation marker. In this regard, the P522R gain-of-function and protective variant of PLCG2 has also been shown to promote the expression of antigen presentation genes by human microglia in 5XFAD-hCSF1 xenotransplanted mice [59]. Overall, these effects are directionally consistent with human genetics and functional genomics evidence linking TREM2 loss-of-function with increased AD risk, and PLCG2 gain-of-function or LACTB loss-of-function with decreased AD risk. In the future, we will also investigate the effect of LACTB KO in mouse models of tauopathy, encouraged by the fact that succinylcarnitine levels have been shown to correlate with phospho-Tau (181P) in CSF [109].

Together these data implicate a novel gene (the mitochondrial serine hydrolase LACTB) and metabolite (succinylcarnitine) pair, in myeloid cells like microglia and macrophages, in the pathogenesis of AD. In contrast to many AD risk genes, LACTB is structurally and enzymatically characterized, and small-molecule inhibitors of its bacterial orthologs are already in clinical use to treat β-lactam–resistant infections, highlighting its potential druggability. In addition, LACTB DAEP activity can be quantified using a scalable fluorometric assay (Figure 2A), enabling high-throughput screening of small molecule inhibitors. Importantly, succinylcarnitine represents a proximal, mechanistically-linked readout of LACTB activity that can be measured in cell culture, mouse brain, and human biofluids (blood, CSF, or urine). The availability of a readily quantifiable causal endophenotype and pharmacodynamic biomarker of target engagement substantially strengthens the translational tractability of LACTB inhibition.

## Conclusions

Our study identifies a novel axis linking reduced LACTB expression to increased succinylcarnitine levels in myeloid cells, which in turn may lower AD risk. Succinylcarnitine can be directly cleaved by LACTB, while succinylation reduces LACTB activity, suggesting a regulatory negative feedback loop. In myeloid cells, reduced LACTB alters cellular metabolism and modulates efferocytosis following IFN/TNF stimulation. Consistently, LACTB KO human microglia xenotransplanted into the brain of mice with amyloid pathology show increased association with amyloid plaques, a phenotype linked to AD protection. LACTB inhibition represents a promising therapeutic strategy for AD, supported by human genetics and the potential use of succinylcarnitine as a causal endophenotype and pharmacodynamic biomarker.

## Acknowledgements

We would like to acknowledge the Stem Cell Core at Mount Sinai (NY, USA) for the generation of the WTC11 LACTB CRISPR-edited iPSC lines (directed by Samuelle Marro). We also would like to acknowledge Tiago Olivera (University of Minho, Portugal) for his advice in interpreting lipidomic data, Isabelle Gerin for her advice on LACTB DAEP activity (UCLouvain University, Belgium) and Ciyang Wang (Washington University, USA) for support regarding succinylcarnitine data in humans. Metabolomic data were generated in part through the use of the MDACC Metabolomics Core Facility, which received partial support from the National Cancer Institute under grant P30CA016672 to MD Anderson Cancer Center. The research reported in this publication was not directly funded through the P30CA016672 grant to MD Anderson Cancer Center and is not within the scope of such grant.

This work was supported in part through the computational and data resources and staff expertise provided by Scientific Computing and Data at the Icahn School of Medicine at Mount Sinai and supported by the Clinical and Translational Science Awards (CTSA) grant UL1TR004419 from the National Center for Advancing Translational Sciences. Research reported in this publication was also supported by the Office of Research Infrastructure of the National Institutes of Health under award number S10OD026880 and S10OD030463. The content is solely the responsibility of the authors and does not necessarily represent the official views of the National Institutes of Health.

## List of abbreviations

AD: Alzheimer’s disease
AUC: area under the curve
BMDMs: bone marrow-derived macrophages
CE: cholesterol ester Cer ceramides
CNS: central nervous system
CSF: cerebrospinal fluid
DAEP: D-aspartyl endopeptidase activity
DEGA: differential gene expression analysis
DMSO: dimethyl Sulfoxide
DREAM: differential expression analysis for repeated measures
dhCer: dihydroceramides
ED: enzymatically-dead
GSEA: gene set enrichment analysis
GWAS: gene wide association study
HPCs: hematopoietic progenitor cells
ICMS: ion chromatography-mass spectometry
IFN: interferon
IHC: immunohistochemistry
IP: immunoprecipitation
iPSC: induced pluripotent stem cell
iMGLs: iPSC-derived microglia
KD: knock-down
KO: knock-out
LACTB: lactamase B
LACTB2: lactamase B2
LacCer: lactosylceramide
LPS: lipopolysaccharide
MCSF: macrophage colony-stimulating factor
MG/DG/TG: mono/di/tri-acylglycerides
MP: mobile phase
MR: mendelian randomization
NES: normalized enrichment score
OXPHOS: oxidative phosphorylation
PMA: Phorbol 12-myristate 13-acetate
QTL: quantitative trait locus
RT: room temperature
SCR: scrambled
TNF: tumor necrosis factor
TWAS: transcriptome wide association studies
WT: wild-type

## Declarations

### Ethics approval and consent to participate

Not applicable

### Consent for publication

Not applicable

### Availability of data and materials

The data generated in this manuscript has been deposited in GEO: bulk RNAseq in THP1 macrophages (GSE323383), WTC11 iMGLs (GSE323384), mouse isolated microglia (GSE324607), human xenotransplanted microglia (GSE324508) and single-cell RNAseq in WTC11 iMGLs (GSE324494). Fastq files of external datasets analyzed in this manuscript are available to download from NCBI GEO for Mancuso et al., (GSE216999), Tuddenham et al., (GSE204702) and Smith et al., (GSE160936), and from the Synapse portal for Olah et al., (syn21438358), Prater et al., (syn51272688 through a data access request) and Lee et al., (syn52795287).

### Authors’ contributions

Conceptualization, study design, and methodology: C.R-M., A.M.G., E.M., R.G-G, J.R; data collection, analysis, and visualization: C.R-M., R.G-G., W.Y.S..; MR analysis: E.M.; bioinformatic support: T.P., M.S., immunoprecipitation experiments: K.A., mouse brain cell isolation: O.C., mouse work support: M.R., Y.L.L., succinylcarnitine measurements: Q.X., xenotransplantation of human cells into the mouse brain: H.D, M.B-J.; enzymatic activity advice: G.B., succinylcarnitine dataset: C.C., writing of original draft: C.R-M., revising: R.G-G., J.R., A.M.G., E.M. All authors read and approved of the final manuscript.

---

Supplementary Figure 1. A) LACTB expression is higher in microglia compared to hematopoietic precursors (HPCs) or induced-pluripotent cells (iPSC). Data from Stemformatics (Abud et al., 2017 dataset [65]). B) Experimental design for the differentiation of THP1 monocytes to macrophages, combined with the small-interfering RNA (siRNA) treatment. We observed a 90% and 50% LACTB reduction at mRNA (C) and protein (D) levels, respectively. Graphs display individual data points (left) alongside estimated marginal means with 95% confidence intervals (right). For the raw data, dot shapes represent independent macrophage differentiations. E) Schematic diagram of the differentiation of induced pluripotent stem cells (iPSC) to microglia (iMGLs). F) Validation of LACTB knock-out iPSC-derived microglia (iMGLs) from WTC11 donor (3 clones per genotype). G) LACTB expression levels in isolated microglia, astrocytes and hippocampal lysate in control and LACTB enzymatically dead (ED) mice. H) LACTB protein levels, assessed by an antibody targeting the active site, in WT and LACTB ED mice (n=2). Statistical details are provided in Table 1.

Supplementary Figure 2. A) Heatmap of Pearson coefficients between LACTB expression and leading-edge genes identified in THP1 KD and iMGL KO bulk-RNAseq GSEA across clusters in re-analyzed publicly-available datasets [70–75], which were consistently positively- (mean r > 0.1) or negatively-correlated (r mean < −0.1) across studies. B) mRNA expression of differentiation markers in control and LACTB KD THP1 macrophages (n = 4) and LACTB KO WTC11 microglia (n=3 independent clones, 2 independent differentiations per clone). Graphs show normalized (adjusted) transcript counts. C) Cell proliferation (EdU assay) in control and LACTB KD THP1 macrophages (n=3) and LACTB KO WTC11 microglia (n=3 clones, 2 independent differentiations per clone). Graphs display individual data points (left) alongside estimated marginal means with 95% confidence intervals (right). For the raw data, dot shapes represent independent macrophage differentiations, and dot colors indicate distinct microglia clones. Statistical details are provided in Table 1.

Supplementary Figure 3. A) Mitochondrial membrane potential in LACTB KD/KO myeloid cells compared to SCR/WT. B) Representative electron microscopy images of LACTB KO iMGLs compared to WT. C) UMAP showing module scores from ARACNe-inferred LACTB co-expression module across iMGLs, with highest enrichment in iMG7, a ribosomal processing-associated cluster. D-E) Lipid droplet accumulation (D) and cholesterol efflux (E) in LACTB KD/KO myeloid cells compared to SCR/WT. Graphs display individual data points (left) alongside estimated marginal means with 95% confidence intervals (right). For the raw data, dot shapes represent independent macrophage differentiations, and dot colors indicate distinct microglia clones. Statistical details are provided in Table 1.

Supplementary Figure 4. A) LACTB protein levels in THP1 macrophages or WTC11 iMGLs stimulated with IFN-β, IFN-γ, or TNF-α for 24 hours. B) mRNA expression (log2FC) in control and LACTB KD/KO myeloid cells at baseline or after stimulation with IFN-β, IFN-γ, or TNF-α (THP-1 macrophages, for 6 and 24h hours n = 3; WTC11 iMGLs, for 24 hours, n=3 clones, 2 differentiations each). C) Levels of cytokines in control and LACTB KD/KO myeloid cells at baseline or following stimulation with IFN-β, IFN-γ, or TNF-α for 24 hours (THP-1 macrophages, n=2 independent differentiations; WTC11 iMGLs, n=3 clones). Statistical details are provided in Table 1.

Supplementary Figure 5. Immunohistochemistry analysis of (A) Percentage of area covered, integrated density and average size of microglia (Iba1+ cells), (B) Number of small (0-20 μm), medium (20-40 μm) and large (> 40 μm) amyloid plaques (n=2 clones per genotype, 4 mice injected per clone). Graphs display individual data points (left) alongside estimated marginal means with 95% confidence intervals (right). For the raw data, dot colors indicate distinct microglia clones. Statistical details are provided in Table 1.

