## Supplementary figures and images for "Reduced LACTB expression in myeloid cells is associated with elevated succinylcarnitine levels and reduced Alzheimer’s disease risk"

### Additional file 2

#### 230314_2463_Korg_Targeting sequence

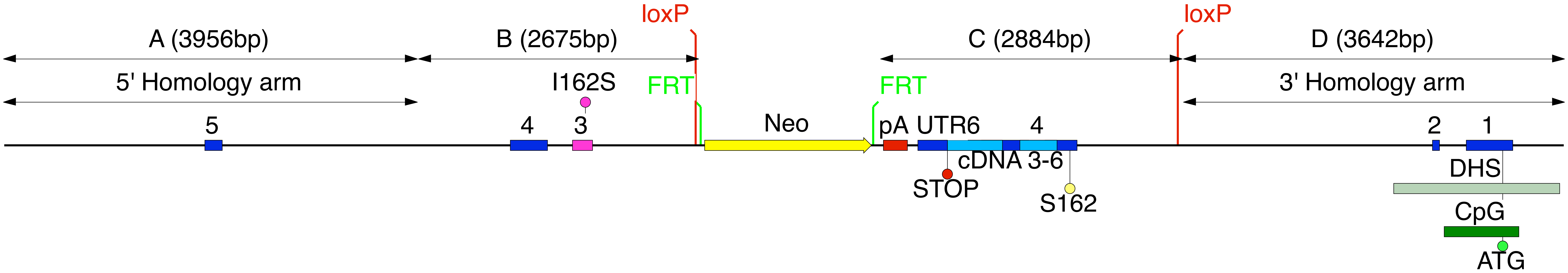

### Additional file 3

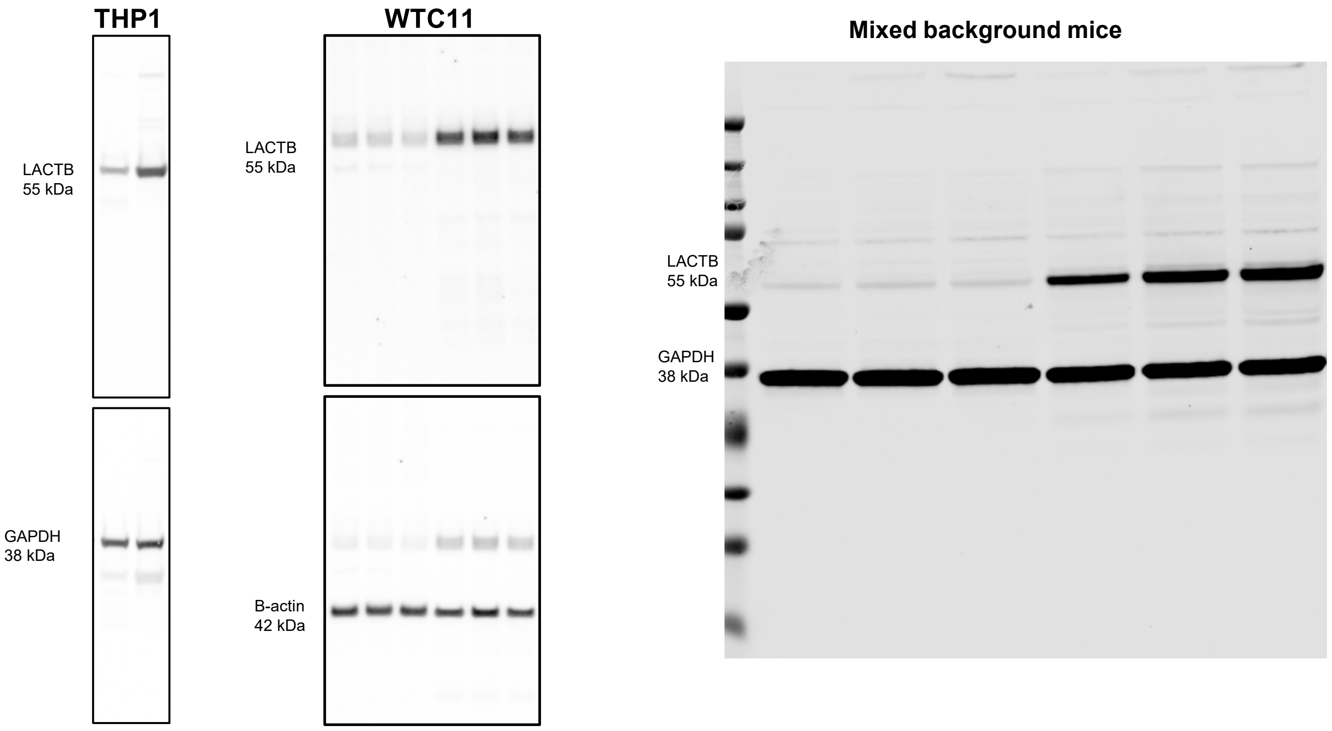


Western-blots in Figure 1

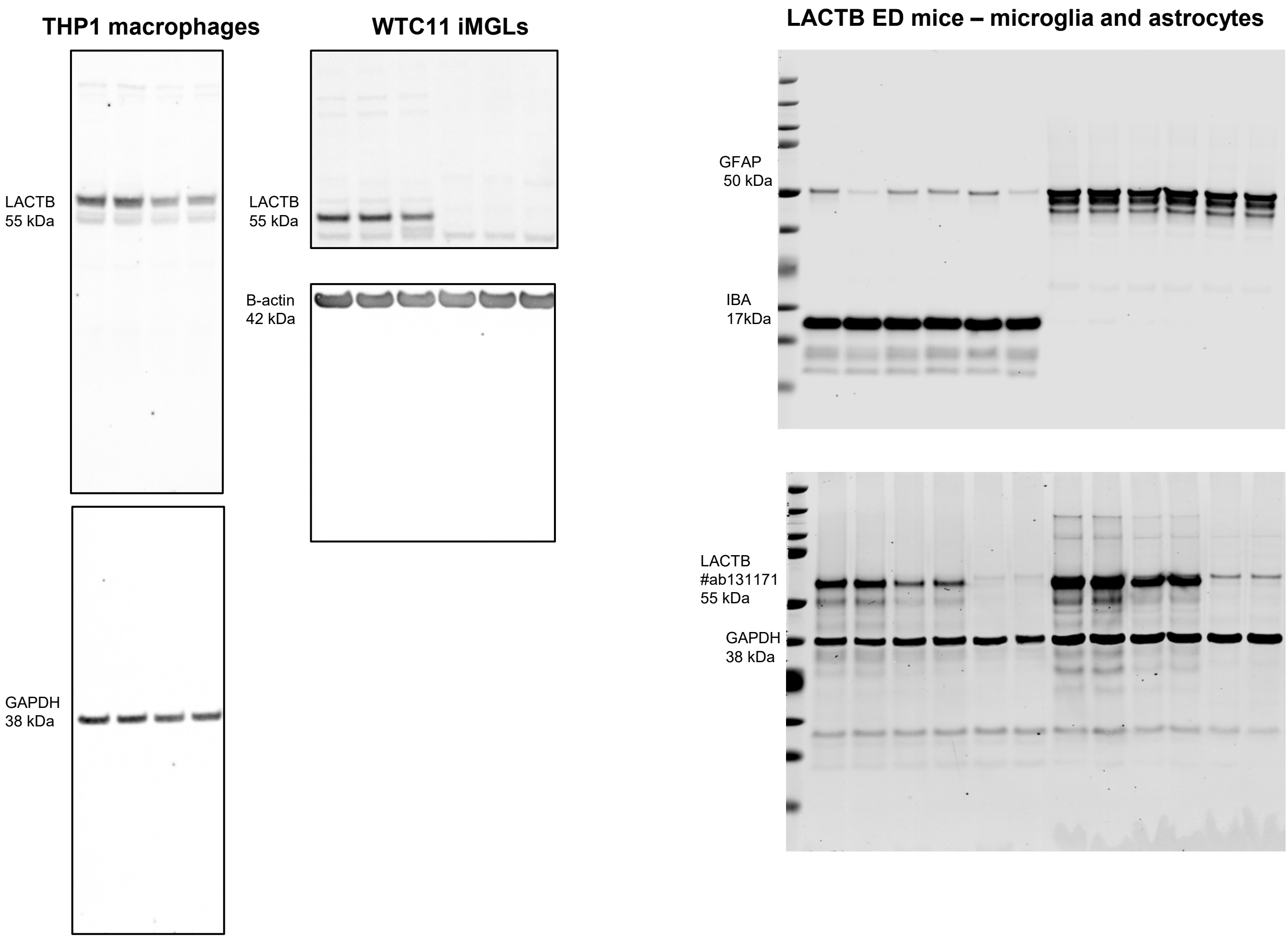


Western-blots in Supplementary Figure 1

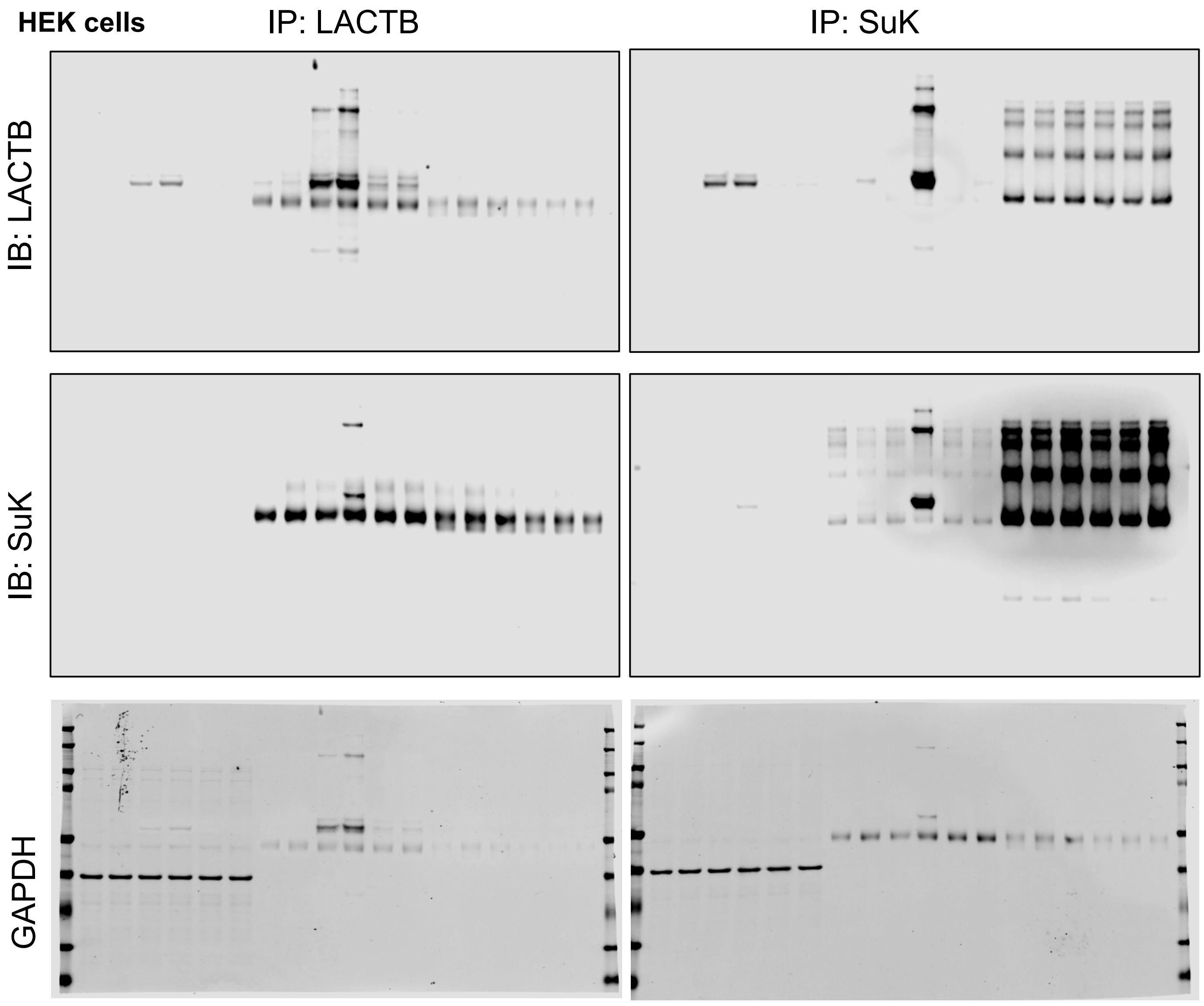


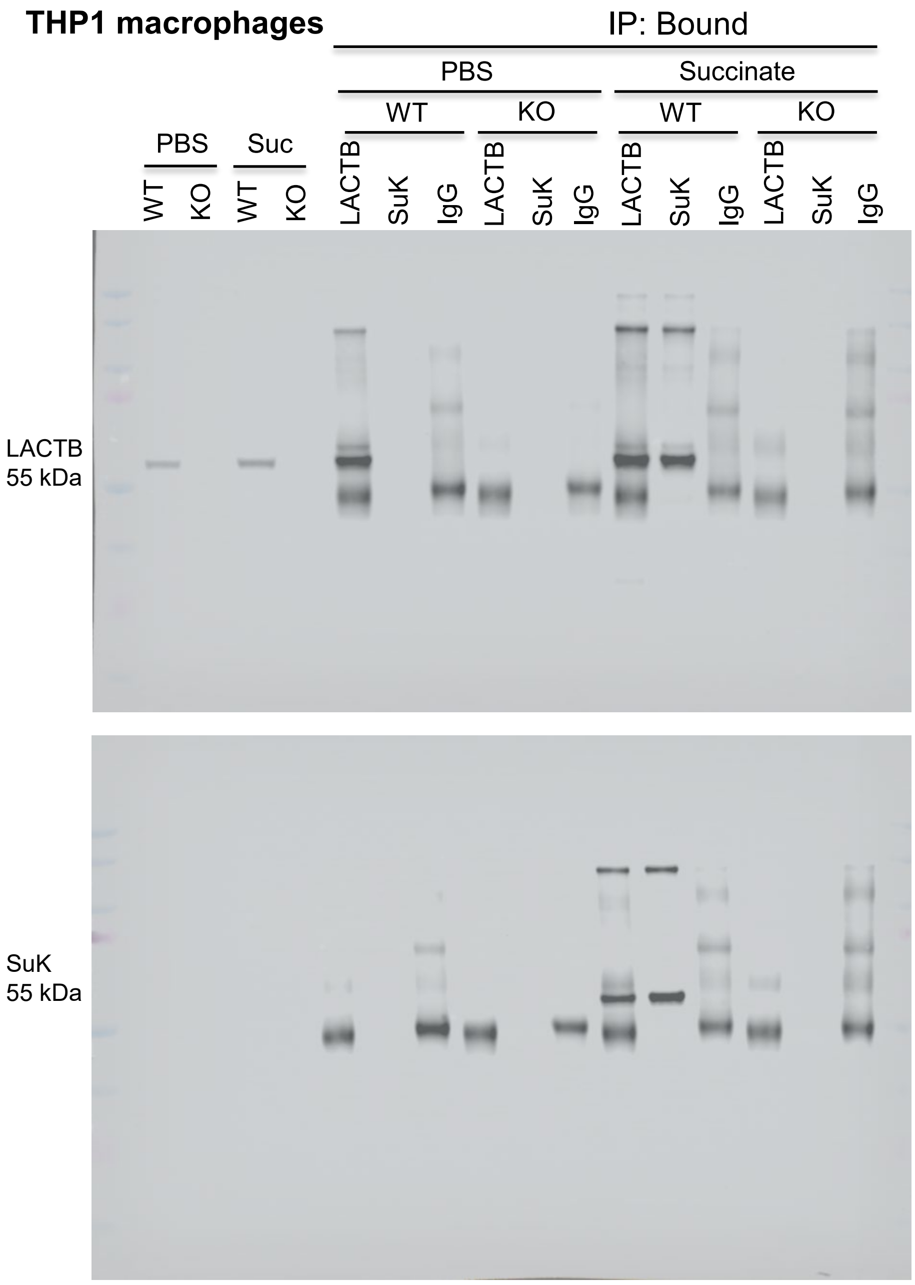


Western-blots in Figure 2

### Supl Figure 1

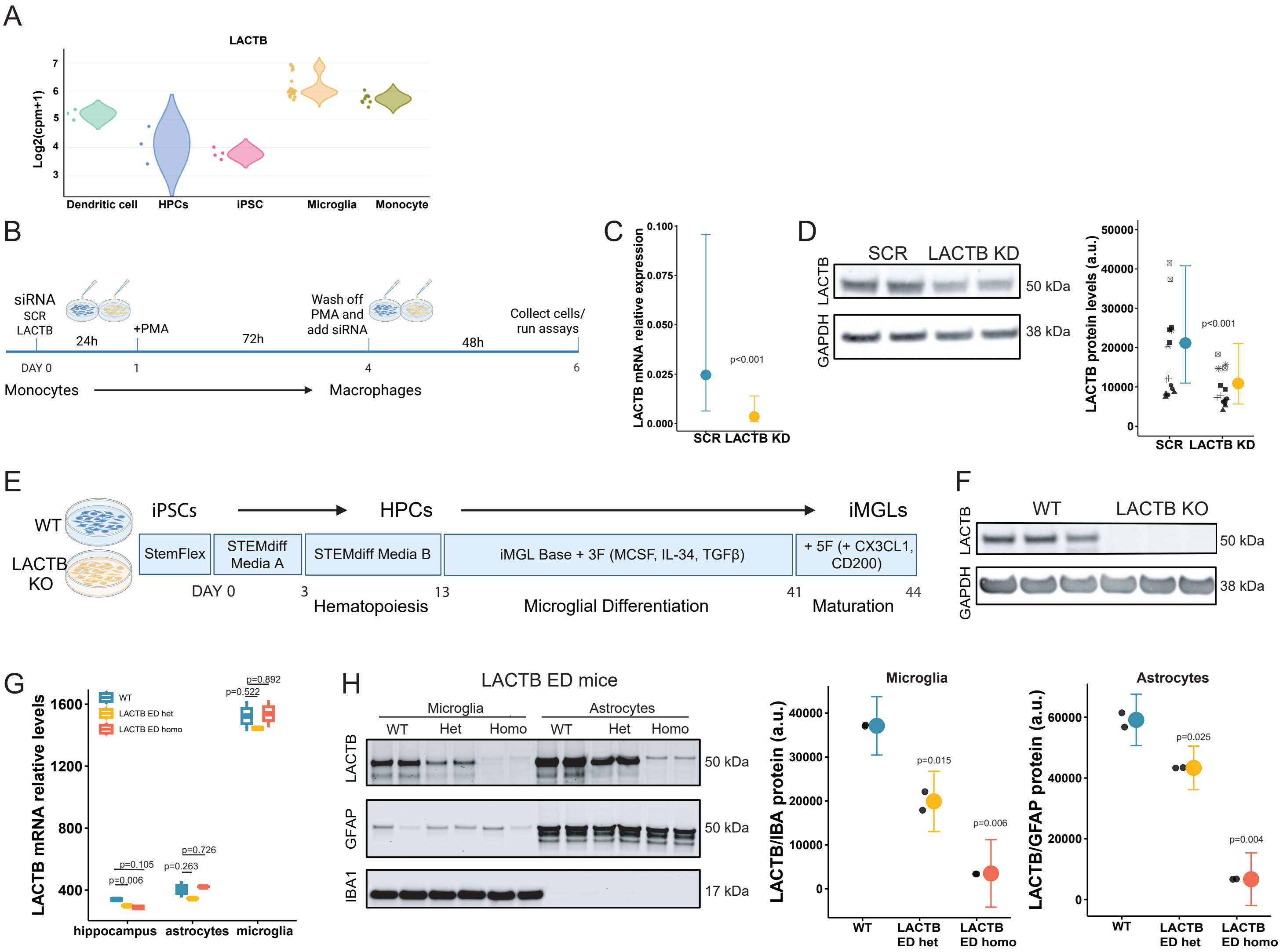

### Supl Figure 2

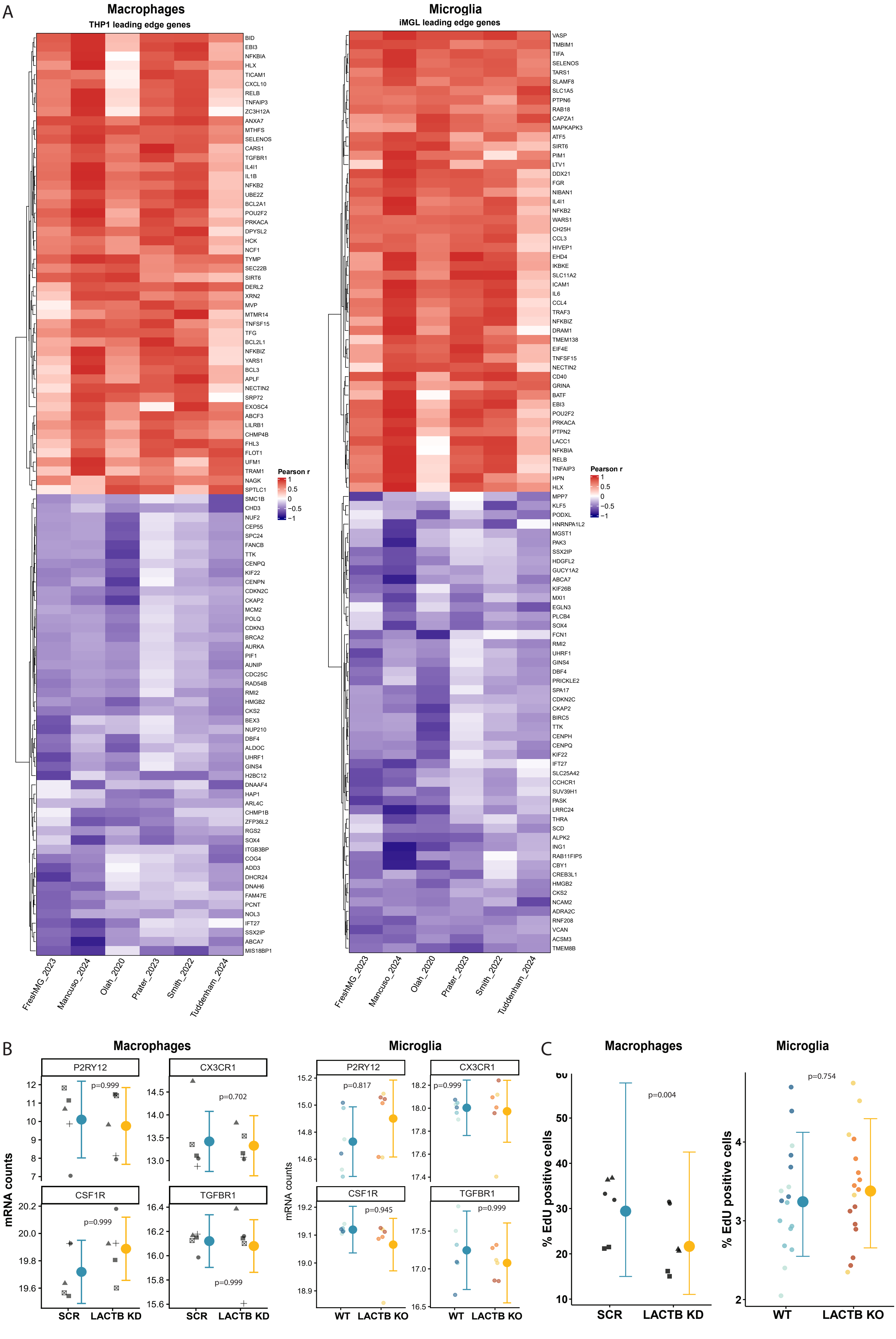

### Supl Figure 3

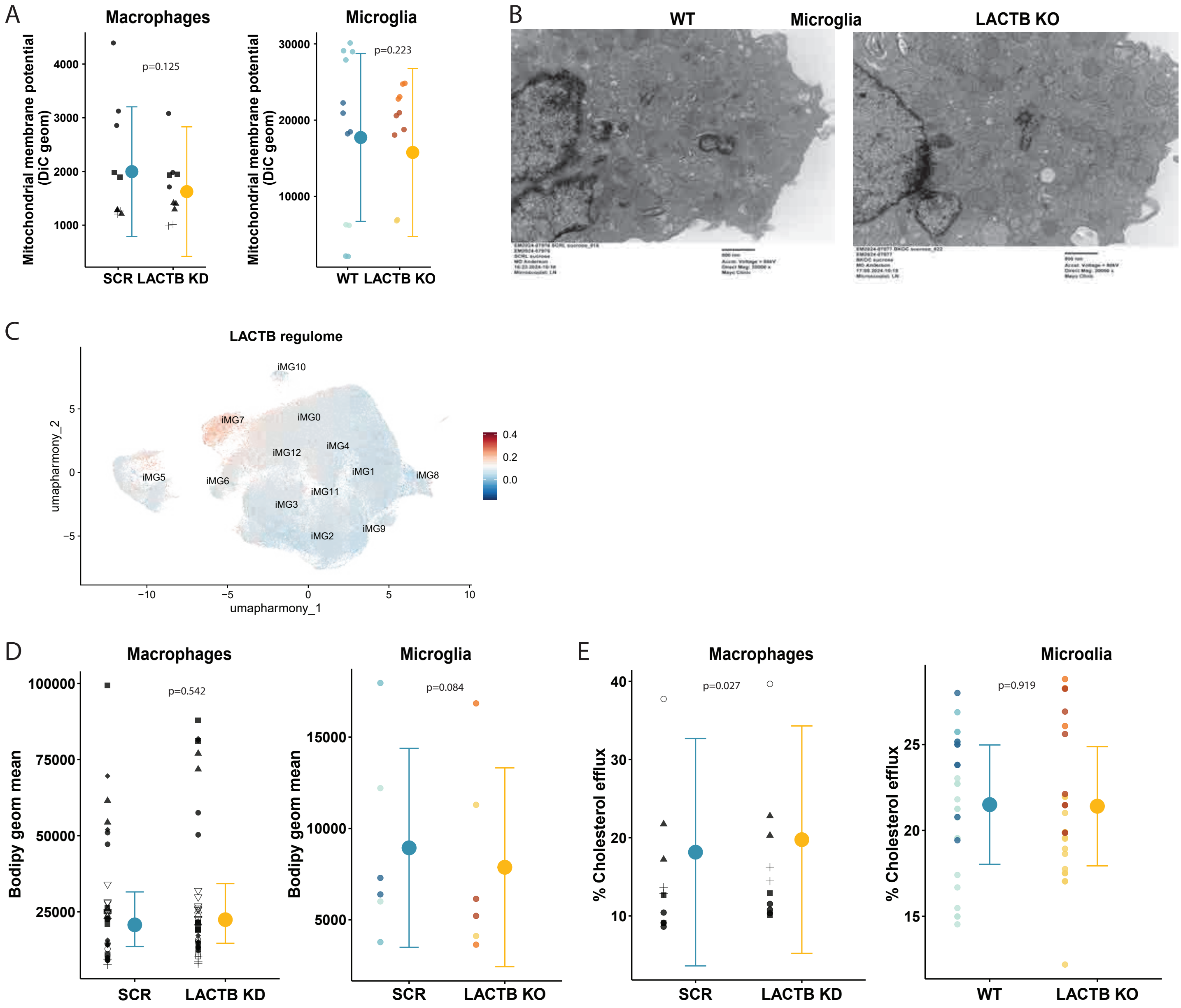

### Supl Figure 4

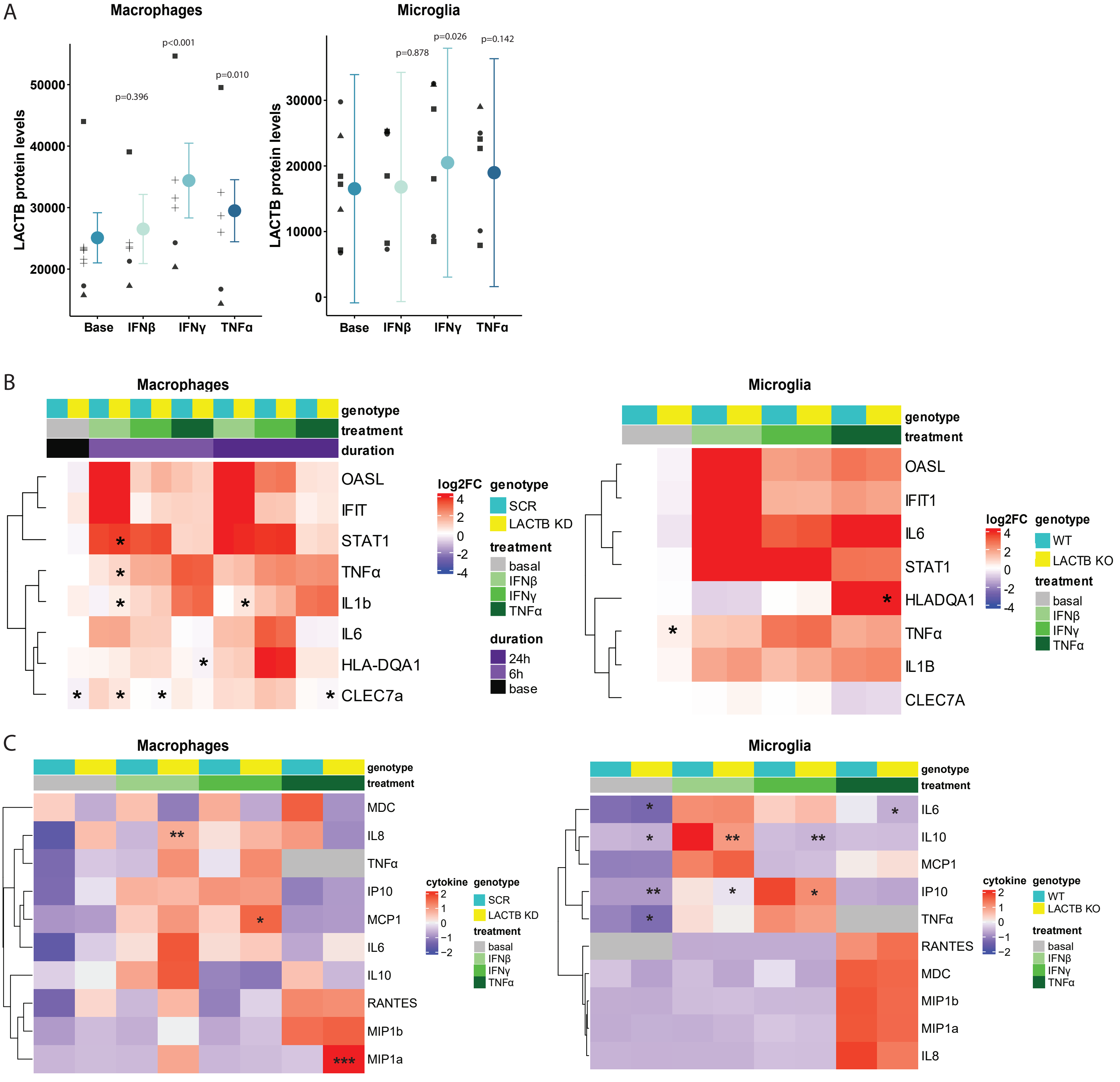

### Supl Figure 5

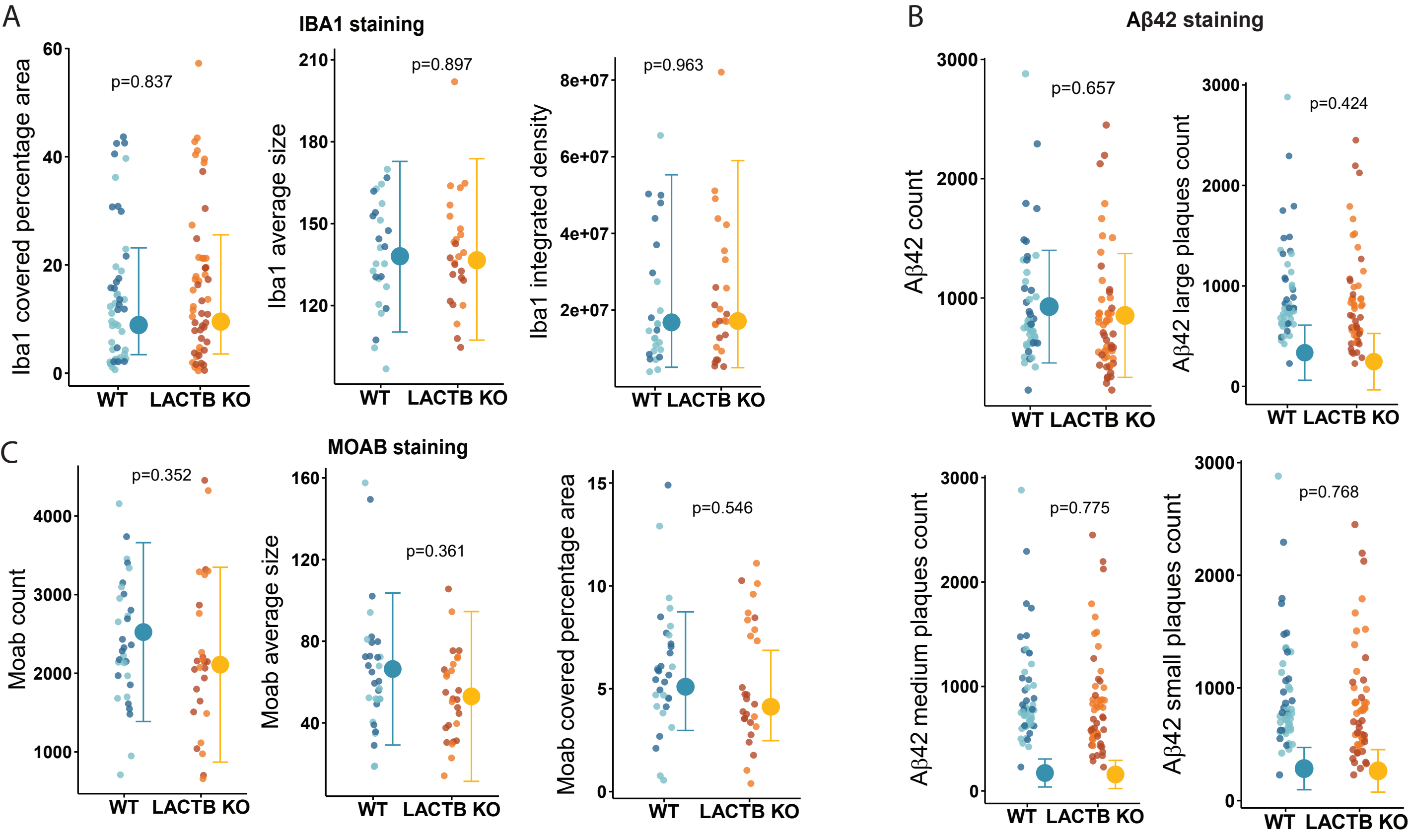
