## Additional file 1 for "Reduced LACTB expression in myeloid cells is associated with elevated succinylcarnitine levels and reduced Alzheimer’s disease risk"

### TwoSampleMR

Edoardo “Dado” Marcora

2022-11-23

#### Setup environment

```
# install.packages("TwoSampleMR", repos = c("https://mrcieu.r-universe.dev", "https://cloud.r
# ip("MRCIEU/TwoSampleMR")
library(TwoSampleMR)
library(EnsDb.Hsapiens.v75)
library(locuszoomr)
library(tidyverse)
library(magrittr)
library(patchwork)

theme_set(theme_bw(base_size = 8))

set.seed(666)

load("rf.rdata")
```

#### Read and prep data

```
eqtl = read.delim("data/sumstats/2022-10-12.STARNET.eQTL.MP.chr15.CPRA_b37.tsv.gz", comment.c

mqtl = read.delim("data/sumstats/Cruchaga2022.mQTL.CSF.X100001948.chr15.CPRA_b37.tsv.gz", com

gwas = read.delim("data/sumstats/Bellenguez2022.chr15.CPRA_b37.tsv.gz", comment.char = "#") %
```

#### LACTB

```
GENE = "ENSG00000103642" # LACTB

eqtl.gene = eqtl %>% filter(TRAIT == GENE)

mqtl.gene = mqtl %>% filter(ID %in% eqtl.gene$ID)
gwas.gene = gwas %>% filter(ID %in% eqtl.gene$ID)
```

##### Format and harmonise data for MR

```
eqtl.gene.ld_clump = ieugwasr::ld_clump(eqtl.gene %>% select(rsid = ID, pval = P), bfile = "d
```

```
Clumping gVL5KT, 6600 variants, using: data/ldref/1000G.EUR.CPRA_b38.chr15

Removing 6585 of 6600 variants due to LD with other variants or absence from LD reference pan
```

```
eqtl.gene.exposure = format_data(eqtl.gene %>%
                                 filter(ID %in% eqtl.gene.ld_clump$rsid) %>%
                                 select(SNP = ID,
                                        beta = BETA,
                                        se = SE,
                                        effect_allele = ALT,
                                        other_allele = REF,
                                        eaf = AF,
                                        Phenotype = TRAIT,
                                        chr = CHROM,
                                        position = POS,
                                        sample_size = N,
                                        pval = P,
                                        gene = TRAIT),
                                 type="exposure")
```

```
mqtl.gene.ld_clump = ieugwasr::ld_clump(mqtl.gene %>% select(rsid = ID, pval = P), bfile = "d
```

```
Clumping 7SxXqj, 6213 variants, using: data/ldref/1000G.EUR.CPRA_b38.chr15

Removing 6204 of 6213 variants due to LD with other variants or absence from LD reference pan
```

```
mqtl.gene.exposure = format_data(mqtl.gene %>%
                                 filter(ID %in% mqtl.gene.ld_clump$rsid) %>%
                                 select(SNP = ID,
                                        beta = BETA,
                                        se = SE,
                                        effect_allele = ALT,
                                        other_allele = REF,
                                        eaf = AF,
                                        Phenotype = TRAIT,
                                        chr = CHROM,
                                        position = POS,
                                        sample_size = N,
                                        pval = P),
                                 type="exposure")
```

```
mqtl.gene.outcome = format_data(mqtl.gene %>%
                                select(SNP = ID,
                                       beta = BETA,
                                       se = SE,
                                       effect_allele = ALT,
                                       other_allele = REF,
                                       eaf = AF,
                                       Phenotype = TRAIT,
                                       chr = CHROM,
                                       position = POS,
                                       sample_size = N,
                                       pval = P),
                                type="outcome")
```

```
gwas.gene.outcome = format_data(gwas.gene %>%
                                select(SNP = ID,
                                       beta = BETA,
                                       se = SE,
                                       effect_allele = ALT,
                                       other_allele = REF,
                                       eaf = AF,
                                       Phenotype = TRAIT,
                                       chr = CHROM,
                                       position = POS,
                                       sample_size = N,
                                       ncase = N_CASES,
                                       ncontrol = N_CTRLS,
```

```
                          pval = P,
                          gene = TRAIT),
                     type="outcome")
```

```
Generating sample size from ncase and ncontrol
```

```
dat = harmonise_data(exposure_dat = eqtl.gene.exposure,
                     outcome_dat = gwas.gene.outcome)
```

```
Harmonising ENSG00000103642 (E3aY79) and AD (5rtOEu)
```

###### Analyse data and print/plot results

```
(res = mr(dat))
```

```
Analysing 'E3aY79' on '5rtOEu'
```

```
  id.exposure id.outcome outcome        exposure                    method nsnp
1      E3aY79     5rtOEu      AD ENSG00000103642                  MR Egger   15
2      E3aY79     5rtOEu      AD ENSG00000103642           Weighted median   15
3      E3aY79     5rtOEu      AD ENSG00000103642 Inverse variance weighted   15
4      E3aY79     5rtOEu      AD ENSG00000103642               Simple mode   15
5      E3aY79     5rtOEu      AD ENSG00000103642             Weighted mode   15
          b       se      pval
1  0.007503 0.002400 0.0080258
2  0.005276 0.001288 0.0000419
3  0.002863 0.001541 0.0631032
4 -0.003801 0.004243 0.3854475
5  0.004797 0.001743 0.0156014
```

```
mr_heterogeneity(dat)
```

```
  id.exposure id.outcome outcome        exposure                    method    Q
1      E3aY79     5rtOEu      AD ENSG00000103642                  MR Egger 19.1
2      E3aY79     5rtOEu      AD ENSG00000103642 Inverse variance weighted 27.1
  Q_df  Q_pval
1   13 0.11999
2   14 0.01871
```

```r
mr_pleiotropy_test(dat)
```

```
  id.exposure id.outcome outcome        exposure egger_intercept      se
1      E3aY79     5rt0Eu      AD ENSG00000103642        -0.02274 0.00975
     pval
1 0.03639
```

```r
mr_leaveoneout(dat)
```

```
          exposure outcome id.exposure id.outcome samplesize              SNP
1  ENSG00000103642      AD      E3aY79     5rt0Eu     485999 15:62208259:g:a
2  ENSG00000103642      AD      E3aY79     5rt0Eu     485999 15:62218106:a:t
3  ENSG00000103642      AD      E3aY79     5rt0Eu     485999 15:62221882:g:a
4  ENSG00000103642      AD      E3aY79     5rt0Eu     485999 15:62268638:t:c
5  ENSG00000103642      AD      E3aY79     5rt0Eu     485999 15:62429972:g:c
6  ENSG00000103642      AD      E3aY79     5rt0Eu     485999 15:62520456:t:g
7  ENSG00000103642      AD      E3aY79     5rt0Eu     485999 15:62648840:c:t
8  ENSG00000103642      AD      E3aY79     5rt0Eu     485999 15:62676645:c:t
9  ENSG00000103642      AD      E3aY79     5rt0Eu     485999 15:62768090:t:c
10 ENSG00000103642      AD      E3aY79     5rt0Eu     485999 15:62930647:c:t
11 ENSG00000103642      AD      E3aY79     5rt0Eu     485999 15:62979292:g:c
12 ENSG00000103642      AD      E3aY79     5rt0Eu     485999 15:63141567:g:a
13 ENSG00000103642      AD      E3aY79     5rt0Eu     485999 15:63353203:c:a
14 ENSG00000103642      AD      E3aY79     5rt0Eu     485999 15:63863320:g:a
15 ENSG00000103642      AD      E3aY79     5rt0Eu     485999 15:63902855:g:a
16 ENSG00000103642      AD      E3aY79     5rt0Eu     485999              All
           b       se       p
1   0.002925 0.001585 0.06492
2   0.002930 0.001543 0.05765
3   0.002943 0.001592 0.06449
4   0.003010 0.001488 0.04312
5   0.002956 0.001550 0.05657
6   0.002953 0.001604 0.06568
7   0.002859 0.001605 0.07489
8   0.002950 0.001573 0.06072
9   0.003600 0.001433 0.01199
10  0.002869 0.001566 0.06699
11  0.002765 0.001637 0.09123
12 -0.004302 0.002118 0.04223
13  0.003019 0.001597 0.05868
14  0.003534 0.001508 0.01906
```

```
15  0.002826 0.001589 0.07532
16  0.002863 0.001541 0.06310
```

```
p1 = mr_scatter_plot(res, dat)[[1]] +
  xlab(expression(paste("SNP effect on M", Phi, " LACTB mRNA levels"))) +
  ylab("SNP effect on AD risk")

p1
```

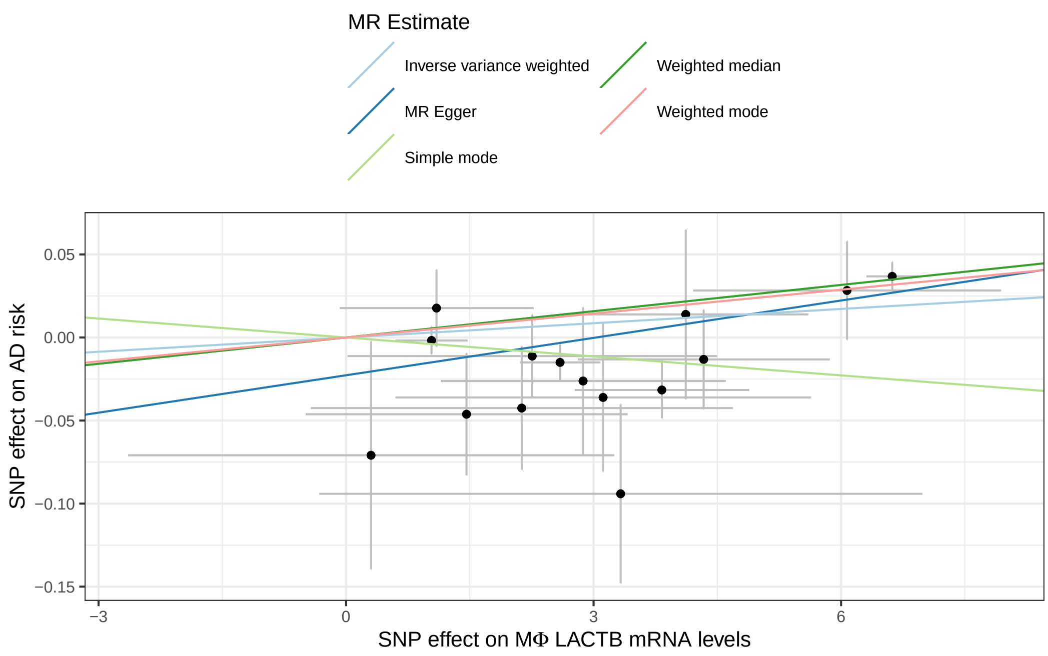


```
ggsave(str_interp("${res$exposure[1]}.${res$outcome[1]}.pdf"), plot = p1, width = 10.5, heig
```

```
dat = harmonise_data(exposure_dat = eqtl.gene.exposure,
                     outcome_dat = mqtl.gene.outcome)
```

```
Harmonising ENSG00000103642 (E3aY79) and CSF_X100001948 (vOF42P)
```

```
(res = mr(dat))
```

```
Analysing 'E3aY79' on 'vOF42P'
```

```
  id.exposure id.outcome        outcome        exposure
1      E3aY79     vOF42P CSF_X100001948 ENSG00000103642
2      E3aY79     vOF42P CSF_X100001948 ENSG00000103642
3      E3aY79     vOF42P CSF_X100001948 ENSG00000103642
4      E3aY79     vOF42P CSF_X100001948 ENSG00000103642
5      E3aY79     vOF42P CSF_X100001948 ENSG00000103642
                     method nsnp         b        se                       pval
1                  MR Egger   13 -0.008821 0.0011344 0.000008548731961660662186
2           Weighted median   13 -0.006950 0.0007417 0.0000000000000000000007246
3 Inverse variance weighted   13 -0.006062 0.0007953 0.000000000000024924979491
4               Simple mode   13 -0.002544 0.0016503 0.149119095292512604533641
5             Weighted mode   13 -0.007523 0.0006328 0.000000053670470060982310
```

```r
mr_heterogeneity(dat)
```

```
  id.exposure id.outcome        outcome        exposure
1      E3aY79     vOF42P CSF_X100001948 ENSG00000103642
2      E3aY79     vOF42P CSF_X100001948 ENSG00000103642
                     method     Q Q_df      Q_pval
1                  MR Egger 25.96   11 0.006571402
2 Inverse variance weighted 45.98   12 0.000006983
```

```r
mr_pleiotropy_test(dat)
```

```
  id.exposure id.outcome        outcome        exposure egger_intercept
1      E3aY79     vOF42P CSF_X100001948 ENSG00000103642          0.0139
        se    pval
1 0.004773 0.01413
```

```r
mr_leaveoneout(dat)
```

```
         exposure        outcome id.exposure id.outcome samplesize
1  ENSG00000103642 CSF_X100001948      E3aY79     vOF42P         NA
2  ENSG00000103642 CSF_X100001948      E3aY79     vOF42P         NA
3  ENSG00000103642 CSF_X100001948      E3aY79     vOF42P         NA
4  ENSG00000103642 CSF_X100001948      E3aY79     vOF42P         NA
5  ENSG00000103642 CSF_X100001948      E3aY79     vOF42P         NA
6  ENSG00000103642 CSF_X100001948      E3aY79     vOF42P         NA
7  ENSG00000103642 CSF_X100001948      E3aY79     vOF42P         NA
8  ENSG00000103642 CSF_X100001948      E3aY79     vOF42P         NA
```

```
9  ENSG00000103642 CSF_X100001948      E3aY79     vOF42P         NA
10 ENSG00000103642 CSF_X100001948      E3aY79     vOF42P         NA
11 ENSG00000103642 CSF_X100001948      E3aY79     vOF42P         NA
12 ENSG00000103642 CSF_X100001948      E3aY79     vOF42P         NA
13 ENSG00000103642 CSF_X100001948      E3aY79     vOF42P         NA
14 ENSG00000103642 CSF_X100001948      E3aY79     vOF42P         NA
               SNP         b        se                        p
1  15:62208259:g:a -0.006062 0.0008324 0.00000000000032815606033
2  15:62218106:a:t -0.006082 0.0008199 0.00000000000011910055833
3  15:62221882:g:a -0.006079 0.0008325 0.00000000000028288758719
4  15:62268638:t:c -0.006068 0.0008318 0.00000000000029868612309
5  15:62429972:g:c -0.006094 0.0008149 0.00000000000007525744255
6  15:62520456:t:g -0.006162 0.0008061 0.00000000000002095786625
7  15:62676645:c:t -0.006109 0.0008107 0.00000000000004843344437
8  15:62768090:t:c -0.006310 0.0008021 0.00000000000000364390023
9  15:62979292:g:c -0.006268 0.0008077 0.00000000000000849284729
10 15:63141567:g:a -0.001523 0.0008172 0.06237707688136712169680
11 15:63353203:c:a -0.006145 0.0008235 0.00000000000008518687468
12 15:63863320:g:a -0.006494 0.0006947 0.00000000000000000000889
13 15:63902855:g:a -0.006055 0.0008304 0.00000000000030651940511
14             All -0.006062 0.0007953 0.00000000000002492497949
```

```
p2 = mr_scatter_plot(res, dat)[[1]] +
  xlab(expression(paste("SNP effect on M", Phi, " LACTB mRNA levels"))) +
  ylab("SNP effect on CSF succynilcarnitine levels")

p2
```

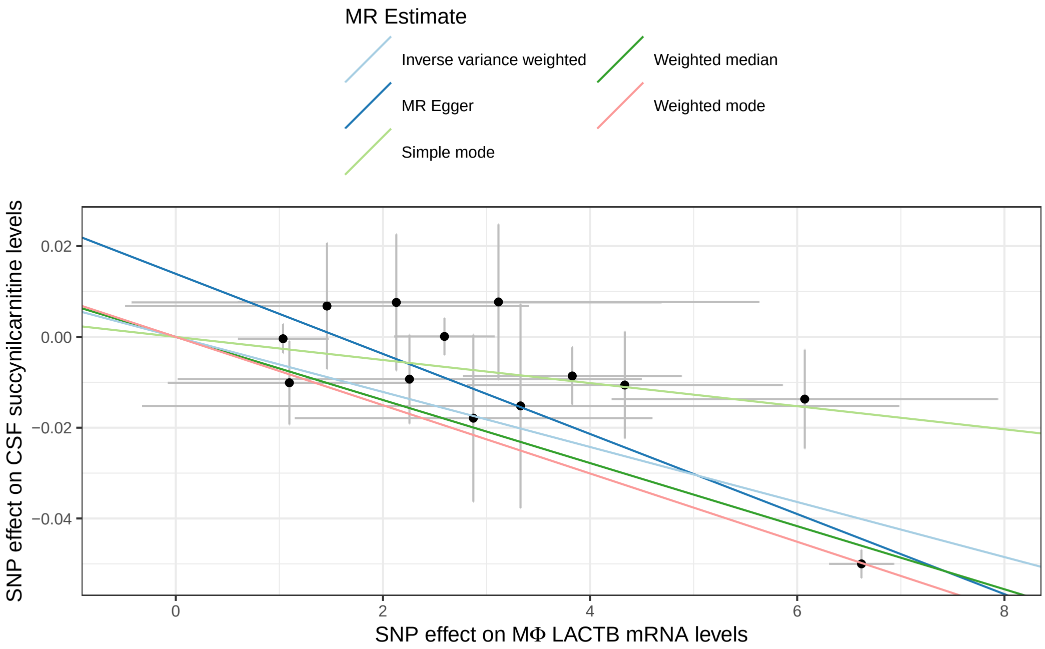


```
ggsave(str_interp("${res$exposure[1]}.${res$outcome[1]}.pdf"), plot = p2, width = 10.5, heig
```

```
dat = harmonise_data(exposure_dat = mqtl.gene.exposure,
                     outcome_dat = gwas.gene.outcome)
```

```
Harmonising CSF_X100001948 (JfjjnJ) and AD (5rt0Eu)

Removing the following SNPs for being palindromic with intermediate allele frequencies:
15:62148640:a:t
```

```
(res = mr(dat))
```

```
Analysing 'JfjjnJ' on '5rt0Eu'

  id.exposure id.outcome outcome       exposure                    method nsnp
1      JfjjnJ     5rt0Eu      AD CSF_X100001948                  MR Egger    8
2      JfjjnJ     5rt0Eu      AD CSF_X100001948           Weighted median    8
3      JfjjnJ     5rt0Eu      AD CSF_X100001948 Inverse variance weighted    8
4      JfjjnJ     5rt0Eu      AD CSF_X100001948               Simple mode    8
5      JfjjnJ     5rt0Eu      AD CSF_X100001948             Weighted mode    8
```

```
        b     se         pval
1 -0.8805 0.4005 0.070238964
2 -0.8684 0.1780 0.000001062
3 -0.8873 0.2028 0.000012128
4 -1.5549 0.6158 0.039505462
5 -0.8809 0.1803 0.001780223
```

```r
mr_heterogeneity(dat)
```

```
  id.exposure id.outcome outcome       exposure                    method    Q
1      JfjjnJ     5rtOEu      AD CSF_X100001948                  MR Egger 11.7
2      JfjjnJ     5rtOEu      AD CSF_X100001948 Inverse variance weighted 11.7
  Q_df  Q_pval
1    6 0.06902
2    7 0.11086
```

```r
mr_pleiotropy_test(dat)
```

```
  id.exposure id.outcome outcome       exposure egger_intercept      se   pval
1      JfjjnJ     5rtOEu      AD CSF_X100001948      -0.0002445 0.01199 0.9844
```

```r
mr_leaveoneout(dat)
```

```
        exposure outcome id.exposure id.outcome samplesize             SNP
1 CSF_X100001948      AD      JfjjnJ     5rtOEu     487511 15:62455571:c:t
2 CSF_X100001948      AD      JfjjnJ     5rtOEu     487511 15:62561062:c:t
3 CSF_X100001948      AD      JfjjnJ     5rtOEu     487511 15:62926987:t:c
4 CSF_X100001948      AD      JfjjnJ     5rtOEu     487511 15:62929201:c:t
5 CSF_X100001948      AD      JfjjnJ     5rtOEu     487511 15:62979292:g:c
6 CSF_X100001948      AD      JfjjnJ     5rtOEu     487511 15:63138257:c:t
7 CSF_X100001948      AD      JfjjnJ     5rtOEu     487511 15:63142564:a:g
8 CSF_X100001948      AD      JfjjnJ     5rtOEu     487511 15:63518986:g:c
9 CSF_X100001948      AD      JfjjnJ     5rtOEu     487511             All
        b     se           p
1 -0.9288 0.1956 0.000002043
2 -0.9065 0.1960 0.000003737
3 -0.8666 0.2073 0.000029091
4 -0.8857 0.2227 0.000069647
5 -0.8811 0.2168 0.000048266
6 -0.8044 0.1842 0.000012591
```

```
7 -0.9891 0.5258 0.059959714
8 -0.9167 0.2076 0.000010031
9 -0.8873 0.2028 0.000012128
```

```
p3 = mr_scatter_plot(res, dat)[[1]] +
  xlab("SNP effect on CSF succynilcarnitine levels") +
  ylab("SNP effect on AD risk")

p3
```

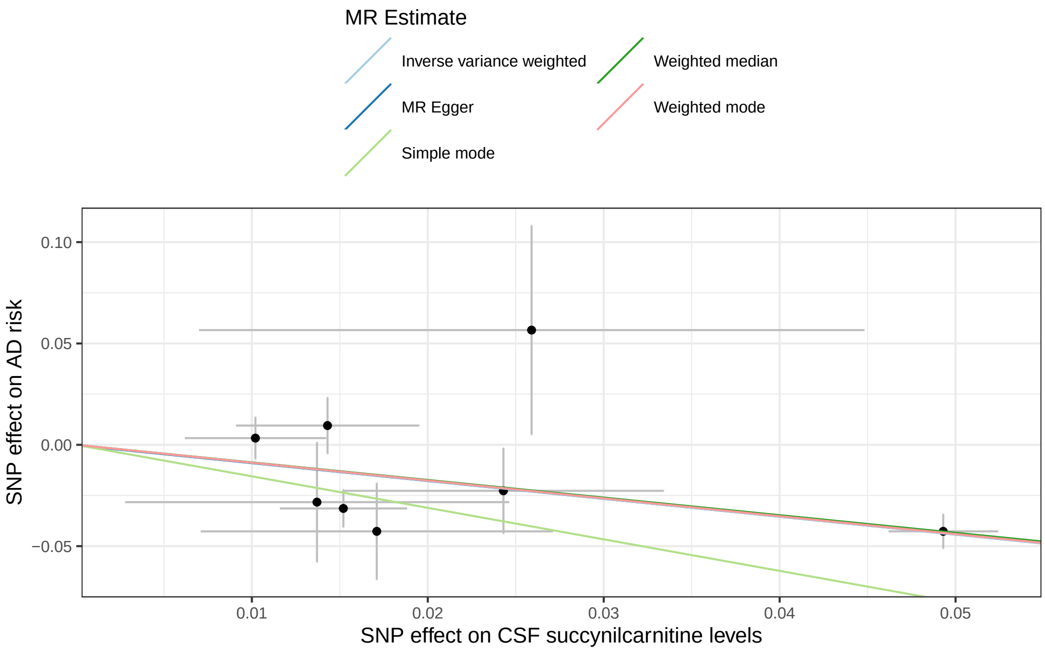


```
ggsave(str_interp("${res$exposure[1]}.${res$outcome[1]}.pdf"), plot = p3, width = 10.5, heig
```

```
p <- p1 + p2 + p3 + plot_layout(guides = "collect") & theme(legend.position = 'top')

p
```

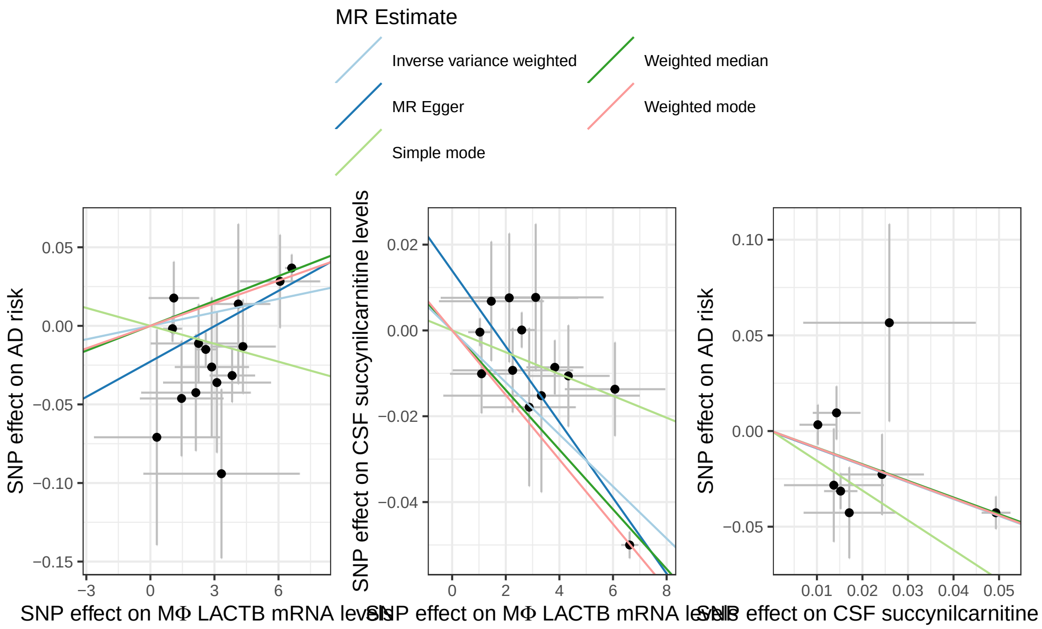


```
ggsave("lactb_succinylcarnitine_ad.mr_plot.pdf", plot = p, width = 7.5, height = 4.5, units =
```

##### Locuszoom plots

```
gwas.loc <- locus(data = gwas, gene = "LACTB", flank = 5e5, ens_db = EnsDb.Hsapiens.v75, chro
```

```
rs2729835, LACTB, chromosome 15, position 62913999 to 63934260

8147 SNPs/datapoints
```

```
summary(gwas.loc)
```

```
Gene LACTB
Chromosome 15, position 62,913,999 to 63,934,260
8147 SNPs/datapoints
23 gene transcripts
11 protein_coding, 5 antisense, 4 lincRNA, 2 miRNA, 1 pseudogene
Ensembl version: 75
Organism: Homo sapiens
Genome build: GRCh37
```

```r
gwas.loc <- link_LD(gwas.loc, token = "8656d0cbaeac")
```

```
LDlink server is working...

Matched 1219 SNPs (6.87 secs)
```

```r
gwas.loc <- link_recomb(gwas.loc)
```

```
Retrieving recombination data from UCSC
```

```r
locus_plot(gwas.loc, highlight = "LACTB", filter_gene_biotype = 'protein_coding')
```

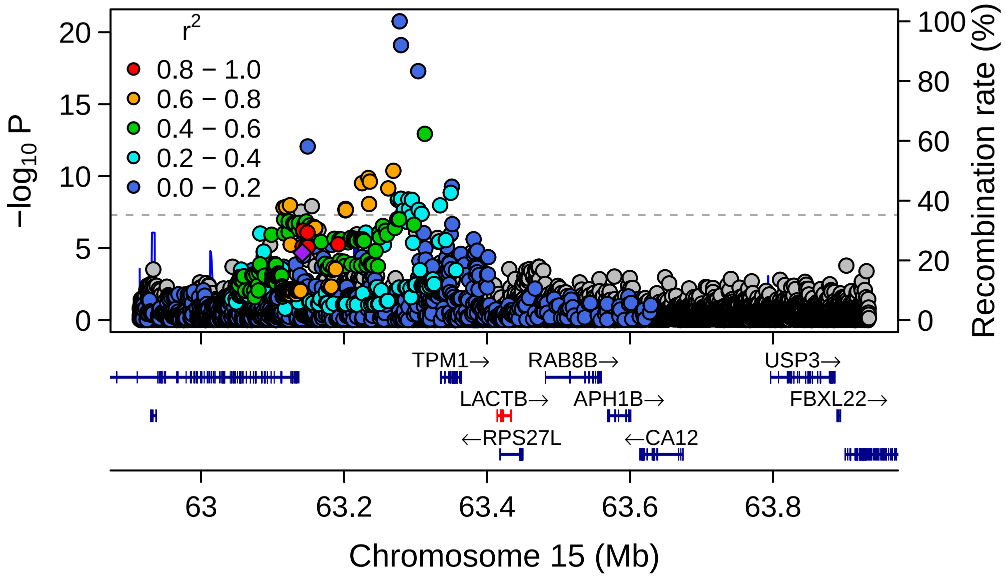


```r
eqtl.gene.loc <- locus(data = eqtl.gene, gene = "LACTB", flank = 5e5, ens_db = EnsDb.Hsapiens
```

```
rs2729835, LACTB, chromosome 15, position 62913999 to 63934260

3355 SNPs/datapoints
```

```r
summary(eqtl.gene.loc)
```

```
Gene LACTB
Chromosome 15, position 62,913,999 to 63,934,260
3355 SNPs/datapoints
23 gene transcripts
11 protein_coding, 5 antisense, 4 lincRNA, 2 miRNA, 1 pseudogene
Ensembl version: 75
Organism: Homo sapiens
Genome build: GRCh37
```

```r
eqtl.gene.loc <- link_LD(eqtl.gene.loc, token = "8656d0cbaeac")
```

```
Matched 1087 SNPs (0.000451 secs)
```

```r
eqtl.gene.loc <- link_recomb(eqtl.gene.loc)

locus_plot(eqtl.gene.loc, highlight = "LACTB", filter_gene_biotype = 'protein_coding')
```

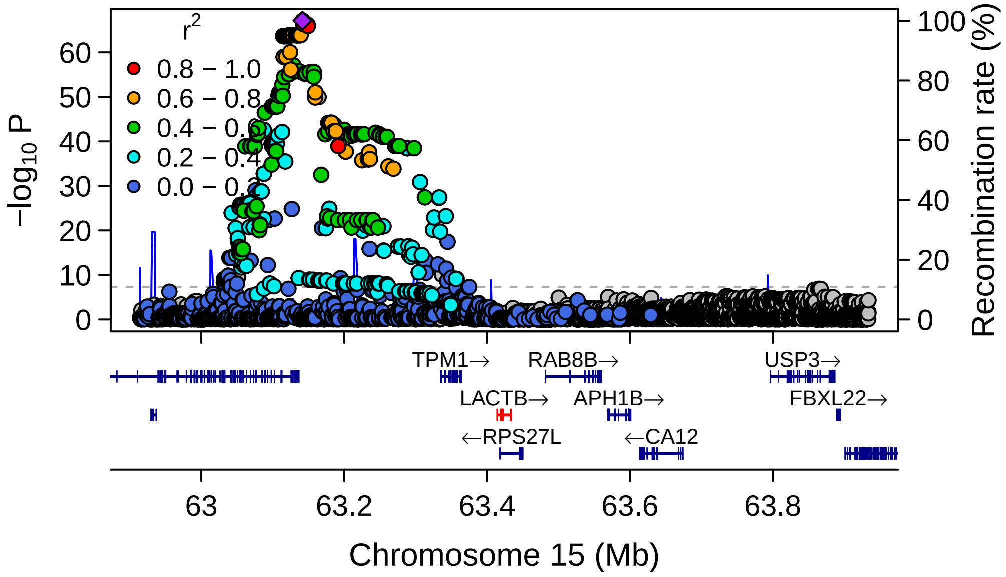

```r
mqtl.loc <- locus(data = eqtl, gene = "LACTB", flank = 5e5, ens_db = EnsDb.Hsapiens.v75, chro
```

```
rs2729835, LACTB, chromosome 15, position 62913999 to 63934260
```

```
76622 SNPs/datapoints
```

```r
summary(mqtl.loc)
```

```
Gene LACTB
Chromosome 15, position 62,913,999 to 63,934,260
76622 SNPs/datapoints
23 gene transcripts
11 protein_coding, 5 antisense, 4 lincRNA, 2 miRNA, 1 pseudogene
Ensembl version: 75
Organism: Homo sapiens
Genome build: GRCh37
```

```r
mqtl.loc <- link_LD(mqtl.loc, token = "8656d0cbaeac")
```

```
Matched 20563 SNPs (0.00403 secs)
```

```r
mqtl.loc <- link_recomb(mqtl.loc)

locus_plot(mqtl.loc, highlight = "LACTB", filter_gene_biotype = 'protein_coding')
```

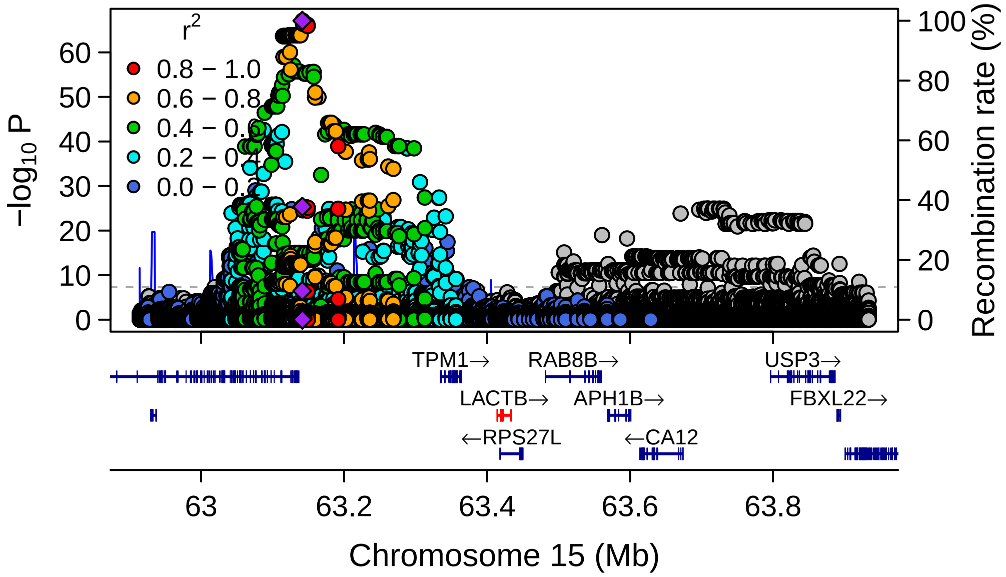


#### APH1B

```
GENE = "ENSG00000138613" # APH1B

eqtl.gene = eqtl %>% filter(TRAIT == GENE)

mqtl.gene = mqtl %>% filter(ID %in% eqtl.gene$ID)
gwas.gene = gwas %>% filter(ID %in% eqtl.gene$ID)
```

##### Format and harmonise data for MR

```
eqtl.gene.ld_clump = ieugwasr::ld_clump(eqtl.gene %>% select(rsid = ID, pval = P), bfile = "d
```

```
Clumping dHBOLF, 6472 variants, using: data/ldref/1000G.EUR.CPRA_b38.chr15
```

```
Removing 6453 of 6472 variants due to LD with other variants or absence from LD reference pa
```

```
eqtl.gene.exposure = format_data(eqtl.gene %>%
                                 filter(ID %in% eqtl.gene.ld_clump$rsid) %>%
                                 select(SNP = ID,
                                        beta = BETA,
                                        se = SE,
                                        effect_allele = ALT,
                                        other_allele = REF,
                                        eaf = AF,
                                        Phenotype = TRAIT,
                                        chr = CHROM,
                                        position = POS,
                                        sample_size = N,
                                        pval = P,
                                        gene = TRAIT),
                                 type="exposure")
```

```
mqtl.gene.ld_clump = ieugwasr::ld_clump(mqtl.gene %>% select(rsid = ID, pval = P), bfile = "[illegible]
```

```
Clumping uLTF71, 6097 variants, using: data/ldref/1000G.EUR.CPRA_b38.chr15
```

```
Removing 6087 of 6097 variants due to LD with other variants or absence from LD reference pa[illegible]
```

```
mqtl.gene.exposure = format_data(mqtl.gene %>%
                                 filter(ID %in% mqtl.gene.ld_clump$rsid) %>%
                                 select(SNP = ID,
                                        beta = BETA,
                                        se = SE,
                                        effect_allele = ALT,
                                        other_allele = REF,
                                        eaf = AF,
                                        Phenotype = TRAIT,
                                        chr = CHROM,
                                        position = POS,
                                        sample_size = N,
                                        pval = P),
                                 type="exposure")
```

```
mqtl.gene.outcome = format_data(mqtl.gene %>%
                                select(SNP = ID,
                                       beta = BETA,
```

```
                                                        se = SE,
                                                        effect_allele = ALT,
                                                        other_allele = REF,
                                                        eaf = AF,
                                                        Phenotype = TRAIT,
                                                        chr = CHROM,
                                                        position = POS,
                                                        sample_size = N,
                                                        pval = P),
                                                type="outcome")
```

```
gwas.gene.outcome = format_data(gwas.gene %>%
                                select(SNP = ID,
                                       beta = BETA,
                                       se = SE,
                                       effect_allele = ALT,
                                       other_allele = REF,
                                       eaf = AF,
                                       Phenotype = TRAIT,
                                       chr = CHROM,
                                       position = POS,
                                       sample_size = N,
                                       ncase = N_CASES,
                                       ncontrol = N_CTRLS,
                                       pval = P,
                                       gene = TRAIT),
                                type="outcome")
```

```
Generating sample size from ncase and ncontrol
```

```
dat = harmonise_data(exposure_dat = eqtl.gene.exposure,
                     outcome_dat = gwas.gene.outcome)
```

```
Harmonising ENSG00000138613 (mLJtA1) and AD (mwJnAA)
```

###### Analyse data and print/plot results

```
(res = mr(dat))
```

```
Analysing 'mLJtA1' on 'mwJnAA'
```

```
  id.exposure id.outcome outcome        exposure                    method nsnp
1      mLJtA1     mwJnAA      AD ENSG00000138613                  MR Egger   19
2      mLJtA1     mwJnAA      AD ENSG00000138613           Weighted median   19
3      mLJtA1     mwJnAA      AD ENSG00000138613 Inverse variance weighted   19
4      mLJtA1     mwJnAA      AD ENSG00000138613               Simple mode   19
5      mLJtA1     mwJnAA      AD ENSG00000138613             Weighted mode   19
            b       se   pval
1 -0.02877396 0.017004 0.1089
2 -0.00758644 0.010150 0.4548
3  0.00002654 0.007999 0.9974
4 -0.01693756 0.014170 0.2475
5 -0.00909930 0.008696 0.3093
```

```
mr_heterogeneity(dat)
```

```
  id.exposure id.outcome outcome        exposure                    method
1      mLJtA1     mwJnAA      AD ENSG00000138613                  MR Egger
2      mLJtA1     mwJnAA      AD ENSG00000138613 Inverse variance weighted
      Q Q_df Q_pval
1 18.62   17 0.3505
2 22.52   18 0.2096
```

```
mr_pleiotropy_test(dat)
```

```
  id.exposure id.outcome outcome        exposure egger_intercept      se
1      mLJtA1     mwJnAA      AD ENSG00000138613         0.03138 0.01664
     pval
1 0.07646
```

```
mr_leaveoneout(dat)
```

```
         exposure outcome id.exposure id.outcome samplesize             SNP
1 ENSG00000138613      AD      mLJtA1     mwJnAA     487511 15:62311651:t:c
2 ENSG00000138613      AD      mLJtA1     mwJnAA     487511 15:62456877:c:t
3 ENSG00000138613      AD      mLJtA1     mwJnAA     487511 15:62592045:t:c
4 ENSG00000138613      AD      mLJtA1     mwJnAA     487511 15:62595386:c:t
5 ENSG00000138613      AD      mLJtA1     mwJnAA     487511 15:62687961:a:g
6 ENSG00000138613      AD      mLJtA1     mwJnAA     487511 15:62780067:c:t
```

```
7  ENSG00000138613      AD      mLJtA1     mwJnAA     487511 15:62794527:c:t
8  ENSG00000138613      AD      mLJtA1     mwJnAA     487511 15:62822417:c:t
9  ENSG00000138613      AD      mLJtA1     mwJnAA     487511 15:62979292:g:c
10 ENSG00000138613      AD      mLJtA1     mwJnAA     487511 15:63164189:c:t
11 ENSG00000138613      AD      mLJtA1     mwJnAA     487511 15:63203962:c:t
12 ENSG00000138613      AD      mLJtA1     mwJnAA     487511 15:63273445:t:a
13 ENSG00000138613      AD      mLJtA1     mwJnAA     487511 15:63414875:a:t
14 ENSG00000138613      AD      mLJtA1     mwJnAA     487511 15:63564573:g:t
15 ENSG00000138613      AD      mLJtA1     mwJnAA     487511 15:63901140:g:a
16 ENSG00000138613      AD      mLJtA1     mwJnAA     487511 15:64008168:c:t
17 ENSG00000138613      AD      mLJtA1     mwJnAA     487511 15:64035151:g:a
18 ENSG00000138613      AD      mLJtA1     mwJnAA     487511 15:64053387:c:t
19 ENSG00000138613      AD      mLJtA1     mwJnAA     487511 15:64063833:g:a
20 ENSG00000138613      AD      mLJtA1     mwJnAA     487511              All
             b       se      p
1   0.00092932 0.008664 0.9146
2   0.00067406 0.008172 0.9343
3  -0.00035967 0.008015 0.9642
4   0.00005741 0.008270 0.9945
5  -0.00337940 0.007715 0.6613
6   0.00037103 0.008192 0.9639
7   0.00092112 0.008183 0.9104
8   0.00259991 0.008810 0.7679
9   0.00318750 0.008726 0.7149
10 -0.00131227 0.008771 0.8811
11  0.00060715 0.008209 0.9410
12 -0.00262957 0.007228 0.7160
13  0.00193277 0.009253 0.8345
14  0.00085777 0.008235 0.9170
15 -0.00107957 0.008186 0.8951
16  0.00069680 0.008058 0.9311
17 -0.00019107 0.008273 0.9816
18 -0.00020376 0.008223 0.9802
19 -0.00222237 0.007412 0.7643
20  0.00002654 0.007999 0.9974
```

```
p1 = mr_scatter_plot(res, dat)[[1]] +
  xlab(expression(paste("SNP effect on M", Phi, " APH1B mRNA levels"))) +
  ylab("SNP effect on AD risk")

p1
```

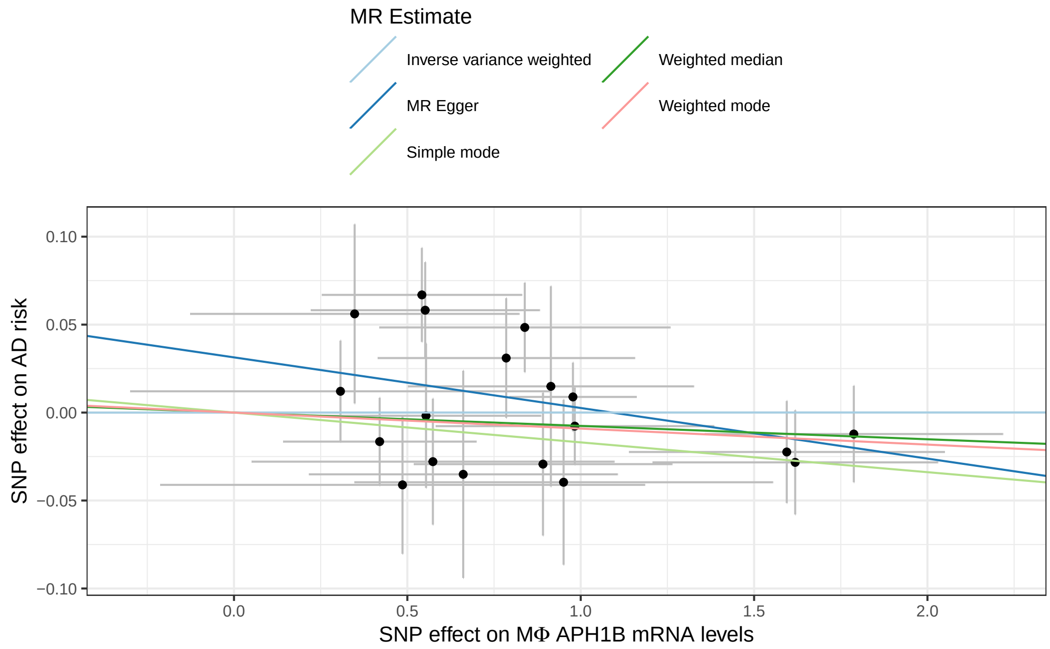


```
ggsave(str_interp("${res$exposure[1]}.${res$outcome[1]}.pdf"), plot = p1, width = 10.5, heig
```

```
dat = harmonise_data(exposure_dat = eqtl.gene.exposure,
                     outcome_dat = mqtl.gene.outcome)
```

```
Harmonising ENSG00000138613 (mLJtA1) and CSF_X100001948 (UjcIe9)
```

```
(res = mr(dat))
```

```
Analysing 'mLJtA1' on 'UjcIe9'
```

```
  id.exposure id.outcome        outcome        exposure
1      mLJtA1     UjcIe9 CSF_X100001948 ENSG00000138613
2      mLJtA1     UjcIe9 CSF_X100001948 ENSG00000138613
3      mLJtA1     UjcIe9 CSF_X100001948 ENSG00000138613
4      mLJtA1     UjcIe9 CSF_X100001948 ENSG00000138613
5      mLJtA1     UjcIe9 CSF_X100001948 ENSG00000138613
                     method nsnp        b       se    pval
1                  MR Egger   19 0.004992 0.008471 0.56338
2           Weighted median   19 0.003132 0.004295 0.46588
3 Inverse variance weighted   19 0.005930 0.003604 0.09988
```

```
4              Simple mode   19 0.005543 0.006263 0.38783
5            Weighted mode   19 0.005543 0.003923 0.17473
```

```r
mr_heterogeneity(dat)
```

```
  id.exposure id.outcome        outcome        exposure
1      mLJtA1     UjcIe9 CSF_X100001948 ENSG00000138613
2      mLJtA1     UjcIe9 CSF_X100001948 ENSG00000138613
                   method     Q Q_df  Q_pval
1                MR Egger 28.75   17 0.03689
2 Inverse variance weighted 28.78   18 0.05118
```

```r
mr_pleiotropy_test(dat)
```

```
  id.exposure id.outcome        outcome        exposure egger_intercept
1      mLJtA1     UjcIe9 CSF_X100001948 ENSG00000138613        0.001022
        se   pval
1 0.008304 0.9035
```

```r
mr_leaveoneout(dat)
```

```
          exposure        outcome id.exposure id.outcome samplesize
1  ENSG00000138613 CSF_X100001948      mLJtA1     UjcIe9         NA
2  ENSG00000138613 CSF_X100001948      mLJtA1     UjcIe9         NA
3  ENSG00000138613 CSF_X100001948      mLJtA1     UjcIe9         NA
4  ENSG00000138613 CSF_X100001948      mLJtA1     UjcIe9         NA
5  ENSG00000138613 CSF_X100001948      mLJtA1     UjcIe9         NA
6  ENSG00000138613 CSF_X100001948      mLJtA1     UjcIe9         NA
7  ENSG00000138613 CSF_X100001948      mLJtA1     UjcIe9         NA
8  ENSG00000138613 CSF_X100001948      mLJtA1     UjcIe9         NA
9  ENSG00000138613 CSF_X100001948      mLJtA1     UjcIe9         NA
10 ENSG00000138613 CSF_X100001948      mLJtA1     UjcIe9         NA
11 ENSG00000138613 CSF_X100001948      mLJtA1     UjcIe9         NA
12 ENSG00000138613 CSF_X100001948      mLJtA1     UjcIe9         NA
13 ENSG00000138613 CSF_X100001948      mLJtA1     UjcIe9         NA
14 ENSG00000138613 CSF_X100001948      mLJtA1     UjcIe9         NA
15 ENSG00000138613 CSF_X100001948      mLJtA1     UjcIe9         NA
16 ENSG00000138613 CSF_X100001948      mLJtA1     UjcIe9         NA
17 ENSG00000138613 CSF_X100001948      mLJtA1     UjcIe9         NA
18 ENSG00000138613 CSF_X100001948      mLJtA1     UjcIe9         NA
```

```
19 ENSG00000138613 CSF_X100001948      mLJtA1     UjcIe9          NA
20 ENSG00000138613 CSF_X100001948      mLJtA1     UjcIe9          NA
               SNP        b       se       p
1  15:62311651:t:c 0.004681 0.003819 0.22027
2  15:62456877:c:t 0.005974 0.003732 0.10941
3  15:62592045:t:c 0.005650 0.003481 0.10458
4  15:62595386:c:t 0.006286 0.003631 0.08344
5  15:62687961:a:g 0.006139 0.003838 0.10970
6  15:62780067:c:t 0.006064 0.003676 0.09905
7  15:62794527:c:t 0.005448 0.003579 0.12789
8  15:62822417:c:t 0.007435 0.003805 0.05067
9  15:62979292:g:c 0.005377 0.004081 0.18762
10 15:63164189:c:t 0.004126 0.003790 0.27625
11 15:63203962:c:t 0.005299 0.003478 0.12758
12 15:63273445:t:a 0.007230 0.003121 0.02053
13 15:63414875:a:t 0.006858 0.004178 0.10071
14 15:63564573:g:t 0.005814 0.003745 0.12048
15 15:63901140:g:a 0.005844 0.003755 0.11966
16 15:64008168:c:t 0.006252 0.003608 0.08309
17 15:64035151:g:a 0.006208 0.003691 0.09262
18 15:64053387:c:t 0.005942 0.003719 0.11011
19 15:64063833:g:a 0.005990 0.003741 0.10937
20             All 0.005930 0.003604 0.09988
```

```r
p2 = mr_scatter_plot(res, dat)[[1]] +
  xlab(expression(paste("SNP effect on M", Phi, " APH1B mRNA levels"))) +
  ylab("SNP effect on CSF succynilcarnitine levels")

p2
```

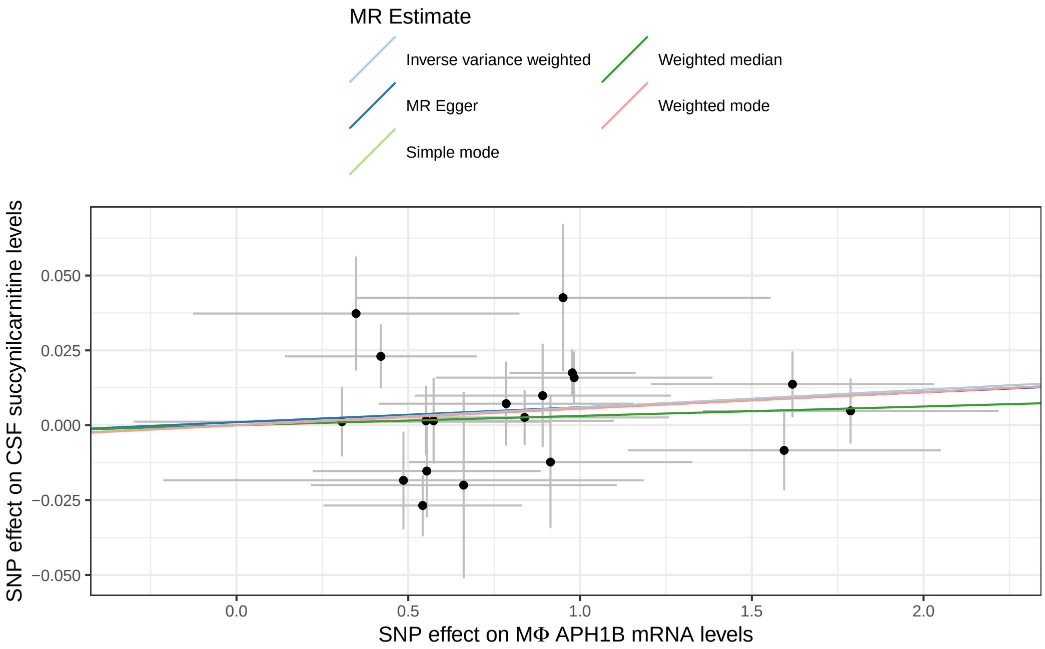


```
ggsave(str_interp("${res$exposure[1]}.${res$outcome[1]}.pdf"), plot = p2, width = 10.5, heig
```

```
dat = harmonise_data(exposure_dat = mqtl.gene.exposure,
                     outcome_dat = gwas.gene.outcome)
```

```
Harmonising CSF_X100001948 (1H2bXQ) and AD (mwJnAA)
```

```
(res = mr(dat))
```

```
Analysing '1H2bXQ' on 'mwJnAA'
```

```
  id.exposure id.outcome outcome      exposure                    method nsnp
1      1H2bXQ     mwJnAA      AD CSF_X100001948                  MR Egger   10
2      1H2bXQ     mwJnAA      AD CSF_X100001948           Weighted median   10
3      1H2bXQ     mwJnAA      AD CSF_X100001948 Inverse variance weighted   10
4      1H2bXQ     mwJnAA      AD CSF_X100001948               Simple mode   10
5      1H2bXQ     mwJnAA      AD CSF_X100001948             Weighted mode   10
        b     se        pval
1 -0.8343 0.3744 0.056419510
2 -0.8685 0.1809 0.000001574
3 -0.9014 0.1946 0.000003615
```

```
4 -1.8114 0.5942 0.013824366
5 -0.8596 0.1670 0.000605754
```

```
mr_heterogeneity(dat)
```

```
  id.exposure id.outcome outcome      exposure                    method     Q
1      1H2bXQ     mwJnAA      AD CSF_X100001948                  MR Egger 13.83
2      1H2bXQ     mwJnAA      AD CSF_X100001948 Inverse variance weighted 13.91
  Q_df  Q_pval
1    8 0.08623
2    9 0.12547
```

```
mr_pleiotropy_test(dat)
```

```
  id.exposure id.outcome outcome      exposure egger_intercept      se   pval
1      1H2bXQ     mwJnAA      AD CSF_X100001948       -0.002385 0.01111 0.8355
```

```
mr_leaveoneout(dat)
```

```
         exposure outcome id.exposure id.outcome samplesize              SNP
1  CSF_X100001948      AD      1H2bXQ     mwJnAA     483897 15:62390324:c:t
2  CSF_X100001948      AD      1H2bXQ     mwJnAA     483897 15:62455571:c:t
3  CSF_X100001948      AD      1H2bXQ     mwJnAA     483897 15:62561062:c:t
4  CSF_X100001948      AD      1H2bXQ     mwJnAA     483897 15:62926987:t:c
5  CSF_X100001948      AD      1H2bXQ     mwJnAA     483897 15:62929201:c:t
6  CSF_X100001948      AD      1H2bXQ     mwJnAA     483897 15:62979292:g:c
7  CSF_X100001948      AD      1H2bXQ     mwJnAA     483897 15:63138257:c:t
8  CSF_X100001948      AD      1H2bXQ     mwJnAA     483897 15:63142564:a:g
9  CSF_X100001948      AD      1H2bXQ     mwJnAA     483897 15:63518986:g:c
10 CSF_X100001948      AD      1H2bXQ     mwJnAA     483897 15:64050396:c:t
11 CSF_X100001948      AD      1H2bXQ     mwJnAA     483897              All
         b     se            p
1  -0.8940 0.2020 0.0000095852
2  -0.9430 0.1880 0.0000005294
3  -0.9205 0.1882 0.0000010014
4  -0.8809 0.1975 0.0000081903
5  -0.9003 0.2098 0.0000177930
6  -0.8952 0.2047 0.0000122868
7  -0.8199 0.1813 0.0000061441
8  -1.0663 0.4876 0.0287649500
```

```
9  -0.9309 0.1975 0.0000024208
10 -0.8947 0.1945 0.0000042094
11 -0.9014 0.1946 0.0000036145
```

```
p3 = mr_scatter_plot(res, dat)[[1]] +
  xlab("SNP effect on CSF succynilcarnitine levels") +
  ylab("SNP effect on AD risk")

p3
```

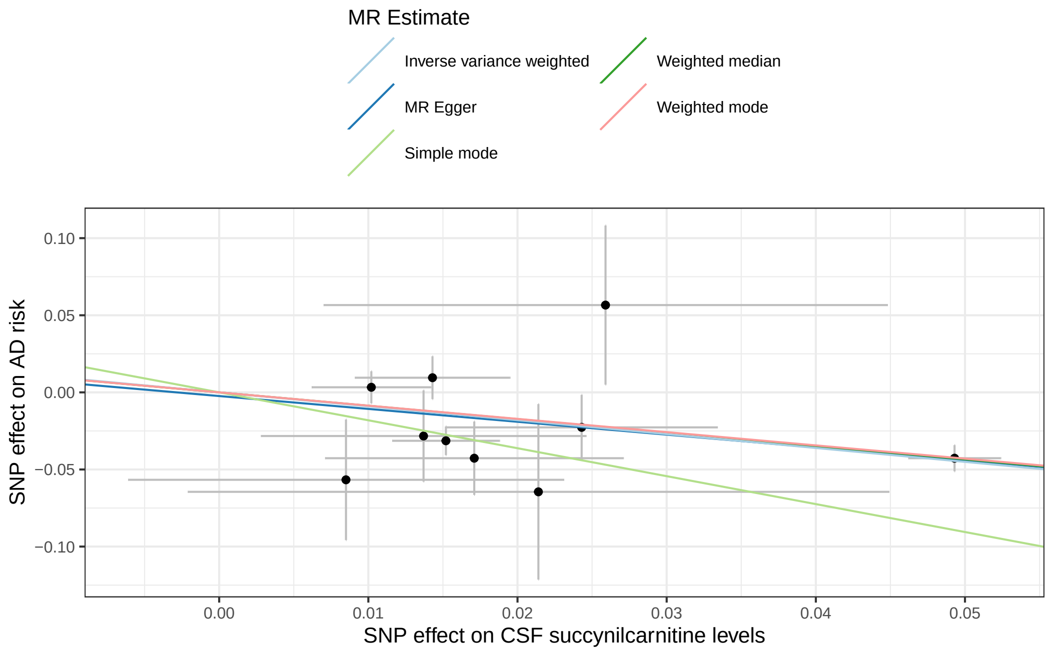


```
ggsave(str_interp("${res$exposure[1]}.${res$outcome[1]}.pdf"), plot = p3, width = 10.5, heigh
```

```
p <- p1 + p2 + p3 + plot_layout(guides = "collect") & theme(legend.position = 'top')

p
```

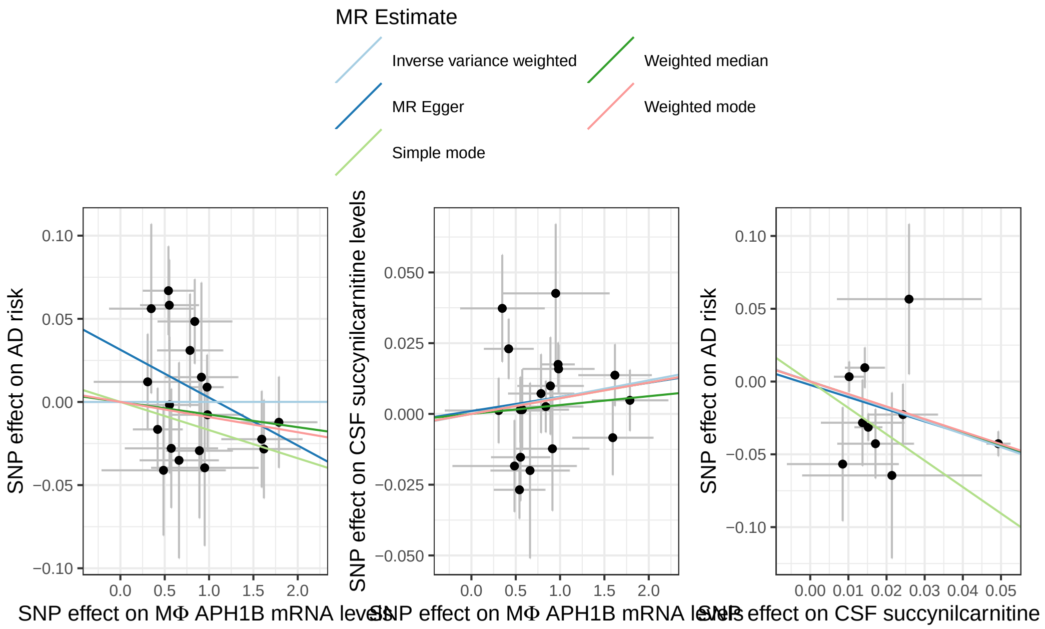


```r
ggsave("aph1b_succinylcarnitine_ad.mr_plot.pdf", plot = p, width = 7.5, height = 4.5, units =
```

##### Locuszoom plots

```r
gwas.loc <- locus(data = gwas, gene = "APH1B", flank = 5e5, ens_db = EnsDb.Hsapiens.v75, chro
```

```
rs2729835, APH1B, chromosome 15, position 63068217 to 64101325
```

```
8020 SNPs/datapoints
```

```r
summary(gwas.loc)
```

```
Gene APH1B
Chromosome 15, position 63,068,217 to 64,101,325
8020 SNPs/datapoints
22 gene transcripts
10 protein_coding, 5 antisense, 4 lincRNA, 2 miRNA, 1 pseudogene
Ensembl version: 75
Organism: Homo sapiens
Genome build: GRCh37
```

```r
gwas.loc <- link_LD(gwas.loc, token = "8656d0cbaeac")
```

```
Matched 848 SNPs (0.000954 secs)
```

```r
gwas.loc <- link_recomb(gwas.loc)
```

```
Retrieving recombination data from UCSC
```

```r
locus_plot(gwas.loc, highlight = "APH1B", filter_gene_biotype = 'protein_coding')
```

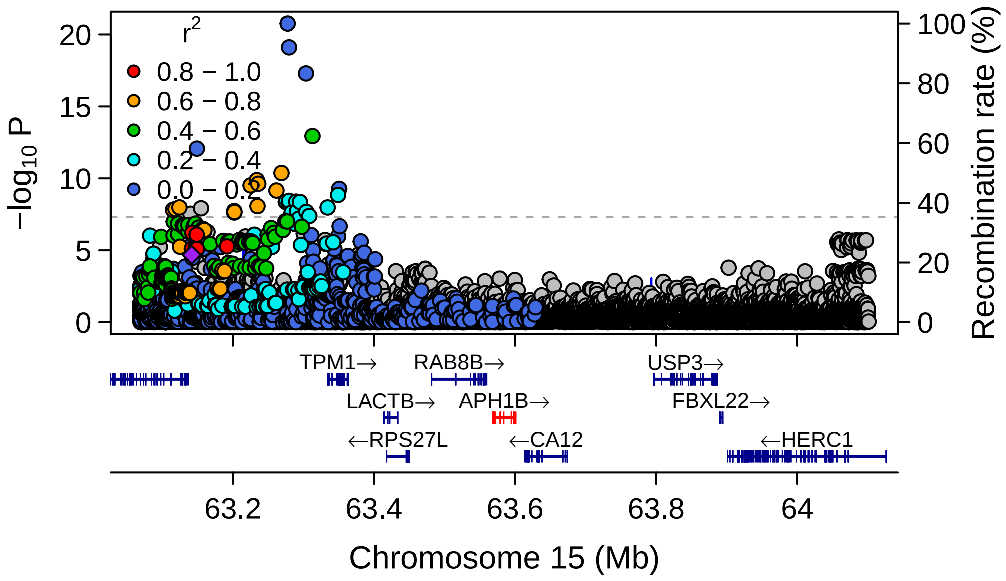


```r
eqtl.gene.loc <- locus(data = eqtl.gene, gene = "APH1B", flank = 5e5, ens_db = EnsDb.Hsapiens
```

```
rs2729835, APH1B, chromosome 15, position 63068217 to 64101325
```

```
3278 SNPs/datapoints
```

```r
summary(eqtl.gene.loc)
```

```
Gene APH1B
Chromosome 15, position 63,068,217 to 64,101,325
3278 SNPs/datapoints
22 gene transcripts
10 protein_coding, 5 antisense, 4 lincRNA, 2 miRNA, 1 pseudogene
Ensembl version: 75
Organism: Homo sapiens
Genome build: GRCh37
```

```
eqtl.gene.loc <- link_LD(eqtl.gene.loc, token = "8656d0cbaeac")
```

```
Matched 770 SNPs (0.000434 secs)
```

```
eqtl.gene.loc <- link_recomb(eqtl.gene.loc)

locus_plot(eqtl.gene.loc, highlight = "APH1B", filter_gene_biotype = 'protein_coding')
```

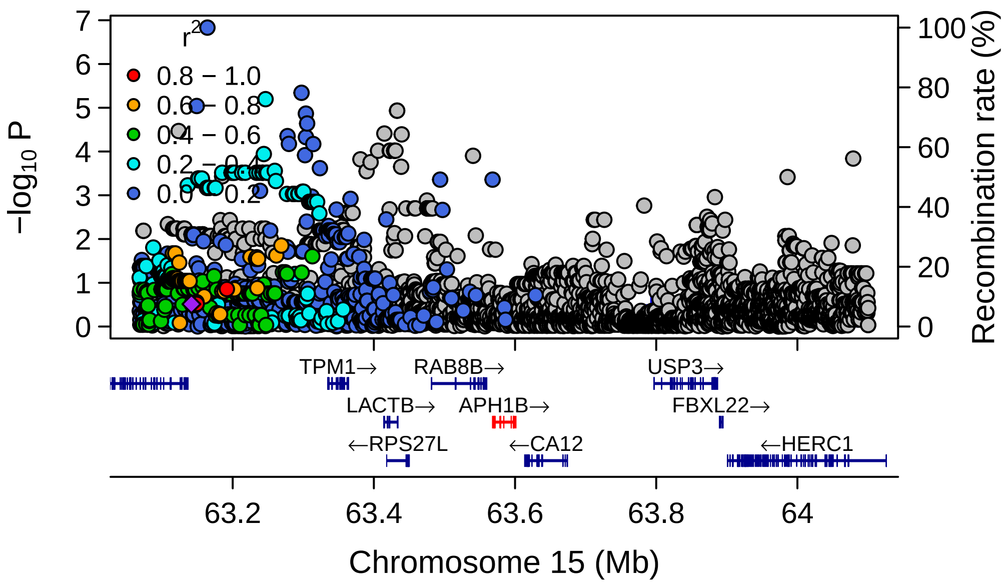


```
mqtl.loc <- locus(data = eqtl, gene = "APH1B", flank = 5e5, ens_db = EnsDb.Hsapiens.v75, chr
```

```
rs2729835, APH1B, chromosome 15, position 63068217 to 64101325
```

```
85570 SNPs/datapoints
```

```r
summary(mqtl.loc)
```

```
Gene APH1B
Chromosome 15, position 63,068,217 to 64,101,325
85570 SNPs/datapoints
22 gene transcripts
10 protein_coding, 5 antisense, 4 lincRNA, 2 miRNA, 1 pseudogene
Ensembl version: 75
Organism: Homo sapiens
Genome build: GRCh37
```

```r
mqtl.loc <- link_LD(mqtl.loc, token = "8656d0cbaeac")
```

```
Matched 15208 SNPs (0.00404 secs)
```

```r
mqtl.loc <- link_recomb(mqtl.loc)

locus_plot(mqtl.loc, highlight = "APH1B", filter_gene_biotype = 'protein_coding')
```

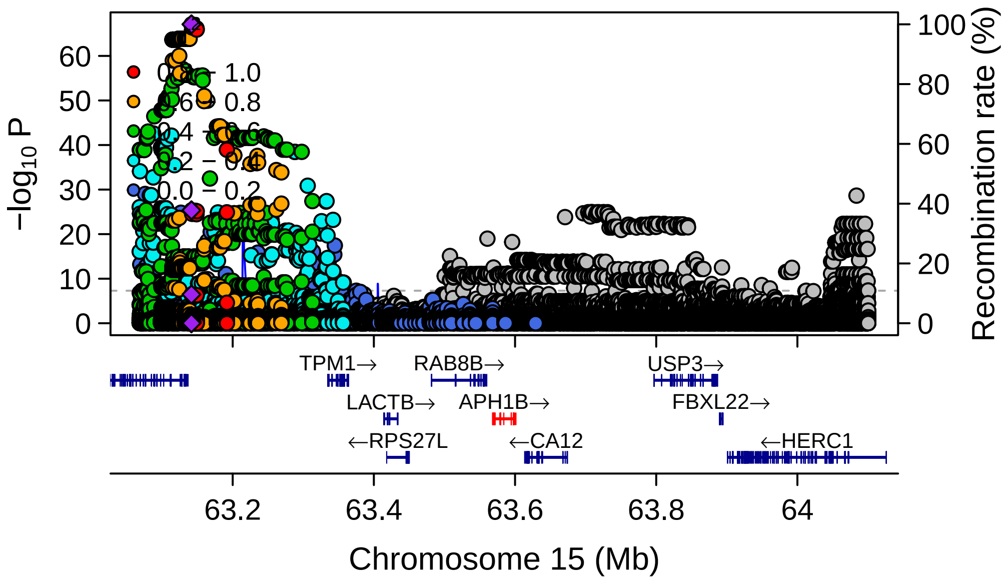

##### Print environment

```r
sessioninfo::session_info()
```

```
- Session info ---------------------------------------------------------------
 setting  value
 version  R version 4.5.2 (2025-10-31)
 os       macOS Sequoia 15.7.4
 system   aarch64, darwin20
 ui       X11
 language (EN)
 collate  en_US.UTF-8
 ctype    en_US.UTF-8
 tz       America/New_York
 date     2026-02-15
 pandoc   3.9 @ /opt/homebrew/bin/ (via rmarkdown)
 quarto   1.8.27 @ /usr/local/bin/quarto

- Packages -------------------------------------------------------------------
 package             * version   date (UTC) lib source
 abind                 1.4-8     2024-09-12 [1] CRAN (R 4.5.0)
 AnnotationDbi       * 1.72.0    2025-10-29 [1] Bioconductor 3.22 (R 4.5.1)
 AnnotationFilter    * 1.34.0    2025-10-29 [1] Bioconductor 3.22 (R 4.5.1)
 Biobase             * 2.70.0    2025-10-29 [1] Bioconductor 3.22 (R 4.5.1)
 BiocGenerics        * 0.56.0    2025-10-29 [1] Bioconductor 3.22 (R 4.5.1)
 BiocIO                1.20.0    2025-10-29 [1] Bioconductor 3.22 (R 4.5.1)
 BiocParallel          1.44.0    2025-10-29 [1] Bioconductor 3.22 (R 4.5.1)
 Biostrings            2.78.0    2025-10-29 [1] Bioconduc~
 bit                   4.6.0     2025-03-06 [1] CRAN (R 4.5.0)
 bit64                 4.6.0-1   2025-01-16 [1] CRAN (R 4.5.0)
 bitops                1.0-9     2024-10-03 [1] CRAN (R 4.5.0)
 blob                  1.3.0     2026-01-14 [1] CRAN (R 4.5.2)
 cachem                1.1.0     2024-05-16 [1] CRAN (R 4.5.0)
 cigarillo             1.0.0     2025-10-29 [1] Bioconduc~
 cli                   3.6.5     2025-04-23 [1] CRAN (R 4.5.0)
 codetools             0.2-20    2024-03-31 [2] CRAN (R 4.5.2)
 coro                  1.1.0     2024-11-05 [1] CRAN (R 4.5.0)
 cowplot               1.2.0     2025-07-07 [1] CRAN (R 4.5.0)
 crayon                1.5.3     2024-06-20 [1] CRAN (R 4.5.0)
 curl                  7.0.0     2025-08-19 [1] CRAN (R 4.5.0)
 data.table            1.18.2.1  2026-01-27 [1] CRAN (R 4.5.2)
```

```
DBI                    1.2.3    2024-06-02 [1] CRAN (R 4.5.0)
DelayedArray           0.36.0   2025-10-29 [1] Bioconductor 3.22 (R 4.5.1)
digest                 0.6.39   2025-11-19 [1] CRAN (R 4.5.2)
dplyr                * 1.2.0    2026-02-03 [1] CRAN (R 4.5.2)
ellmer                 0.4.0    2025-11-15 [1] CRAN (R 4.5.2)
EnsDb.Hsapiens.v75   * 2.99.0   2026-02-13 [1] Bioconductor
ensembldb            * 2.34.0   2025-10-29 [1] Bioconductor 3.22 (R 4.5.1)
evaluate               1.0.5    2025-08-27 [1] CRAN (R 4.5.0)
farver                 2.1.2    2024-05-13 [1] CRAN (R 4.5.0)
fastmap                1.2.0    2024-05-15 [1] CRAN (R 4.5.0)
forcats              * 1.0.1    2025-09-25 [1] CRAN (R 4.5.0)
generics             * 0.1.4    2025-05-09 [1] CRAN (R 4.5.0)
GenomeInfoDb           1.46.2   2025-12-08 [1] Bioconductor 3.22 (R 4.5.2)
GenomicAlignments      1.46.0   2025-10-29 [1] Bioconduc~
GenomicFeatures      * 1.62.0   2025-10-29 [1] Bioconduc~
GenomicRanges        * 1.62.1   2025-12-08 [1] Bioconductor 3.22 (R 4.5.2)
gggrid                 0.2-0    2022-01-11 [1] CRAN (R 4.5.0)
ggplot2              * 4.0.2    2026-02-03 [1] CRAN (R 4.5.2)
ggrepel                0.9.6    2024-09-07 [1] CRAN (R 4.5.0)
glue                   1.8.0    2024-09-30 [1] CRAN (R 4.5.0)
gtable                 0.3.6    2024-10-25 [1] CRAN (R 4.5.0)
hms                    1.1.4    2025-10-17 [1] CRAN (R 4.5.0)
htmltools              0.5.9    2025-12-04 [1] CRAN (R 4.5.2)
htmlwidgets            1.6.4    2023-12-06 [1] CRAN (R 4.5.0)
httr                   1.4.8    2026-02-13 [1] CRAN (R 4.5.2)
httr2                  1.2.2    2025-12-08 [1] CRAN (R 4.5.2)
ieugwasr               1.1.0    2025-07-31 [1] CRAN (R 4.5.0)
IRanges              * 2.44.0   2025-10-29 [1] Bioconduc~
jsonlite               2.0.0    2025-03-27 [1] CRAN (R 4.5.0)
KEGGREST               1.50.0   2025-10-29 [1] Bioconductor 3.22 (R 4.5.1)
knitr                  1.51     2025-12-20 [1] CRAN (R 4.5.2)
labeling               0.4.3    2023-08-29 [1] CRAN (R 4.5.0)
lattice                0.22-9   2026-02-09 [1] CRAN (R 4.5.2)
lazyeval               0.2.2    2019-03-15 [1] CRAN (R 4.5.0)
LDlinkR                1.4.0    2024-04-10 [1] CRAN (R 4.5.0)
lifecycle              1.0.5    2026-01-08 [1] CRAN (R 4.5.2)
locuszoomr           * 0.3.8    2025-03-03 [1] CRAN (R 4.5.0)
lubridate            * 1.9.5    2026-02-04 [1] CRAN (R 4.5.2)
magrittr             * 2.0.4    2025-09-12 [1] CRAN (R 4.5.0)
Matrix                 1.7-4    2025-08-28 [2] CRAN (R 4.5.2)
MatrixGenerics         1.22.0   2025-10-29 [1] Bioconductor 3.22 (R 4.5.1)
matrixStats            1.5.0    2025-01-07 [1] CRAN (R 4.5.0)
memoise                2.0.1    2021-11-26 [1] CRAN (R 4.5.0)
```

```
 otel                     0.2.0     2025-08-29 [1] CRAN (R 4.5.0)
 patchwork              * 1.3.2     2025-08-25 [1] CRAN (R 4.5.0)
 pillar                   1.11.1    2025-09-17 [1] CRAN (R 4.5.0)
 pkgconfig                2.0.3     2019-09-22 [1] CRAN (R 4.5.0)
 plotly                   4.12.0    2026-01-24 [1] CRAN (R 4.5.2)
 plyr                     1.8.9     2023-10-02 [1] CRAN (R 4.5.0)
 png                      0.1-8     2022-11-29 [1] CRAN (R 4.5.0)
 ProtGenerics             1.42.0    2025-10-29 [1] Bioconductor 3.22 (R 4.5.1)
 purrr                  * 1.2.1     2026-01-09 [1] CRAN (R 4.5.2)
 R6                       2.6.1     2025-02-15 [1] CRAN (R 4.5.0)
 ragg                     1.5.0     2025-09-02 [1] CRAN (R 4.5.0)
 rappdirs                 0.3.4     2026-01-17 [1] CRAN (R 4.5.2)
 RColorBrewer             1.1-3     2022-04-03 [1] CRAN (R 4.5.0)
 Rcpp                     1.1.1     2026-01-10 [1] CRAN (R 4.5.2)
 RCurl                    1.98-1.17 2025-03-22 [1] CRAN (R 4.5.0)
 readr                  * 2.1.6     2025-11-14 [1] CRAN (R 4.5.2)
 restfulr                 0.0.16    2025-06-27 [1] CRAN (R 4.5.0)
 rjson                    0.2.23    2024-09-16 [1] CRAN (R 4.5.0)
 rlang                    1.1.7     2026-01-09 [1] CRAN (R 4.5.2)
 rmarkdown                2.30      2025-09-28 [1] CRAN (R 4.5.0)
 Rsamtools                2.26.0    2025-10-29 [1] Bioconduc~
 RSQLite                  2.4.6     2026-02-06 [1] CRAN (R 4.5.2)
 rstudioapi               0.18.0    2026-01-16 [1] CRAN (R 4.5.2)
 rtracklayer              1.70.1    2025-12-20 [1] https://bioc-release.r-
universe.dev (R 4.5.2)
 S4Arrays                 1.10.1    2025-12-08 [1] Bioconductor 3.22 (R 4.5.2)
 S4Vectors              * 0.48.0    2025-10-29 [1] Bioconduc~
 S7                       0.2.1     2025-11-14 [1] CRAN (R 4.5.2)
 scales                   1.4.0     2025-04-24 [1] CRAN (R 4.5.0)
 Seqinfo                * 1.0.0     2025-10-29 [1] Bioconduc~
 sessioninfo              1.2.3     2025-02-05 [1] CRAN (R 4.5.0)
 SparseArray              1.10.8    2025-12-18 [1] Bioconductor 3.22 (R 4.5.2)
 stringi                  1.8.7     2025-03-27 [1] CRAN (R 4.5.0)
 stringr                * 1.6.0     2025-11-04 [1] CRAN (R 4.5.0)
 SummarizedExperiment     1.40.0    2025-10-29 [1] Bioconduc~
 systemfonts              1.3.1     2025-10-01 [1] CRAN (R 4.5.0)
 textshaping              1.0.4     2025-10-10 [1] CRAN (R 4.5.0)
 tibble                 * 3.3.1     2026-01-11 [1] CRAN (R 4.5.2)
 tidyr                  * 1.3.2     2025-12-19 [1] CRAN (R 4.5.2)
 tidyselect               1.2.1     2024-03-11 [1] CRAN (R 4.5.0)
 tidyverse              * 2.0.0     2023-02-22 [1] CRAN (R 4.5.0)
 timechange               0.4.0     2026-01-29 [1] CRAN (R 4.5.2)
 TwoSampleMR            * 0.6.22    2025-10-25 [1] https://mrcieu.r-universe.dev (R 4.5.1)
```

```
tzdb                   0.5.0      2025-03-15 [1] CRAN (R 4.5.0)
UCSC.utils             1.6.1      2025-12-11 [1] Bioconductor 3.22 (R 4.5.2)
vctrs                  0.7.1      2026-01-23 [1] CRAN (R 4.5.2)
viridisLite            0.4.3      2026-02-04 [1] CRAN (R 4.5.2)
withr                  3.0.2      2024-10-28 [1] CRAN (R 4.5.0)
xfun                   0.56       2026-01-18 [1] CRAN (R 4.5.2)
XML                    3.99-0.22 2026-02-10 [1] CRAN (R 4.5.2)
XVector                0.50.0     2025-10-29 [1] Bioconductor 3.22 (R 4.5.1)
yaml                   2.3.12     2025-12-10 [1] CRAN (R 4.5.2)
zoo                    1.8-15     2025-12-15 [1] CRAN (R 4.5.2)

[1] /Users/marcoe02/.Rlib
[2] /Library/Frameworks/R.framework/Versions/4.5-arm64/Resources/library
* -- Packages attached to the search path.

------------------------------------------------------------------------------
```
